# Growing Glycans in Rosetta: Accurate *de novo* glycan modeling, density fitting, and rational sequon design

**DOI:** 10.1101/2021.09.27.462000

**Authors:** Jared Adolf-Bryfogle, Jason W. Labonte, John C. Kraft, Maxim Shapovalov, Sebastian Raemisch, Thomas Lütteke, Frank DiMaio, Christopher D. Bahl, Jesper Pallesen, Neil P. King, Jeffrey J. Gray, Daniel W. Kulp, William R. Schief

## Abstract

Carbohydrates and glycoproteins modulate key biological functions. Computational approaches inform function to aid in carbohydrate structure prediction, structure determination, and design. However, experimental structure determination of sugar polymers is notoriously difficult as glycans can sample a wide range of low energy conformations, thus limiting the study of glycan-mediated molecular interactions. In this work, we expanded the *RosettaCarbohydrate* framework, developed and benchmarked effective tools for glycan modeling and design, and extended the Rosetta software suite to better aid in structural analysis and benchmarking tasks through the SimpleMetrics framework. We developed a glycan-modeling algorithm, *GlycanTreeModeler*, that computationally builds glycans layer-by-layer, using adaptive kernel density estimates (KDE) of common glycan conformations derived from data in the Protein Data Bank (PDB) and from quantum mechanics (QM) calculations. After a rigorous optimization of kinematic and energetic considerations to improve near-native sampling enrichment and decoy discrimination, *GlycanTreeModeler* was benchmarked on a test set of diverse glycan structures, or “trees”. Structures predicted by *GlycanTreeModeler* agreed with native structures at high accuracy for both *de novo* modeling and experimental density-guided building. *GlycanTreeModeler* algorithms and associated tools were employed to design *de novo* glycan trees into a protein nanoparticle vaccine that are able to direct the immune response by shielding regions of the scaffold from antibody recognition. This work will inform glycoprotein model prediction, aid in both X-ray and electron microscopy density solutions and refinement, and help lead the way towards a new era of computational glycobiology.

## Introduction

Carbohydrates and glycoproteins are ubiquitous in biological organisms^1^. Viral glycoproteins such as HIV envelope trimer, influenza hemagglutinin, and SARS-CoV-2 spike, employ N-linked glycosylation as an immune evasion strategy, taking advantage of the fact that host glycans on the surface of proteins are usually recognized as “self” by the adaptive immune system^2^. Yet, HIV broadly neutralizing antibodies often target glycans as part of their epitopes^3,4,5^. Small carbohydrate residues attached to serine or threonine can act in signaling pathways akin to phosphorylation^6^, while glycans on the constant region of antibodies act as mediators of effector function^7,8^. Glycans can also improve stability^9^ and solubility^10^, reduce aggregation^11^, and even improve biological drug-targeting and vaccine design through glycan masking of off-target regions^12^.

The biosynthesis of glycoconjugates is complex. Carbohydrates can be attached to certain amino acid residues including serine, threonine, asparagine, and (rarely) tryptophan through covalent modification, forming glycoproteins. The attachment can be made to nitrogen, oxygen, or carbon atoms, (known as N*-*, O*-*, or C-linked glycosylation, respectively), with each process involving a multitude of enzymes, sugar moieties and resulting carbohydrate structures. These processes are stochastic in nature, producing glycoproteins that are heterogeneous in both the occupancy of a glycan at the glycosylation site (macro-heterogenicity) and the chemical makeup of the N-, C-, or O-linked glycan (micro-heterogenicity)^13^.

The most common form of glycosylation observed in glycoprotein structures is N-linked glycosylation. Initiation of this process occurs during translation, by the protein oligosaccharyltransferase (OST), which recognizes a multi-residue consensus motif, or sequon, of NX(S/T) (where X is any residue except proline), and covalently attaches a lipid-linked core-oligosaccharide to the asparagine residue through an N-glycoside linkage ^1^. This process is not deterministic (not every sequon results in attachment of a glycan) and certain amino acids in and around the sequon motif can affect the efficiency of this process, resulting in higher or lower glycan occupancy at the site^14,15^.

Upon successful protein folding in the endoplasmic reticulum, the initial N-linked glycan is “trimmed down” by removal of several terminal glucosyl residues, while many sugar processing enzymes in the Golgi apparatus can add or remove sugar residues from the nascent branched sugar (tree). The resulting chemical makeup of the glycan tree depends on which enzymes are available in the Golgi, which is heavily influenced by species, disease state^16^, developmental stage^17^; and the local structure, sequence, and environment of the glycosylation site^18^. In addition, a particular glycosylation site can result in vastly different glycans^19^, though this can be controlled to some extent through various bioengineering techniques^13,20,21^.

Glycans are also conformationally flexible, being highly hydrophilic and typically exposed on the surface of proteins, with a large number of conformational degrees of freedom. However, as has been observed in molecular dynamics and NMR experiments, glycan conformations can be influenced by their structural environment^22^. Through the plethora of high-resolution crystallographic and cryo-EM studies, we also know that glycans can adopt stable conformations with well-defined density observed for many of the glycan residues in each tree, especially towards the root of the glycan tree, even for some unrestrained glycans^23,24^. Presumably, these low-energy, stable conformations are occupied at higher frequency in solution. In addition, a recent QM study on glycan torsional energies showed that the QM-derived conformational preferences of glycan torsions match well with glycan structures analyzed from the protein data bank, indicating that conformational diversity is also influenced by the chemical makeup of each glycan structure^25^.

Given the complex chemistry and conformational diversity involved, accurate modeling of glycans is currently a grand challenge in computational biology. Computational glycobiology tools and webapps have been developed for protein glycosylations^26^, validation of carbohydrate structural chemistry^27^, statistical analysis^28^, and docking^29,30^. Common methods in glycoprotein modeling typically involve molecular dynamics (MD) simulations^31^ or adding glycans by manual placement and conformational tweaking into their density for structure determination^32^. Recently, a new method for automatic building of glycan structures from sequence was described^33^; this method, the CHARM-GUI Glycan Modeler, was benchmarked only up to the first and second sugar.

Here we describe a new glycan modeling algorithm built within the Rosetta software suite, a platform that incorporates state-of-the-art applications and modules for a variety of macromolecular modeling and design tasks^34^. The new algorithm provides user interfaces for the creation of tailor-made protocols^35,36^ and includes a reliable knowledge-based energy function to evaluate models and designs^37^. We build on earlier work that enabled representing and evaluating carbohydrate structures within Rosetta^38^ and in loading, representing, and refining glycans from the Protein Data Bank^39^. We expand on this foundational work through the addition of new carbohydrate-specific sampling methods, an updated conformer database employing adaptive kernel density estimates, a new framework for general analysis in Rosetta (SimpleMetrics), and a new algorithm for accurately modeling complex carbohydrates, the *GlycanTreeModeler*.

We rigorously benchmark the new method on a set of diverse high-resolution crystal structures of glycans in symmetric crystal environments, and we show that the *GlycanTreeModeler* is capable of recapitulating native glycan structures with high accuracy both through *de novo* and density-guided modeling^40^. We then applied our glycan modeling protocol with Rosetta sequence design of glycan sequons to engineer optimal new glycans onto a protein nanoparticle vaccine scaffold and evaluated changes in immune responses. We observed reduced reactivity to the underlying protein surface in immunization experiments, thus demonstrating that glycans can be computationally engineered to tailor immunogenicity of vaccines.

## Results

The Rosetta *GlycanTreeModeler* builds whole glycan “trees” through an algorithm that mimics the growth of natural trees. A primary difficulty in *de novo* glycan modeling is the correct prediction of the base of glycoconjugate structures. To increase the accuracy of the first few sugars of the tree, our algorithm begins modeling from the “root” (reducing end) of the glycan tree out to the branching “foliage”. Monte Carlo optimization through sampling of glycan degrees of freedom (DOFs) is carried out through the new *GlycanSampler*, which includes routines for glycosidic torsion angle (backbone) sampling, structure minimization, hydroxyl and other side-chain optimization, and neighbor protein side-chain optimization. During the protocol, the total amount of sampling scales linearly with the number of glycan residues being modeled, ensuring even sampling regardless of the size or quantity of glycans being modeled.

The *GlycanSampler* optimizes glycosidic torsion angles using statistically favorable sets of phi, psi, and omega angles (conformers) and single torsions sampled from QM-derived probabilities originally used for energetic evaluation of glycosidic linkages^25,29^. Conformer sets are dependent on each chemically distinct pair of saccharides making up a glycosidic bond, whereas single torsions depend on the anomeric chemistry of the linkage. We derived the conformers for this work by carrying out a new bioinformatic analysis of glycans in the PDB through the use of adaptive kernel density estimates in a similar manner to what was done for the 2010 Dunbrack Backbone-dependent Rotamer Library^41^ (see supplemental).

To optimize the conformations of glycan residues on different branches at the same time, the glycan tree is built layer-by-layer, with a layer defined as the residue distance to the root (Figure 1a). Once each new layer is built and optimized, all previous layers are then optimized further (Figure 1b). After all layers are built and optimized, a final optimization is conducted. The lowest energy model (decoy) found during this Monte Carlo algorithm is output at the end of the program as a PDB file. The lowest-energy structure of all the output decoys is used as the “best” model produced by the algorithm. See the supplemental material for more details [supp vid 1].

**Figure 1:**
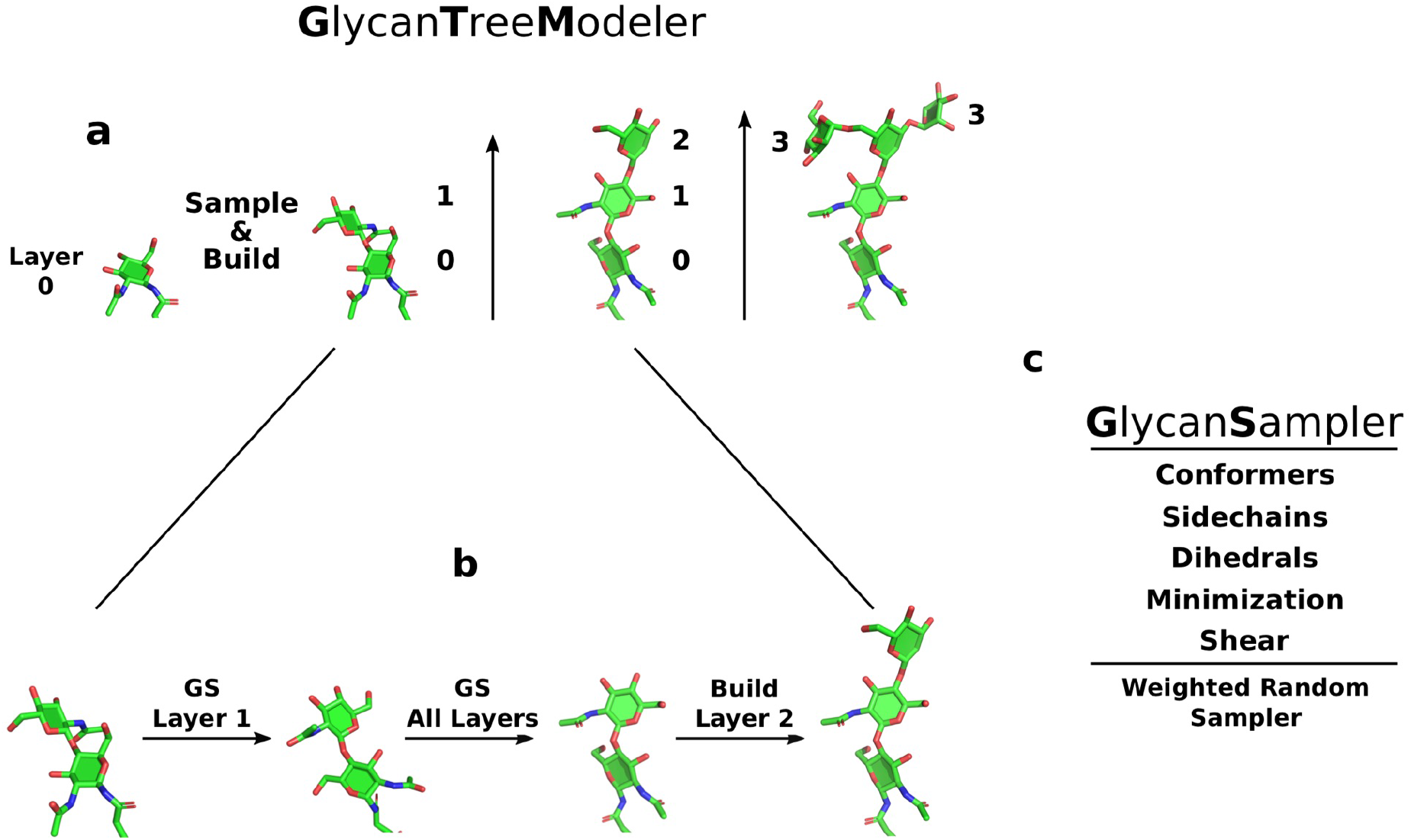
Glycan Modeling Diagram. a. Glycan trees building layer by layer. Numbers indicate distance to root of the glycan tree, which is the first residue. b. After a layer is built, Glycan Sampling is performed on the new layer, and then all layers, before building the next layer. c. Diagram showing major components of the GlycanSampler. The GS is a weighted random sampler, indicating that each DOF is sampled with a specific probability. See supplemental for details.

In order to examine the performance of GlycanTreeModeler, we built a new benchmarking infrastructure in Rosetta. We developed the *SimpleMetrics* framework within the XML interface to Rosetta (*RosettaScripts*^35^), which allows for robust analysis through more than 20 associated structural and energetic metrics, with data reporting at any step in a *RosettaScripts* protocol. The Python scripting language was used to load the resulting JSON scorefile for data analysis and figure creation using the numpy^42^, pandas^43^, and seaborn^44^ libraries. To facilitate large scale benchmarking, we developed a general application for parallel RosettaScripts computing, *rosetta_scripts_jd3*, enabling glycan calculations to be run in parallel on a high-performance computing cluster. This application can run multiple jobs within a single parallel run of Rosetta, with individually configured glycan trees to be modeled, and any associated input files for each. The *SimpleMetric* framework and *rosetta_scripts_jd3* application are reviewed in detail in the supplemental material.

Glycan masking was carried out through the use of two new *RosettaScript* components; the *CreateGlycanSequonMover*, which designs typical and enhanced^45,15^ glycan sequons into a protein at a desired position, and the *SimpleGlycosylateMover*, which adds whole glycans of a given IUPAC onto a protein. Glycans were then sampled using the *GlycanTreeModeler*.

### Glycan Structure Test Set

The Rosetta *GlycanTreeModeler* algorithm was benchmarked against a set of 25 unique N-linked glycan trees ranging from three to twelve residues, across 19 unrelated glycoprotein structures of better than 2 Å resolution, totaling 139 sugar residues. Each glycan tree was checked for chemical and structural inconsistencies (such as incorrect isoform assignments, wrong linkages, or missing atoms) using the glycosciences.de pdb-care webserver^27^. Preparation and analysis of the structures can be found in the supplemental material.

To assess the predictive capability of the *GlycanTreeRelax* algorithm, the dihedral angles of the glycans are randomized at the start of the algorithm, and waters are removed. Models are compared to the crystal structures using the all-heavy-atom Root Mean Square Deviation (RMSD) metric, with the lowest energy model of all output decoys used for assessment (Figure 2). The RMSD is calculated on all glycan residues that have an acceptable fit to the density in the native model, as terminal glycan residues of some glycans often cannot be observed in the density due to their higher flexibility. A description of the methods used for the RMSD calculation is provided in the supplemental.

**Figure 2:**
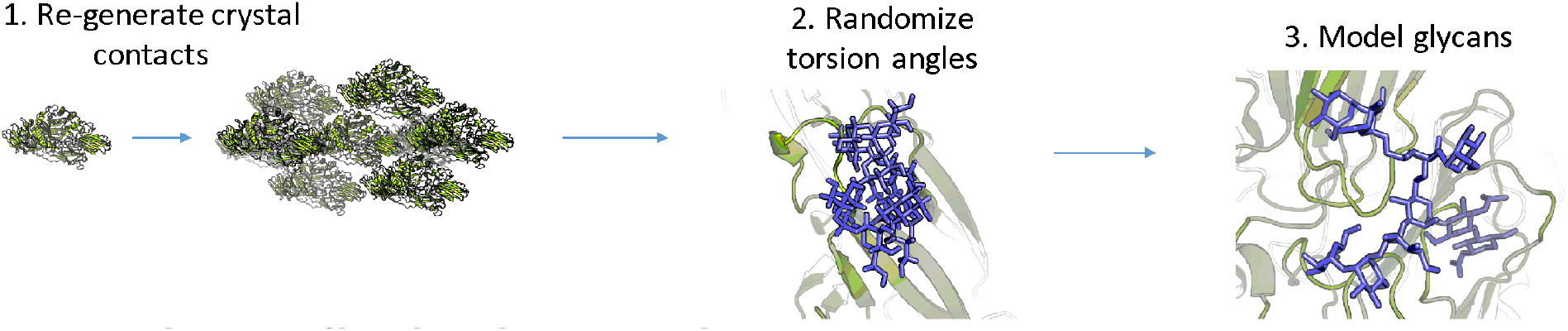
Schematic of benchmarking protocol.

### Protocol Optimization

Development of the glycan modeling protocol began with the implementation of the *GlycanSampler*. Initial results showed that the sampler alone produced models that were energetically favorable, but most final models were well above 5 Å RMSD from native—the mean and median RMSD values over the benchmark set were 7.6 Å and 7.2 Å, respectively. For some of the more sterically confined input glycans, many of the decoys had major clashes in their first few glycan residues, indicating that sampling of these residues was insufficient, even after increasing the overall amount of sampling.

To correct for the sampling problem, we modeled our algorithm after the growth of natural trees, in which we kinematically build and sample the glycan layer-by-layer, essentially “growing” a glycan “tree”. We defined a layer as the number of residues to the glycan root to enable branched glycan residues to sample conformations together. This *build-by-layer* algorithm improved enrichment of near-native output models and decreased the median RMSD to 6.1 Å, but did not improve the overall mean. (Figure 3). We then sought to systematically improve the algorithm through iterative benchmarking and optimization of kinematic and energetic experiments.

**Figure 3:**
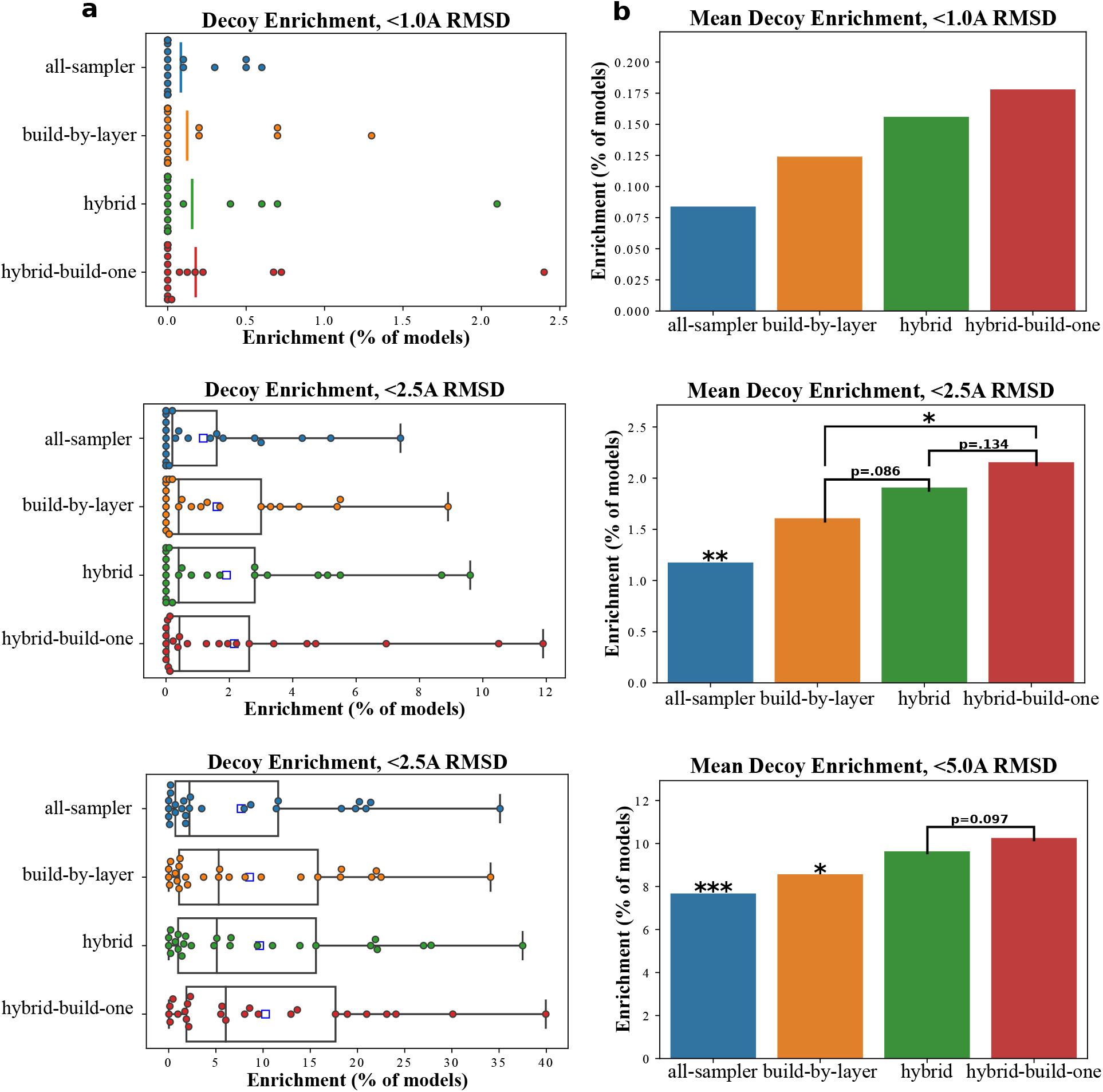
Kinematic Sampling Optimization, Decoy Enrichment from each individual experiment. All experiments were conducted with the same total amount of sampling. **a**. Boxplots without whiskers at decoy enrichments of <1A, <2.5A, and <5.0A. **b**. Means of decoy enrichments of <1A, <2.5A, and <5.0A. Asterisks indicate statistically significant differences through paired t-test. Asterisk above bar indicate statistical significance with all other groups. **\***,p <.05; **\*\***,p<.005; **\*\*\***,p<.0005

The original build-by-layer algorithm builds two layers at a time, with an overlap of one layer (window). Although this algorithm improved enrichments compared to the *GlycanSampler* alone (*all_sampler*, Figure 3), once a layer is built and the overlap refinement is complete, those layers are not optimized further, which makes refinement of the overall orientation of the glycan difficult, especially for large, branching glycans (which performed worse using this algorithm).

Two protocols were tested that include the build-by-layer protocol with more optimization of previously built layers. The *hybrid* algorithm builds and optimizes glycan layers as before but optimizes all previously built layers before the next build occurs. This algorithm significantly improved enrichments for all near-native definitions (Figure 3). The *hybrid-GS* algorithm is a simplified protocol implemented in *RosettaScripts* that splits sampling time across the first two tested algorithms. It first runs *build-by-layer* and then runs the *GlycanSampler* for optimization. This protocol did not improve enrichments, indicating that additional sampling of previous layers during the build process instead of after is important for improved model quality (figure S3).

Finally, since the *hybrid* algorithm is refining previously built layers, we removed the window sampling and benchmarked the number of build layers. By building a single layer at a time (*hybrid-build-one*), we further improved decoy enrichments (Figure 3); however, building two layers at a time did not improve enrichments (Figure S4).

Each of the major kinematic experiments generally improved near-native model quality (Figure 3, S5, and S6), but this was much more pronounced for the stem region of the glycan, defined as the first two layers of the glycan tree (Figure S7). Through better optimization of the base region through kinematics, the overall quality of output models was improved. We then sought to improve decoy discrimination of these low-RMSD models through improvements to the scoring function.

### Scoring Optimization

The default Rosetta energy function is composed of many individual energy terms, each with an associated, optimized weight^37^. A core component of the *RosettaCarbohydrate* framework is a specific energy term for the carbohydrate backbone, analogous to that of the Ramachandran term used for peptide bonds. This QM-derived term is used to improve backbone geometry arising from anomeric stereochemistry of both glycan residues in the bond^18^ and is on by default at a weight of 1.0 when working with glycans in Rosetta. We first sought to find a balance between the overall energetics of the glycan and the penalties arising from this term when native geometries are not ideal. An initial small test of various weights of this term indicated that a weight of 0.5 resulted in better decoy discrimination through the PNear metric, though this was not statistically significant (Figure S8). However, comparing the weight of 1.0 and 0.5 using a larger benchmark did result in statistical significance at a lambda of 1 Å (Figure 4).

**Figure 4:**
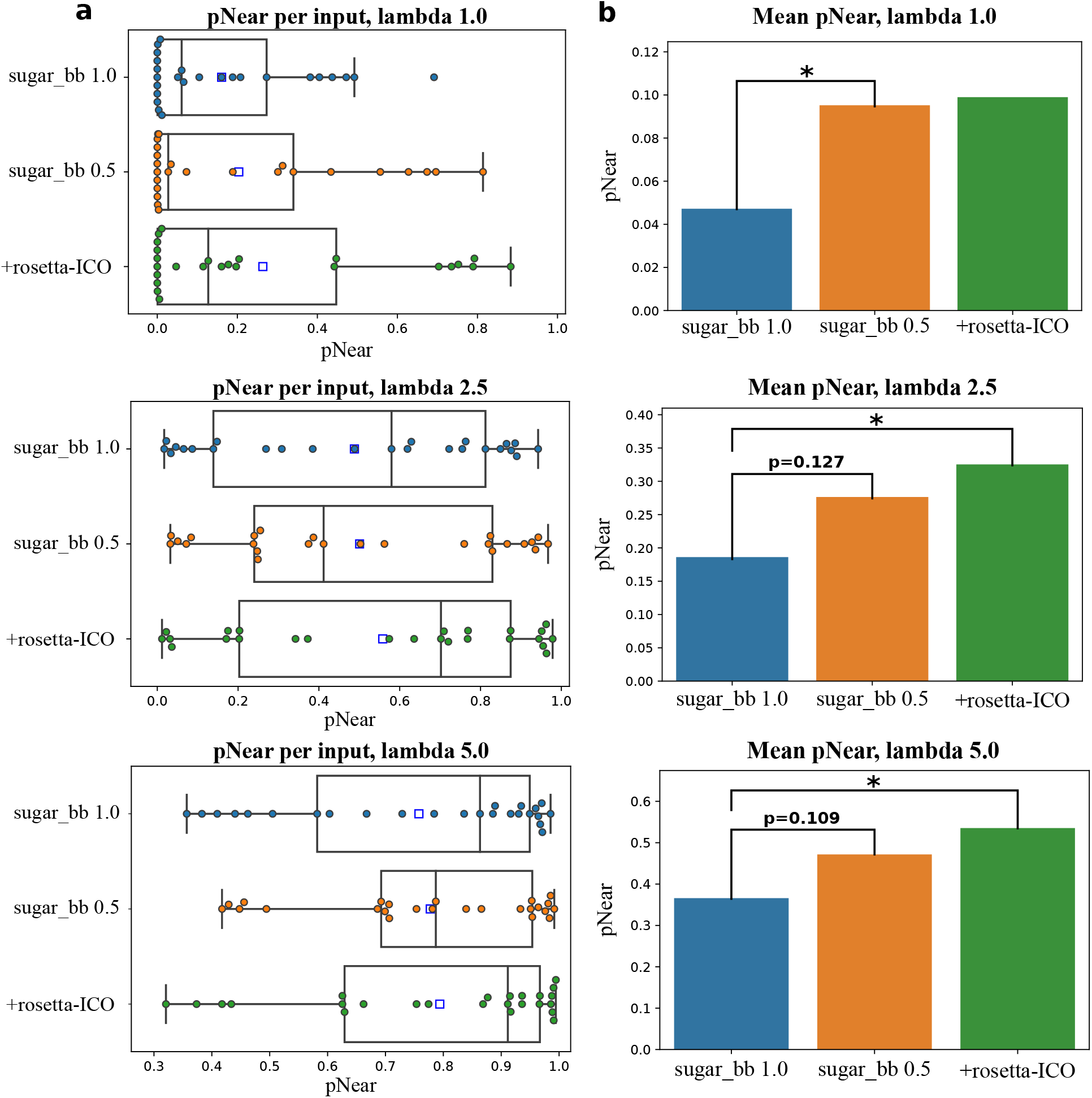
Funnel Plot quality of Scoring Benchmarks assessed by the pNear metric. **a**. Boxplot of pNear values for each benchmark glycan, indicating funnel plot quality for lambdas of 1.0, 2.5, and 5.0 RMSD to native. Higher pNear indicates better near-native discrimination from other decoys. Blue squares indicate mean. **b**. Means of pNear over each experiment. Significance from paired t-test; **\*** indicates p <.05.

We then sought to improve the overall discrimination through the recently developed Rosetta-ICO (*beta*) energy function^46^. Among other improvements, this energy function includes an atomistic repulsive term within residues (*intra_rep*), and a more accurate implicit solvation model that takes into account potential bridging waters – both of which are important considerations for carbohydrate modeling.

Each of these energy function changes improved decoy discrimination through the PNear metric. The *sugar_bb* energy term change was statistically significant at a lambda of 1.0 Å, indicating that too high of a *sugar_bb* weight reduces the energy function’s ability to discriminate near-native models from decoys (Figure 4). Notably, both optimizations together improved decoy discrimination for lambdas of both 2.5 and 5.0 Å, which can help distinguish poor quality models from acceptable ones. This improvement was statistically significant compared to the base energy function of ref2015.

Although these improvements were observed in decoy discrimination, using the Rosetta-ICO energy function (*beta*) in combination with a lower *sugar_bb* weight also improved enrichment of near-native models, especially in the root/STEM region (First two sugars, Figure 5). These changes also directly influenced the quality of the final models, most likely due to improvements in PNear (Figure S9).

**Figure 5:**
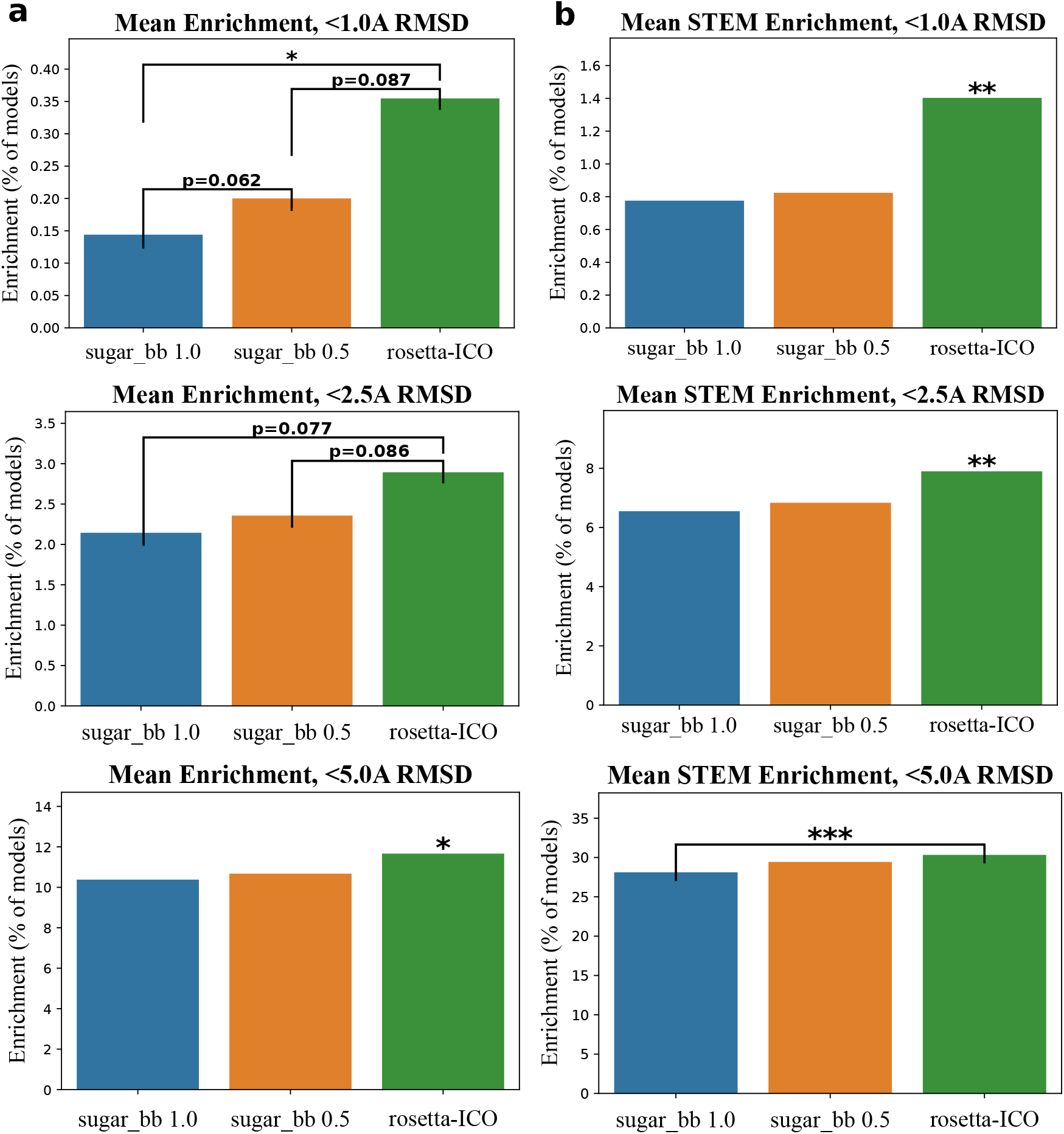
Scoring Optimization, Decoy Enrichments of each experiment. Asterisk above bar indicate statistical significance with all other groups through paired t-test. **\***|p <.05 **\*\***|p<.005 **\*\*\***|p<.0005 **a**. Decoy Enrichment in output models at <1.0A, <2.5A, and <5.0A RMSD. **b**. Decoy Enrichment in output models of the base (STEM) region indicating layers 0 and 1.

### Benchmarking of De novo modeling

Using the optimized protocol and scoring function found during protocol optimization, benchmarking was done on the set of 25 glycans described above. Across the benchmark dataset, the median RMSD of the glycan predictions to the native structures was 2.7 Å, while the mean was 5 Å. For the first two residues of the glycan tree, the median was 1.28 Å with a mean of 2.17 Å. Of the 25 glycan trees, 20% of the glycans were predicted at < 1 Å accuracy and 72% (18/25) of the glycans were predicted at < 5 Å accuracy (Figure 6 and 7). The largest glycan in our dataset, with twelve residues, was benchmarked at 2.5 Å. Full results for each glycan are listed in Table S3.

**Figure 6:**
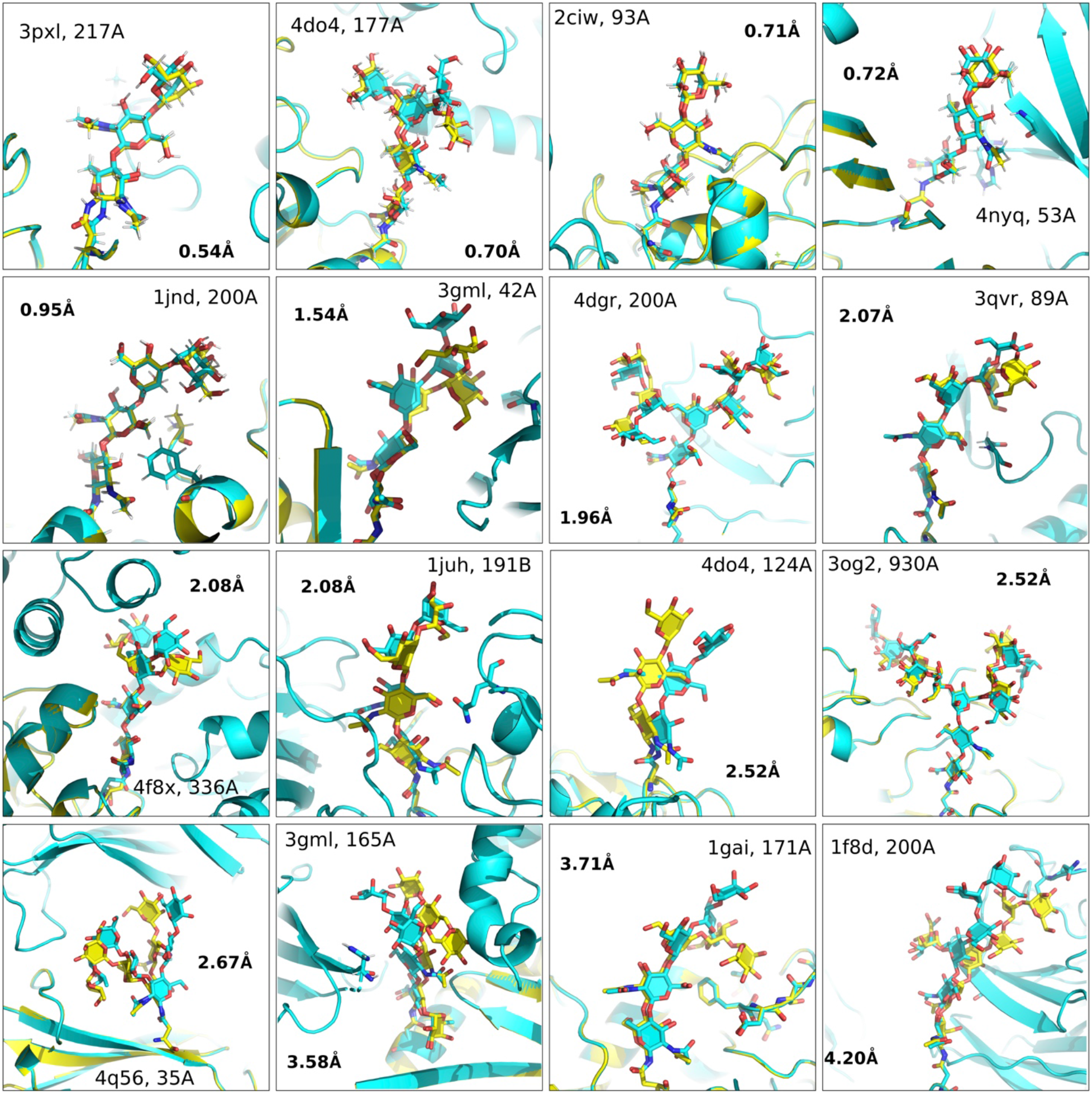
Near native structures from *de novo* modeling. (Top Scoring models for each glycan in the benchmark set) Yellow=Native, Cyan=Model

**Figure 7:**
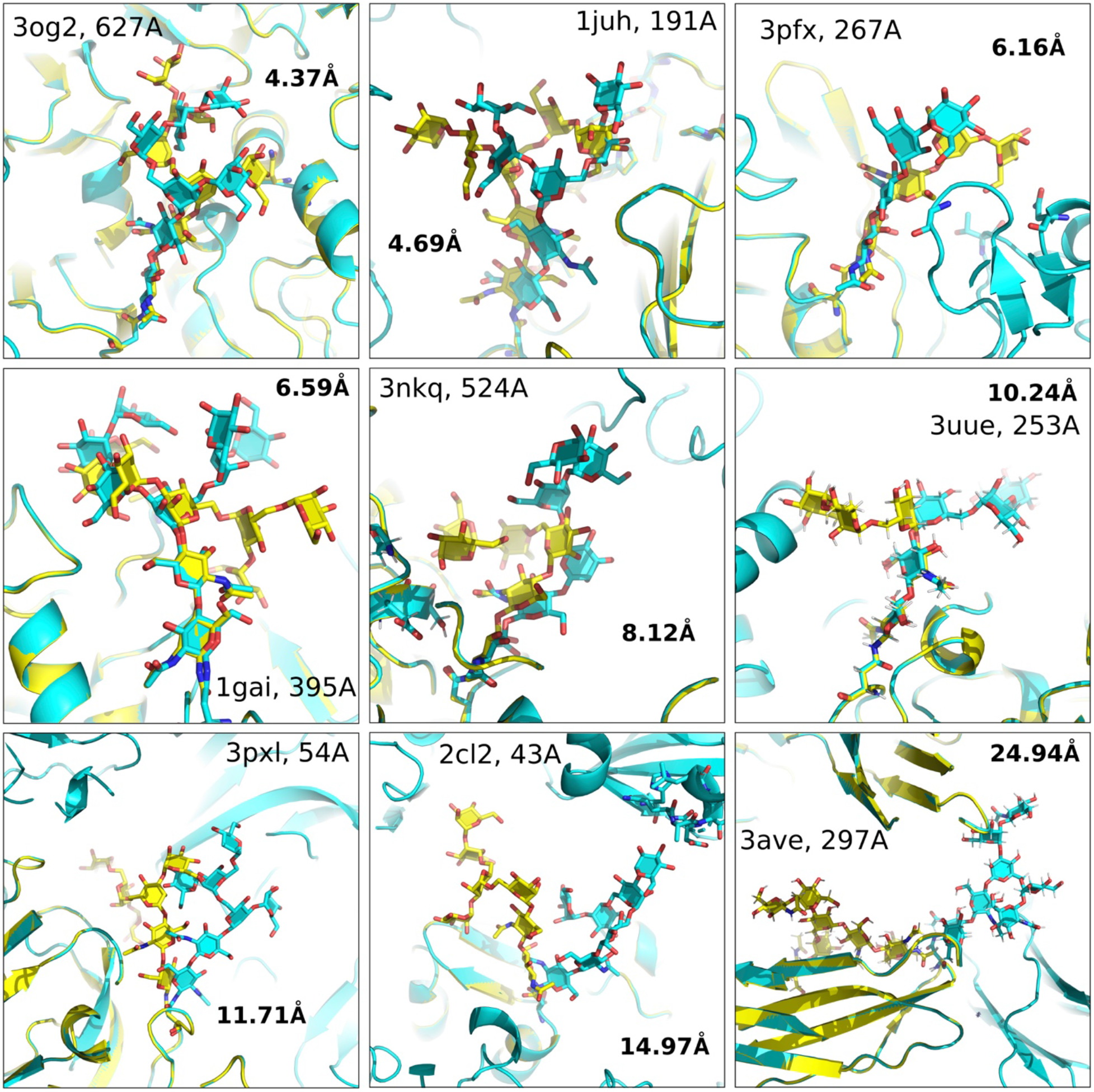
*De novo* predictions, Farthest from native. (Top Scoring models for each glycan in the benchmark set) Yellow=Native, Cyan=Model

It is also useful to understand how well the algorithm predicts the internal structure of the glycans, as a single dihedral angle change at the root of the glycan can significantly change the overall structure of the glycan relative to the protein. For each of these structures, the same lowest-energy models were superimposed onto the input glycan. The median superimposed RMSD is 1.1 Å, with a mean of 2.7 Å. Overall, 32% (8/25) were < 1 Å RMSD, 64% < 2.5 Å RMSD and 92% of the predictions < 5 Å. Both RMSD measurements of the glycans were generally correlated to each other (Figure S10).

In addition, most of the glycan benchmarks in our dataset had convergent score *vs*. RMSD (funnel) plots (Figure S11). This funnel-like quality is directly related to the ability of the scoring function to discriminate near-native models from decoys and was quantified using the PNear metric^47^ that estimates the Boltzmann-weighted probability of finding a system near its native state at various near-native cutoffs (lambdas) [See supp]. A PNear closer to 1.0 indicates the highest quality funnel possible. The worst-performing glycans in our benchmark set had poor score *vs*. RMSD funnels, indicating that the scoring function was not able to capture important biophysical properties of the structure (Figure S12). The worst-performing glycan from the Fc antibody fragment of 3ave, had an RMSD of almost 25 Å with an internal (superimposed) RMSD of 3.6 Å. In this lowest-scoring model (and others), the modeled glycan interacts with the more hydrophilic surface of a crystallographic symmetry mate rather than the more hydrophobic glycan-interacting surface of the parent protein that includes two aromatic rings (Figures 7, S13). This result is further detailed through the low pNear metrics of the funnel plot with all lambdas being less than .01, showing that the current energy function is unable to score these types of interactions well. However, a scoreterm that accurately represents glycan-aromatic CH-π interactions^48^ may improve these results.

Solvent is implicitly represented in most Rosetta applications, but we observe that half of the benchmark glycans have significant crystallographic waters in contact with the surrounding protein. Attempting to understand the effect of waters, we modeled the worst-performing and best-performing glycans and then predicted explicit waters around the glycan for each output decoy using Rosetta-ECO^46^ in order to score more native-like conformations that have these bridged waters. However, decoy discrimination as measured by pNear was significantly worse for all lambda cutoffs (even for the best-performing glycans), indicating that even with explicit waters and sufficient near-native sampling distributions, the Rosetta energy function was unable to use this information to accurately distinguish near-native decoys. (Table S4)

In the benchmark set, the internal (superimposed) RMSDs are generally low in comparison to the overall RMSD (84% < 3 Å), showing that the energy function, guided by the QM-derived *sugar_bb* energy term, can accurately predict many glycan structures, but may need to be further improved to more accurately score glycan-protein interactions in the future.

#### Density Building

There are an increased number of glycoprotein structures being determined. To assist structure determination, many recent glycan modeling tools have focused on their ability to aid in glycan structure building and refinement using the experimental density, especially for structures with many resolved glycans such as HIV Env. We tested the ability of the *GlycanTreeModeler* to build glycan structures using crystallographic density information to guide modeling and decoy discrimination using integrated density scoring^40^. The experiment was conducted in the same manner as *de novo* modeling, by first randomizing all backbone dihedral angles of the glycan to be modeled for each output decoy and removing all crystallographic waters. For each of the 25 glycans, the lowest-energy model was used for assessment.

Without further refinement or any additional changes to the protocol, all glycans were modeled at sub-angstrom accuracy. The best glycan in the current benchmark, with six residues, was built at 0.08 Å RMSD to native (3GML position 165A glycan), while the worst, a five-residue glycan, was modeled at 0.88 Å RMSD (1GAI position 171A glycan). For both of these glycans, funnel plots were generally good, with respective PNear values of 0.99 and 0.46 at a lambda of 1.0 Å (Figure 8). For 1GAI glycan 171A, the last residue in the glycan is twisted in the best model compared to the native and fits two constituent oxygens into the low residue density at a different angle than the solved structure. This twist can clearly be seen in the funnel plot where the distribution of models less than 1 Å is bimodal, indicating two primary close solutions of the electron density. (Figure 8F).

**Figure 8.**
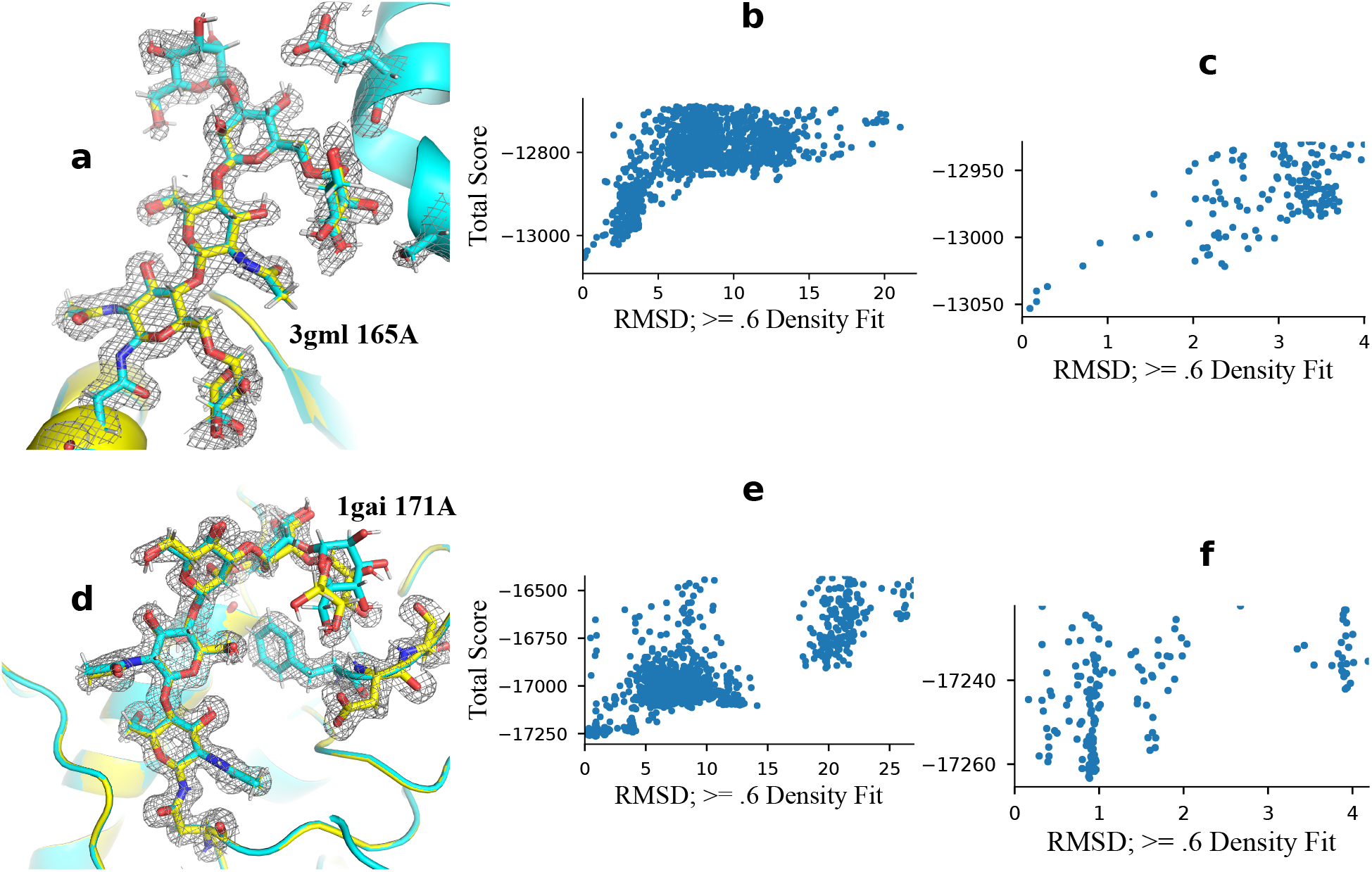
Best and Worst results from Density-guided modeling: **a**. Structural comparison of 3GML 165A glycan; **0.08A RMSD;** *cyan=model* | *yellow=native* **b (and e)**. RMSD *vs*. Score (funnel) plot, top 80% by energy. **c (and f)**. Funnel plot of top 10% models with pNear metrics. **d**. Structural comparison of 1GAI 171A glycan; **0.88A RMSD;** *cyan=model* | *yellow=native*

Overall, the *GlycanTreeModeler* achieved a mean heavy atom RMSD of 0.48 Å using all residues and 0.34 Å using residues that had acceptable fits into the density (133/139 total glycan residues, see supplemental). For both inclusion types, the median RMSD was 0.31 Å and 0.28 Å respectively, while the mean RMSD of the glycan root (first two sugar residues) was .23 Å (Figure 9a) (Table S5). Values for PNear with lambda of 1.0 Å were generally quite favorable, indicating high-quality funnels, with a mean of 0.86 and median of 0.92 (Figure 9b). These results show that the *GlycanTreeModeler* can be effective for modeling known glycans into electron density, especially with available methods for refinement.

**Figure 9.**
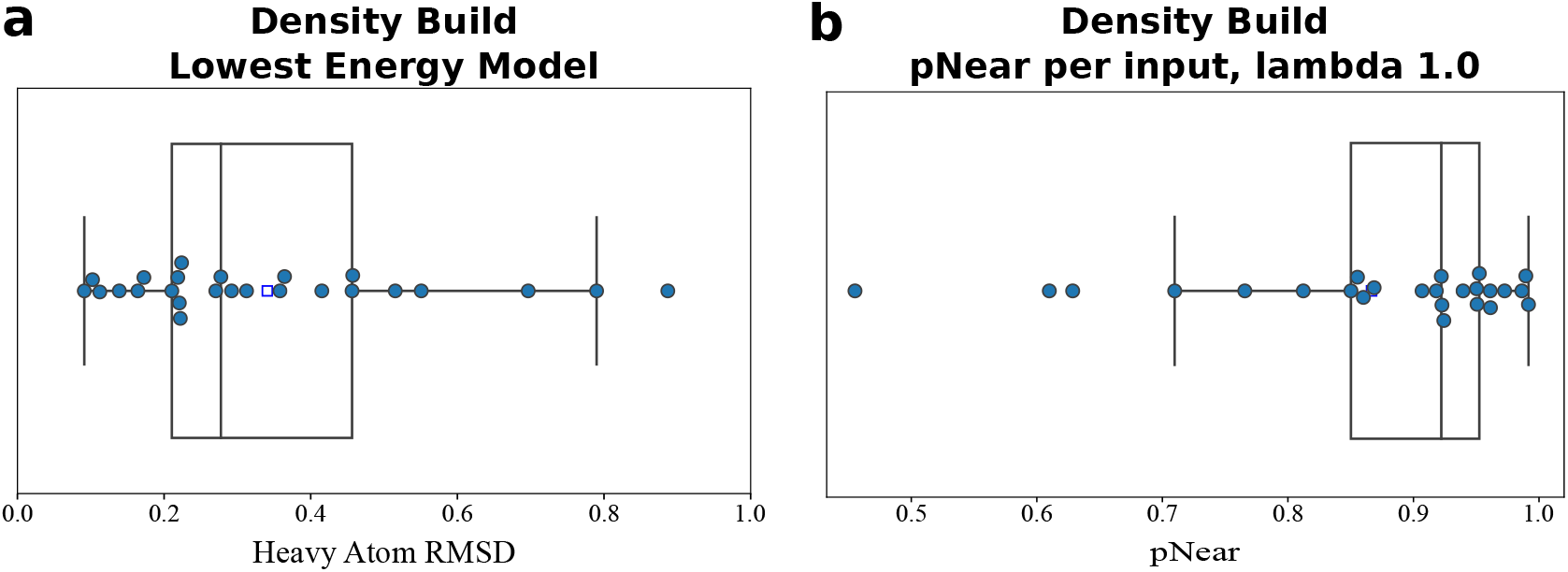
Density-guided Modeling Quality: **a**. Boxplot of the RMSD to native of the best-scoring decoy for each of the benchmarked input glycans. **b**. Boxplot of the funnel quality for each of the benchmark glycans as measured by the pNear metric. A value closer to 1.0 indicates a high-quality funnel.

#### Sugar coating protein surfaces

Addition of glycans to exposed protein surfaces can reduce B cell receptor access to underlying surface epitopes; this approach (called “glycan masking”) has been used to decrease the amount of antibodies elicited against off-target epitopes of designed immunogens^12,49,50,51^. Given the predictive capability of the *GlycanTreeModeler* to accurately model the spatial arrangement of complex glycans, we used the algorithm in combination with *RosettaScript SugarCoating* methods for sequon design and computational glycosylation to design four N-linked glycans onto the outer surface of the I53-50A trimeric component of the I53-50 protein nanoparticle scaffold (Figure 10A; details of the design approach are described in *Materials and Methods of the supplemental text*). I53-50 was selected as a model immunogen because it is currently in clinical trials as the nanoparticle scaffold for SARS-CoV-2^52^ and RSV^53^ vaccines.

**Figure 10.**
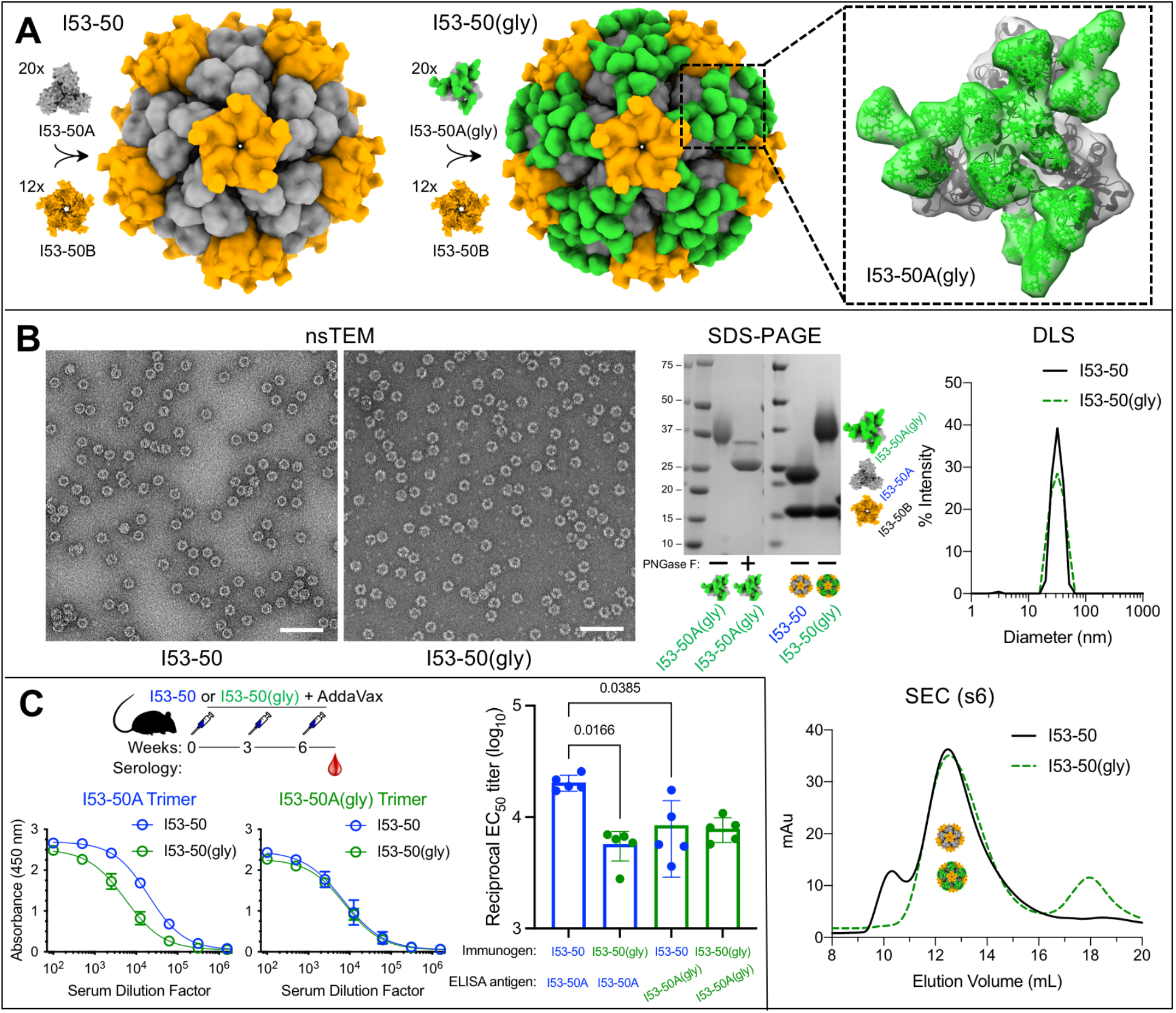
**(A)** Schematic of protein design models. On the left, twenty I53-50A trimers (gray) and twelve I53-50B pentamers (orange) self-assemble into I53-50 protein nanoparticles^54^. Rosetta sugarcoating design protocols were used to glycosylate the outer surface of I53-50A trimers with 4 N-linked glycans (green) per protomer to form I53-50 particles with 240 N-linked glycans (middle). The inset on the right is a close-up view of glycosylated I53-50A trimers with 12 total glycans on the outward-facing surface. **(B)** Characterization of bare versus glycosylated I53-50 particles using negative stain transmission electron microscopy (nsTEM; scale bar, 100 nm), SDS-PAGE, dynamic light scattering (DLS), and size exclusion chromatography (SEC) on a Superose 6 Increase 10/300 GL column (GE Healthcare). In the SEC chromatogram, both I53-50 and I53-50(gly) particles reach peak elution at 12.5 mL; unassembled I53-50A and I53-50B components elute at ∼18 mL. **(C)** ELISA curves (left two plots) and corresponding EC_50_ titers (right bar plot) showing reduction in anti-I53-50A antibody responses when mice were immunized with I53-50(gly) versus I53-50. BALB/c mice were immunized intramuscularly at 0, 3, and 6 weeks with 5.57 μg of I53-50 or I53-50(gly) and serum antibody binding to I53-50A trimer (left) or I53-50A(gly) trimer (right) was quantified via ELISA using 8-week sera (N=5 mice/group). For statistical analysis, Mann-Whitney tests were used to compare among the experimental groups.

When glycosylated I53-50A trimers and I53-50B pentamers were mixed *in vitro* at equimolar concentrations, the two components self-assembled into I53-50(gly) nanoparticles that display 240 glycans on the outer surface (Figure 10A, B). Biophysical characterization by negative stain transmission microscopy (nsTEM), dynamic light scattering (DLS), and size exclusion chromatography (SEC) confirmed the formation of monodisperse particles with the known I53-50 morphology (Figure 10B). SDS-PAGE analysis of the I53-50A(gly) trimer treated with PNGase F confirmed that the designed glycans were present in the protein (Figure 10B). Mice were immunized three times with 5.57 μg of I53-50 or I53-50(gly) particles. Anti-I53-50A trimer serum antibody titers were significantly lower in mice immunized with I53-50(gly) particles compared to mice immunized with I53-50 particles, whereas anti-I53-50A(gly) trimer titers were unchanged between the two groups (Figure 10C). These data demonstrate that the methods presented here can be used for glycan tools for glycan masking by modeling the spatial arrangement of putative glycans on protein surfaces.

## Discussion

The *GlycanTreeModeler* and associated tools allow modelers to accurately model glycans of interest through *de novo* and density-guided modeling. The algorithm and energy function were rigorously optimized and benchmarked with glycans of varying length and complexity at a median *de novo* RMSD of 2.7A. In fact, even before full optimization and release, the GlycanSampler algorithm (previously the *glycan_relax app*) was used to model glycans on HIV^55^, Hepatitis C^56^, vaccine candidates^57,58^ and (with the final optimized version) SARS-CoV-2^59^; illustrating the general utility of the algorithm.

The modular nature of Rosetta and the tools created for this work allow them to be used in a variety of complex modeling and design tasks. The *GlycanTreeModeler* was used with previously published density tools^40^ to build glycans into their crystallographic or cryoEM experimental density with sub-Angstrom accuracy. However, while the results are encouraging, a truly automated solution for glycoprotein modeling must also sample glycan chemistries, branching, and kinematics simultaneously in order to build potential glycan residues into the density of unknown glycans. Knowledge of the range of glycoforms and occupancy occurring at a glycosylation site can be obtained through mass-spectroscopy techniques^19,60^, but due to chemical and structural heterogeneity at any single glycan site, modelers will typically need to build models for multiple different glycoforms at a single site, especially for complex glycans. The tools presented here can sample and build multiple potential whole glycans at a site through the *SimpleGlycosylateMover*, but core Rosetta methods that also consider species and cell-type dependent glycan chemistries during the *GlycanTreeModeler* or end-to-end deep learning methods would be a welcome addition to the methods presented here.

By combining the tools through *RosettaScripts*, it becomes possible to computationally design glycan sequons at ideal positions on a protein, and then build and model multiple potential glycans at a variety of sites in a symmetric manner. This general workflow was used to sugarcoat a clinically relevant nanoparticle vaccine scaffold with N-linked glycans. In vitro and in vivo testing of this glycosylated scaffold showed a decrease in the humoral immune response to the glycan-masked surface. Sugar coating therapeutics using these methods could potentially reduce off-target effects of many preclinical biologics, especially with respect to immunogenicity.

Most glycans are can sample a wide range of conformations in solution, as they are mostly polar, usually exposed to solvent, and have many conformational degrees of freedom. Thus, accurately predicting the lowest energy states (and highest occupancy conformations) for glycans is difficult. While we can generalize that low energy conformations found through the *GlycanTreeModeler* should be indicative of probable solution conformations, the *GlycanTreeModeler* was not benchmarked on an experimental ensemble of glycan structures. The few glycan ensembles found through solution NMR^61^ may approximate conformational ensembles in solution and could be the bases for future benchmarking. However, even with this consideration, many of the benchmark glycans that were modeled accurately to their crystal structures are not hindered by monomer or crystal contacts, but have interactions to protein in their glycan root. Additionally, predictions of the internal (superimposed) RMSDs of all glycans benchmarked were generally favorable with a median benchmarked accuracy of 1.1 Å and a mean of 2.7 Å, indicating that the glycan root, subsequent torsional preferences, and intra-glycan interactions may be determining structural factors for these isolated glycans.

Although the algorithm is capable of accurate *de novo* modeling of many glycans (especially at their base) and has been used for experimental glycan masking, there is certainly room for improvement. In nearly all of the benchmarks, the native structure is sampled adequately, but in a subset of structures, the energy function is not able to choose near-native structures. Upon further investigation of the many native glycans in the benchmark set with water-mediated hydrogen bonds, we originally hypothesized that explicit water modeling might help the energy function discriminate near-native models. However, we found that implicit modeling actually led to better discrimination scores through the pNear metric. In order to improve the algorithm further, the Rosetta energy function will need to be optimized to improve glycan-protein interactions, specifically in terms of hydrogen bonds, solvation, and the introduction of energy terms that better represent aromatic CH-π interactions^48^. Finally, the algorithm requires more compute time as the number of glycans to model increases, which can be prohibitive for large, multimeric glycoproteins such as HIV.

In this work, optimization of both sampling and scoring was necessary to improve overall accuracy. A key component of the algorithm is the nature-inspired kinematics used during sampling, which was shown to be an important determinant of the overall accuracy of the algorithm. The kinematics were rigorously benchmarked here, though kinematics are not always taken into account or optimized in state-of-the-art classical modeling algorithms. This benchmarking was made possible by the SimpleMetric framework and a new *RosettaScripts* application that were created and used continuously throughout this work.

SimpleMetrics have now become a critical tool for general analysis in Rosetta and as a way to export important information for external algorithms, such as the quantum annealer^62^. As core protocols in Rosetta continue to be optimized, and as deep learning becomes a more integral aspect of modeling and design, SimpleMetrics should allow the robust analysis of new protocols, results, and Rosetta benchmarks, as it has for this work.

These results show that the *GlycanTreeModeler* is able to accurately predict glycan structures *de novo*, build them into known density, and be used in *SugarCoating* protein surfaces. In addition, the modular nature of the components allows them to be further developed for specific engineering tasks such as immunogenicity reduction or the optimization of developability characteristics such as half-life, solubility, and aggregation potential.

## Supporting information

glycan_conformer_table

## Availability and Documentation

The *GlycanTreeModeler, GlycanSampler*, and all tools used in this work are available in the Rosetta Software Suite, which is free for non-commercial use. All tools are available as components for *RosettaScripts* and PyRosetta. In addition, the use of all components are covered in publicly accessible tutorials^63^ and detailed protocol captures^64^. Results of this study are continuously benchmarked using the Rosetta automated scientific testing framework^65^.

## Figures

Figures were created using matplotlib^66^. Glycans were visualized in PyMol using the Azahar plugin^67^, which was expanded for this work: https://github.com/BIOS-IMASL/Azahar/pull/17

### Documentation Links

- **RosettaScripts**
  - https://www.rosettacommons.org/docs/latest/scripting_documentation/RosettaScripts/RosettaScripts
- **Working with Glycans:**
  - https://www.rosettacommons.org/docs/latest/application_documentation/carbohydrates/WorkingWithGlycans
- **Chapter 13 of the PyRosetta Notebooks**:
  - https://github.com/RosettaCommons/PyRosetta.notebooks

## Acknowledgements

We gratefully acknowledge Rashmi Ravichandran for providing I53-50B pentamer, Deleah Pettie and Michael Murphy for assistance with expression of I53-50A(gly) design models, Alex Roederer for preparation of I53-50 and I53-50(gly) particles for mouse immunization, and Minh N Pham for performing mouse immunizations and blood draws.

