## supplemental_text_and_figures for "Growing Glycans in Rosetta: Accurate *de novo* glycan modeling, density fitting, and rational sequon design"

### Detailed Methods and Results [Supplemental]

**PNear .** This metric quantifies score *vs.* RMSD funnel quality and was developed by Bhardwaj, Mulligan et al[1] . It is used in this work to quantify decoy discrimination and aid in score function optimization. The equation is as follows:

$$PNear = \frac{\sum_{i=1}^{N} e^{-\frac{rmsd_i^2}{\lambda^2}} e^{-\frac{\Delta E_i}{k_B T}}}{\sum_{j=1}^{N} e^{-\frac{\Delta E_j}{k_B T}}}$$

Where lambda (**λ**) determines the size of the Gaussian, with lower values defining a more restrictive notion of what is close to native. For this work, different lambdas are used to quantify funnel quality at different definitions of 'near-native', typically at values of 1.0 Å , 2.5 Å, and 5.0 Å RMSD.

**kB*T** is set to 1.0.

**N** is the total number of decoys.

**ΔE** is the $\text{RosettaTotalEnergy}_{\text{model}} - \text{RosettaTotalEnergy}_{\text{lowest}}$

***RMSD Calculations.** Within solved crystal structures, not all atoms or residues fit well into the experimentally determined density. This is especially true for glycan residues, where some residues are more mobile than others, resulting in poor density. In order to accurately represent structural deviations when benchmarking glycan trees, the crystal density was used to assign a 'fit' score to each glycan residue in a benchmark glycan tree. This fit score ranges from 0 to 1, with values of .8 meaning a high fit to density, and values less than .6 being poor fits. The density fit metric is generally equivalent to the coot density fit analysis. All RMSD values reported in this manuscript (including pre-aligned RMSDs) use only residues that have >= .6 correlation to the density unless otherwise noted. For the 25 glycan trees used in benchmarking, only 6 glycan trees had an outer residue that did not fit well into the density. An example of the density fit of is below.

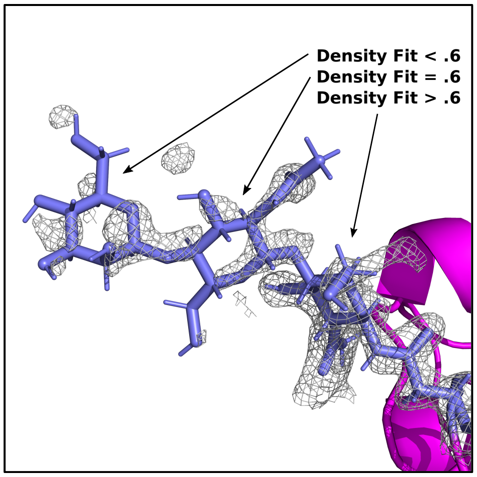


**Figure S1: Density Fit Example**

RMSDs were calculated using the *RMSDMetric*, while density fit was calculated using the *PerResidueDensityFitMetric* of the ***SimpleMetric*** framework. Residues were selected for RMSD calculation using the *DensityFitResidueSelector* and passed onto the *RMSDMetric* for calculation.

**Superposition.** For internal RMSD comparisons where alignment was carried out prior to the RMSD calculation, the whole glycan tree of the decoy model was aligned to the input glycan tree using all atoms of each tree through the *super* option of the *RMSDMetric.*

### Conformer Generation

The data used in the original *RosettaCarbohydrate* framework only has data for the most common saccharide chemistry and contains only a subset of the data that is currently available in the PDB[2,3].

In order to update the underlying data used for conformer sampling, we were provided per-linkage torsion data from Glycosciences.de[4] reflecting the PDB as of June 2017 including N-linked glycan torsions. Each linkage is unique in its reducing end and non-reducing-end amino acid or carbohydrate type (for *example beta-D-GlcpNAc ->4)-D-GlcpNAc*).

The raw data was then filtered for the following, resulting in 14,351 linkages across 64 unique linkage types:

1. Crystal structures only - No NMR or Unknown methods
2. <= 2.0 A resolution.
3. Full torsions for each linkage - Phi/Psi Phi/Psi/Omega, etc. No missing torsions.
4. At least 10 datapoints for a specific linkage.
5. Torsion quality. Linkages were skipped for the following reasons. Integers for each listing are categorical values reported from glycosciences.de dependent on the type of linkage:
    a. *Wrong assignment of anomeric carbon (3)*
    b. *PDB residue name and detected monosaccharide are inconsistent (3)*
    c. *Residue name given in PDB file is unknown in the list of residues (4)*
    d. *Stereochemistry of the carbohydrate could not be assigned (5)*
    e. *No glycosidic O -(S -, N -) atom at C1 could be detected (6)*
    f. *Residue at the reducing end of a chain is neither a carbohydrate nor one of the assigned substituents (10)*
    g. *Glycosidic linkage reported in the PDB is not consistent with the detected one (16)*
    h. *residue has no oxygen or respective atom attached to anomeric carbon (32)*
    i. *residue is linked to another residue (except ASN) by a non-oxygen glycosidic atom (1024)*
    j. *number of rings in oligosaccharide doesn't match expected value (4096)*
    k. *residue could not be assigned (65536)*
    l. *N-glycan chain does not match known biological pathway (524288)*

m. *a monosaccharide is linked to its parent one via a carbon other than the anomeric carbon (1048576)*

Since the glycoprotein conformers do *not* follow the pre-defined canonical configurations, such as *g+*, *trans* or *g-* rotamers at 60°, 180° and -60° respectively, observed for the most side-chain torsion angles of the 20 amino acids, we first sought to define these states following the protocol for the non-rotameric degrees of freedom of the standard amino acids[5]. For each chemically distinct glycan-glycan or amino-acid-glycan linkage type and each of the few (1~4) torsion angles modeling such linkage, we computed an adaptive Kernel Density Estimate (adaptive KDE) using a *von Mises* kernel from an available data set (Fig S2). Due to a limited number (mostly 10~100) of experimental conformations in each linkage data set, a probability density function of each torsion angle was estimated in a backbone-independent manner as described in Eq. 18 of the 2010 Rotamer Library study[5]. We then used the spline toolbox of *Matlab* to create cubic splines of the KDEs and solve for first and second derivatives in order to derive all inflection points (x where F'(x) = 0) and whether a local minimum (F''(x) > 0) or maximum (F''(x) > 0) is observed.

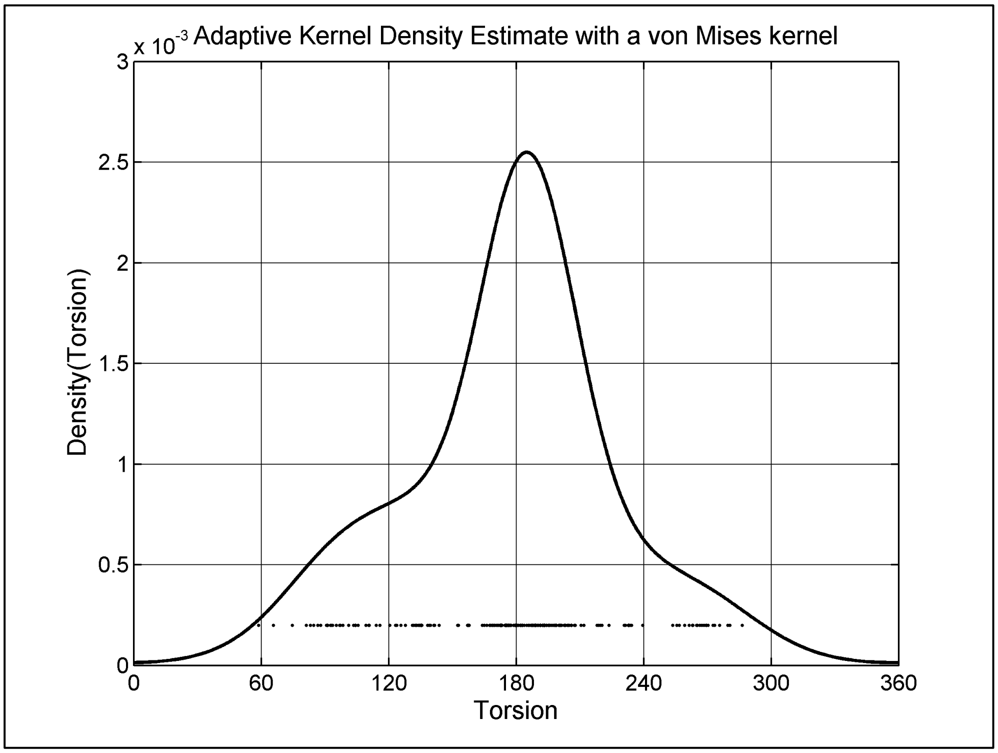


**Figure S2: Probability Density Function of a torsion angle calculated with adaptive Kernel Density Estimate.** The von Mises kernel allows for continuous circular description of the torsion angle distribution. The experimental angles from a sample are shown with small dots at the bottom. Such 1-D density estimates were performed for each torsion comprising a glycan-glycan or amino-acid-glycan linkage type. These 64 linkage types can be found in the resulting conformer table included in the supplemental materials.

The data for each torsion was then binned into one-dimensional conformers using the local maxima of a torsion distribution to define the conformer and based on the left and right adjacent minima to define the conformer boundaries given the circular nature of the angular data. The probability of each 1-D conformer was calculated by integrating the density from the left to right boundaries (processed data is included in this work).

In order to determine linkage conformers defined with the set of torsion angles making up a particular connection, the 1-D conformers were given integer values and a bin was assigned for each torsion in the original data using the boundaries of each 1-D conformer. For example, a 2-1-3 conformer would indicate that phi belongs in conformer bin 2, while psi belongs in conformer bin 1, and omega belongs in conformer bin 3. Probabilities of each full conformer were then calculated using these assignments and final means and standard deviations of each torsion that comprised the conformer were determined. This data was formatted into a parsable text file for use in Rosetta or other programs. The formatted data can be found as-is for further scientific use. All analysis outside of MATLAB was done in Python using standard modules and the SciPy module for circular computations[6] .

### Glycan Modeling

The *GlycanTreeModeler* (GTM) models a glycan in a directional manner, starting with the roots of the glycan and building and optimizing until the whole glycan tree is present. This is done through modeling of glycan 'layers', defined as the residue distance to the start of the glycan tree. We typically model one to two 'layers' at a time, going out to the outer foliage of the tree, however, this is customizable, and has been optimized to our final defaults.

The *GlycanTreeModeler* also defines a 'window', where overlapping of the residues from the previous layer can occur. For example, if we have a layer size of two, and a window size of 1, we would first model layers 1 and 2, and then 2 and 3, until all layers have been modeled.

Modeling a single glycan tree of 9 residues takes approximately 5 minutes per output structure and is typically run in parallel on a compute cluster where 5-10k structures are output and the lowest energy structure is determined and used as the final model. A video of this process where the layer size is set to one can be seen here [link to vid?]

Sampling of the layers is done through the Monte Carlo *GlycanSampler*. This includes sampling individual torsions through statistics derived from QM datana[7,8], sampling whole glycan linkages through updated conformer generation, and general Rosetta sampling techniques including the *ShearMover* (which was updated for carbohydrates), structure minimization, and sampling sugar OH groups, constituents, and any neighboring protein side-chains through the Rosetta side-chain packing algorithm[9] .

**DOF Sampling: GlycanSampler**. Sampling of glycan residues within the *GlycanTreeModeler* is done through the *GlycanSampler*. The *GlycanSampler* is a collection of individual components that collectively aim to sample the diverse DOFs involved in glycan structure. The sampler itself uses a set of weights, which chooses a particular DOF sampler using an associated probability, akin to the sampling in the SnugDock algorithm that is used to refine antibody structures[10]. The probability of selecting a particular DOF sampler each round is given in table S1. Probabilities for each sampler were selected manually in order to strike a balance between fast samplers like conformer sampling and slower samplers like whole-structure minimization and side-chain optimization.

Internally, the GlycanSampler runs a set number of rounds, which is multiplied by the total number of glycan residues set for optimization, in order to normalize the amount of sampling done for each residue. At the end of each round, the energy of the new structure is then assessed using the set *ScoreFunction* and the move is either accepted or rejected using the Metropolis Criterion, where $kT$ is set to a default of 2.0. The *GlycanTreeModeler* calls the *GlycanSampler* each time a new layer is built - first to optimize the new layer, and then to optimize all previous layers. Within the GTM, a common man5 glycan of length 7 would undergo approximately 1100 total sampling rounds before a single decoy is output.

Components of the *GlycanSampler* include sampling individual glycan torsions through probabilities derived from QM as described above, as well as small, medium, and large torsional moves; sets of glycosidic torsions of a particular residue (Linkage) through conformer sampling using updated statistics generated using the adaptive KDEs described above; shear-sampling that aims to reduce downstream effects of a torsional move; the rotation of the OH groups and any constituents in the tree as well as protein side-chain neighbors; and multi-residue minimization of the tree. All components (movers) that comprise the *GlycanSampler* are accessible in PyRosetta and most are accessible to RosettaScripts. A list of these components, their purpose, and their overall probability is listed in table S1.

**Table S1.** GlycanSampler Components and Probabilities

| DOF | Component | Description | Probability |
|---|---|---|---|
| Torsion | *SmallBBSampler* | Random backbone angle sampling. 3 Samplers, sampling at +/- 15, 45, and 90 degrees. Each sampler at a probability of 4:2:1 respectively. | 0.3 \| 4:2:1 ratio |
| Torsion | *SugarBBSampler* | Samples torsions using probabilities derived from QM | 0.2 |
| Conformer | *LinkageConformerMover* | Samples a set of torsions derived from the PDB and dependent on the chemical identity of each glycan in the glycosidic bond. | 0.2 |
| Shear | *ShearMover* | Torsional change and a counter-rotation that results in limiting downstream moves to small translations | 0.1 |
| Side-chains | *PackRotamersMover* | Optimize glycan constituents such as hydroxyls and rotamers of neighbor protein side-chains within 6Å | 0.1 |
| All Atoms | *MinMover* | Structure minimization on the energy function using the *dfpmin_armijo_nonmonotone* at a tolerance of .01 | 0.1 |

**DOFs: Conformer Sampling**. Conformers are a set of low-energy common conformations of a particular structure. Glycan conformers are dependent on the chemical identity of each residue in the glycosidic bond, the anomeric state of each (alpha/beta), and through which carbon atom the i+1 glycan is attached to (i.e. carbon 6 *vs.* carbon 4; making a 1-6 or 1-4 linkage). Conformers are sampled through the *LinkageConformerMover*[11].

At each application of the mover, a random residue in the set is chosen for optimization and a random conformer is chosen to improve sampling of uncommon conformations (even sampling) and reduce possible bias from the PDB. For each glycosidic torsion of the saccharide residue (defined using the IUPAC definition), a new value of the torsion is sampled through a gaussian function using the mean and standard deviation of the angle in the conformer. This is done for all backbone torsions of the residue; effectively replacing them with the conformer. Through the use of Rosetta's internal residue-based graph structure (FoldTree), downstream coordinates are updated in accordance with the connectivity of the glycan tree.

**DOFs: Glycosidic Torsion Sampling.** Individual glycosidic torsions are optimized through four backbone-optimizing movers with a new generalized backbone sampling framework that can sample an arbitrary number of backbone torsions within a particular residue. Each call to one of these movers first randomly chooses a residue and then a dihedral angle ($\Phi$, $\Psi$ , $\omega$, or $\omega_2$ where applicable ) within the backbone of that residue. Each dihedral is independent and can have a different set of parameters and data associated with them.

The movers used for torsional optimization include the *SugarBBSampler*, which uses torsional probabilities derived from QM, and a set of general *SmallBBSampler*s that sample torsions within a set delta of the current angle.

The *SugarBBSampler* optimizes individual glycosidic torsions using probabilities derived from the QM-derived *Ramachandran-like* sugar_bb energy term [ref, ref]. Probabilities for the *SugarBBSampler* are derived by taking the exponent of the negative energy [ e^(-Energy) ]of each dihedral angle from 0 to 360 degrees with a step size of .1 degree, dependent on the anomeric chemistry of the linkage. The energy used in this equation is obtained through direct evaluation of the energy term for a particular torsion angle.

The *SugarBBSampler* stores this data and uses these probabilities to choose a dihedral angle and set that particular torsion to that value. The other backbone movers; small, medium, and large, simply choose a random value within a set range (+/- 15, 30, 90 degrees respectively) and change the current torsion accordingly. These samplers are used at a 4:2:1 ratio, where the small sampler is 4 times more likely to run during sampling than the large mover.

**DOFs: Structure Minimization.** Finally, the *MinMover* handles full-structure minimization of the glycan tree and the *PackRotamersMover* handles optimization of the carbohydrate OH groups and constituents at 60 degree intervals, packing the rotamers of neighboring protein side-chain residues using the 2010 Dunbrack Rotamer Library[5] within 6A.

In order to speed up computations on many glycan trees or large glycans, both the *MinMover* and *PackRotamersMover* randomly choose a residue from the residues currently set to model and obtain a list of residues from that residue out to the end of the parent tree or current modeling layer using the implemented *RandomGlycanFoliageSelector.* This list is then used as the set of glycan residues to optimize during the application of these movers, with the *PackRotamersMover* additionally optimizing neighboring side-chain residues of this list within the 6 Å shell.

**DOFs: Shear Optimization.** A shearing motion is a movement that minimizes the downstream effect of a torsional change by making a counter-rotation that results in limiting downstream moves to small translations. For a geometrical shearing motion of downstream coordinates to occur, the two twisting bonds must be near parallel, and the bond twists must occur in opposite directions with equal magnitude. If the bonds are not near parallel, the downstream chain will spiral. In the case of peptides, because the omega angle is nearly always a trans peptide bond, the two bonds on either side, that is, $\psi_{n-1}$ and $\varphi_n$, are forced to be near parallel.

Traditionally, shear moves in Rosetta have simply made an equal but opposite twist to the $\psi_{n-1}$ and $\varphi_n$ of a peptide pose. In non-peptide cases, in the absence of a trans peptide bond, there is no guarantee of any particular bonds being near-parallel to each other, so functions were written to search for nearby bonds with similar directional/3D orientations. This modification to underlying code has permitted shearing moves to be made during sampling strategies of polysaccharide chains in Rosetta. Furthermore, special checks needed to be added to ensure that a pair of shearing torsional changes were not made across a branch point. Otherwise, a saccharide main chain might shear while its branch or branches twisted.

### Benchmarking

**Datasets.** The benchmarking dataset used for optimization comprises 25 individual glycan trees spanning 19 glycoprotein PDBs that were lower than 2 Å resolution, with most structures being less than 1.5 Å in resolution (Table S2). These glycan trees were unique in structure across the set, and all glycan residues were checked for any inconsistencies using the glycosciences.de pdb-care webtool[12] and checked for proper Rosetta input through the new *glycan_info* Rosetta application.

**Table S2: Structures used for this work.**

| PDB ID | Branch(es) | Length(s) | Resolution |
|---|---|---|---|
| 1f8d | 200A | 7 | 1.4 |
| 1gai | 171A,395A | 5,9 | 1.7 |
| 1jnd | 200A | 4 | 1.3 |
| 1juh | 191A,191B | 7,4 | 1.6 |
| 2ciw | 93A | 3 | 1.2 |
| 2cl2 | 43A | 7 | 1.4 |
| 3ave | 297A | 8 | 2.0 |

| | | | |
|---|---|---|---|
| 3gml | 42A,165A | 5,6 | 1.7 |
| 3nkq | 524A | 6 | 1.7 |
| 3og2 | 627A,930A | 7,12 | 1.2 |
| 3pxl | 54A,217A | 7,3 | 1.2 |
| 3pfx | 267A | 4 | 1.3 |
| 3qvr | 89A | 5 | 1.3 |
| 3uue | 253A | 5 | 1.5 |
| 4dgr | 200A | 9 | 1.6 |
| 4do4 | 124A,177A | 3,5 | 1.4 |
| 4f8x | 336A | 5 | 1.5 |
| 4nyq | 35A | 3 | 1.2 |
| 4q56 | 35A | 6 | 1.4 |

**Dataset preparation**. In order to model the glycan trees in the context of their crystal environment, *RosettaSymmetry*[13] was used to generate and replicate the surrounding crystal environment. Density files were in the CCP4 format and generated using the phenix.maps tool[14]. RosettaDensity[15] was used for density-building studies, crystallographic refinement, and the calculation of the density fit metric.

All input structures were refined into the Rosetta energy function using the FastRelax[16] mover with the generated crystal density used as structural constraints in a symmetric context. The crystal density term, elec_dens_fast, was set to a weight of 20. Waters were ignored by default. Inputs were refined separately for both Ref2015[17] and Rosetta-ICO (beta_nov16) studies[18]. For REF2015 refinement , the *fa_intra_rep_xover4* energy term (which scores atomic repulsion within residues and is part of beta_nov16) was enabled at a weight equal to the *fa_rep* energy term. The lowest energy model for each PDB was chosen from a set of 10 as the *exemplar* model. All glycan torsions, including the ASN linkage, were randomized upon input into the modeling algorithm for all benchmarking experiments for each parallel run. The refinement script can be found below:

```
<ROSETTASCRIPTS>
 <SCOREFXNS>
 </SCOREFXNS>
 <SIMPLE_METRICS>
  <RMSDMetric name="rmsd" use_native="1" rmsd_type="rmsd_all_heavy"/>
 </SIMPLE_METRICS>
 <MOVERS>
  <SetupForSymmetry name="setup_symm" definition="%%symmdef%%"/>
  <LoadDensityMap name="loaddens" mapfile="%%map%%"/>
  <SetupForDensityScoring name="setupdens"/>
  <FastRelax name="relax_dens" scorefxn="commandline" repeats="1" batch="false"     ramp_down_constraints="false" />
  <ExtractAsymmetricUnit name="extract_asymm" keep_virtual="0" />
 </MOVERS>
 <PROTOCOLS>
  <Add mover="setup_symm"/>
  <Add mover="loaddens"/>
  <Add mover="setupdens"/>
  <Add mover="relax_dens" />
  <Add mover="extract_asymm" />
 </PROTOCOLS>
 <OUTPUT scorefxn="commandline"/>
</ROSETTASCRIPTS>
```

**Benchmarking Specifics.** Benchmarking was run on a 500 processor MPI compute cluster. The *rosetta_scripts_jd3* application (described below) was used to run each set of experiments. A Job Definition file defined options for each experiment, while an associated RosettaScript XML defined the protocol and metrics. A total of 1500 decoys were created for each of the input glycans for each experiment. For the final *de-novo* benchmark, 5000 decoys were created for each input glycan, while 1500 were used for the density build benchmark. The scoreterm *elec_dens_fast* was used at a weight of 25 for density building. Both final benchmarks used the Rosetta-ICO (beta) scorefunction, and a *sugar_bb* weight of 0.5 (which is now the default when working with glycans).

In general, the *GlycanTreeModeler* was used to model glycans, while the *GlycanResidueSelector* was used to specify the particular glycan tree being modeled. All glycans were modeled in their symmetric crystal environment unless otherwise noted. Analysis was done using the *SimpleMetric* framework and plots were created using the python packages matplotlib[19] pandas[20] , and seaborn[21]. Figures were created using Inkscape.

### Optimization Figures

*Kinematic Optimization*

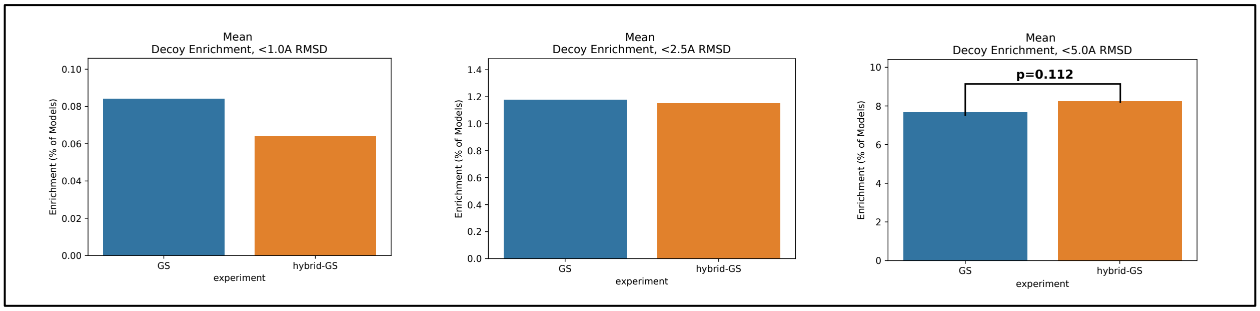


**Figure S3: Enrichments of GlycanSampler(GS) compared to GlycanTreeModeler and then GlycanSampler (hybrid-GS).**

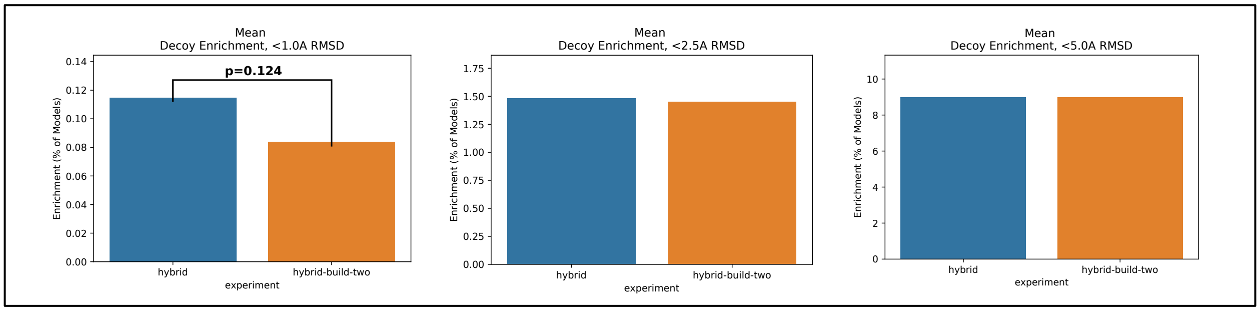


**Figure S4: Hybrid Enrichments compared to Hybrid building two layers at a time.**

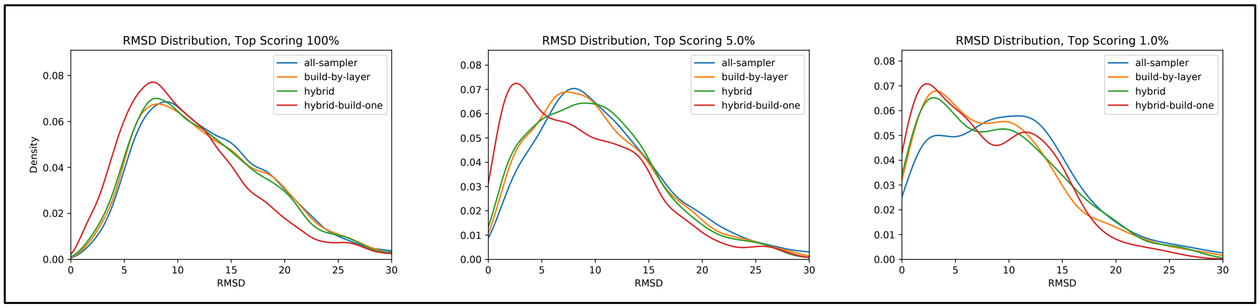


**Figure S5: Overall pool of models at varying filters of total score.** N=150,000; 37,500 per experiment.

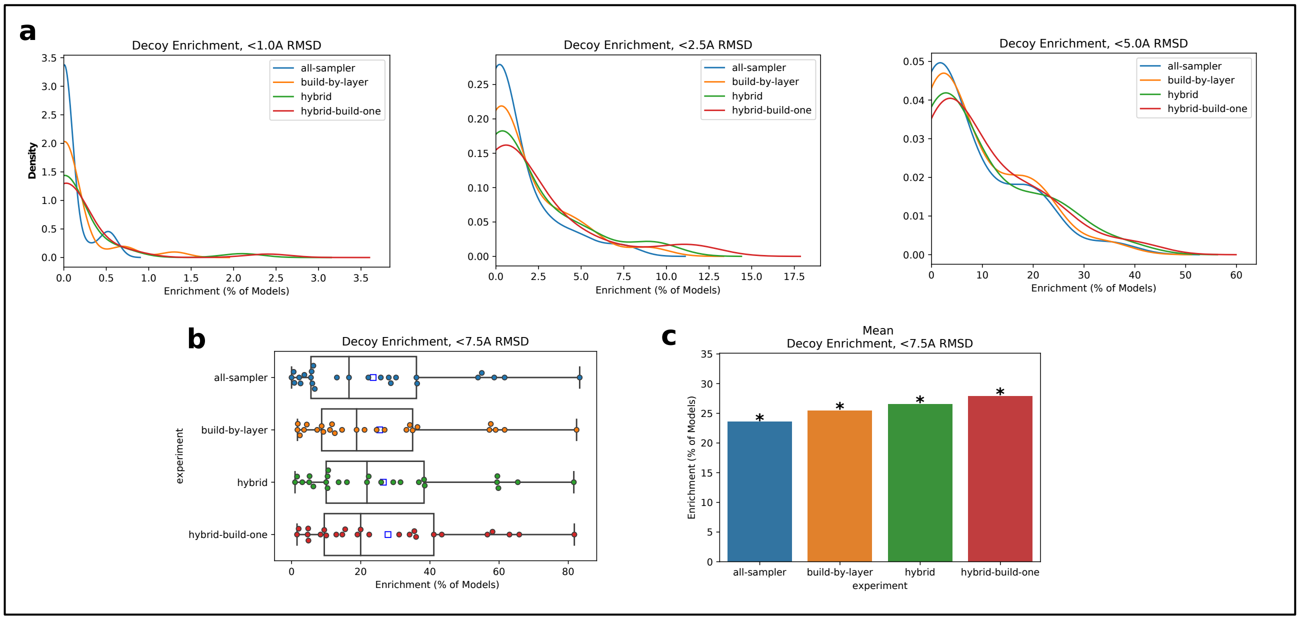


**Figure S6: Kinematic Experiments – Enrichment KDEs and 7.5A.** A. Kernel Density Estimates of enrichment per input model for each major kinematic experiment. B. Box plot of enrichment of each input model per experiment less than 7.5 A RMSD to the native crystal structure. C. Means of B, with paired t-test, All *vs.* All. * indicates $p<.05$. p-value for hybrid-build-one *vs.* build-by-layer $p< .005$, while *vs.* all-sampler $p<.0005$.

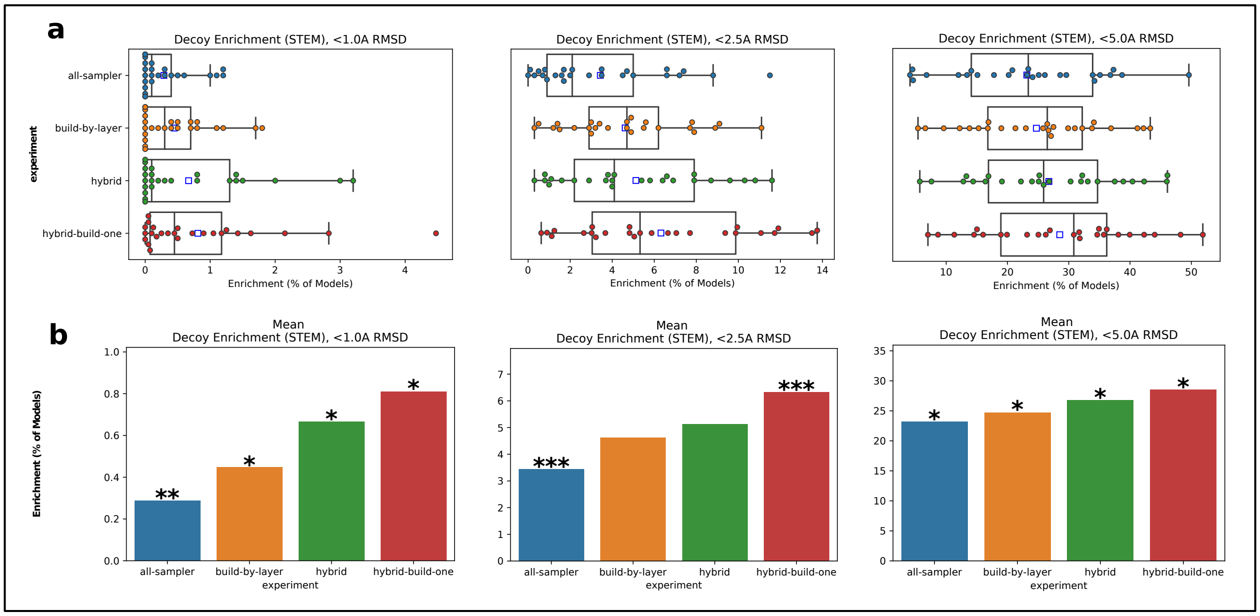


**Figure S7: Kinematic Experiments - Decoy Enrichment of STEM (Layers 0 and 1). a.** Boxplots of each input glycan benchmark at <1.0A, <2.5A, and < 5.0A of the glycan STEM **b.** Means of fig A. Asterisk above bar indicate statistical significance with all other groups through paired t-test. *|p <.05; **|p<.005; ***|p<.0005 . For b <1A, pvalue of all-sampler *vs.* hybrid-build-one is **.

*Scoring Optimization*

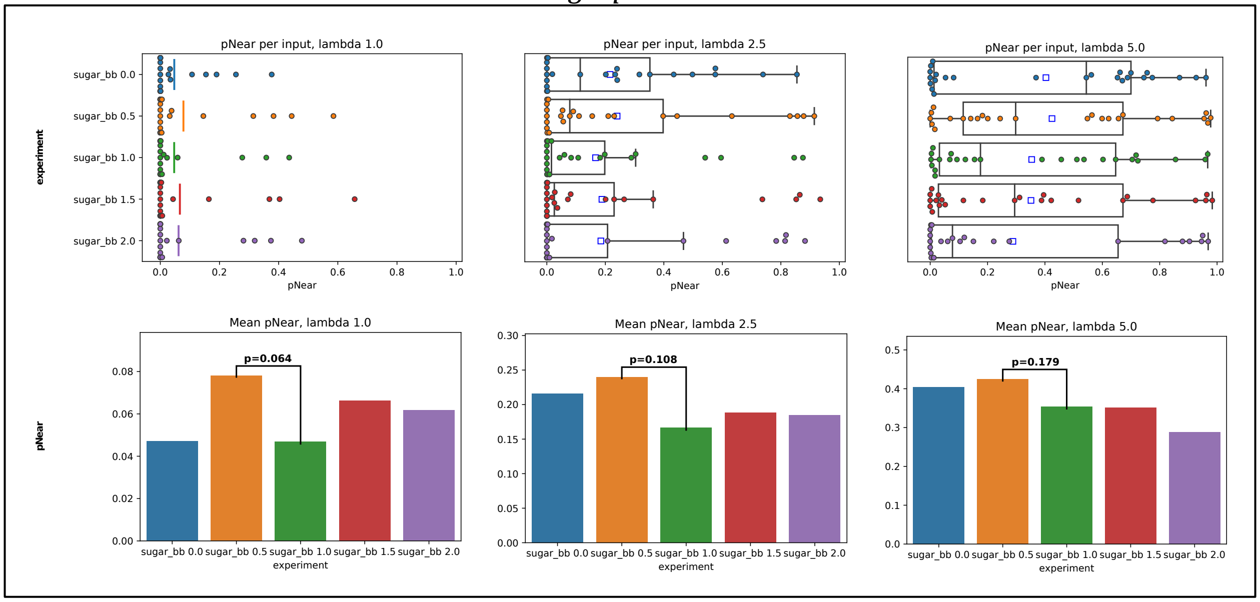


**Figure S8: Scoring Experiments – Decoy Discrimination of various values of sugar_bb energy term.** 1000 decoys were produced for each glycan and each experiment for a total of 125k decoys. Note that this is a third less than all other optimization experiments. **A.** Boxplots of PNear metric at various lambdas. Blue square indicates mean. Line in box indicates median. **B.** Bar plots of PNear for each significant lambda. Paired T-test results between *sugar_bb* weight of 1.0 and .5 are shown. All other comparisons are much worse and are not shown.

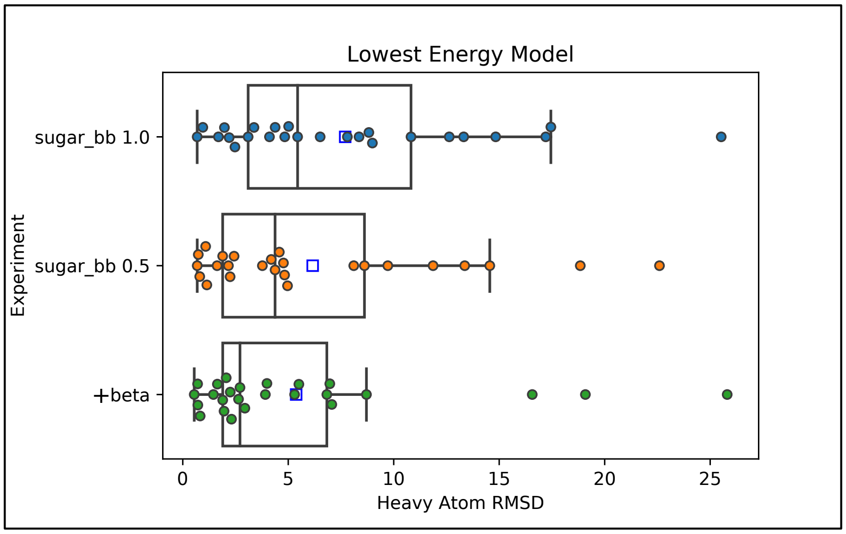


**Figure S9: Boxplot of the lowest energy model for all major scoring experiments across all benchmark glycans**

### De novo Benchmarking

**Table S3: Raw *de novo* Modeling results for each glycan tree**

| pdb branch, size | layer01_rmsd | rmsd | super_rmsd | fit6_rmsd | fit6_super_rmsd |
|---|---|---|---|---|---|
| 3pxl 217A, 3 | 0.17 | 0.54 | 0.45 | 0.54 | 0.45 |
| 4do4 177A, 5 | 0.65 | 1.72 | 1.59 | 0.70 | 0.77 |
| 2ciw 93A, 3 | 0.41 | 0.71 | 0.43 | 0.71 | 0.43 |
| 4nyq 35A, 3 | 0.69 | 0.72 | 0.67 | 0.72 | 0.67 |
| 1jnd 200A, 4 | 0.25 | 0.95 | 0.74 | 0.95 | 0.74 |
| 3gml 42A, 5 | 0.72 | 2.15 | 1.35 | 1.54 | 1.26 |
| 4dgr 200A, 9 | 0.61 | 1.96 | 1.54 | 1.96 | 1.54 |
| 3qvr 89A, 5 | 1.40 | 2.07 | 1.70 | 2.07 | 1.70 |
| 4f8x 336A, 5 | 0.74 | 2.18 | 1.79 | 2.08 | 1.80 |
| 1juh 191B, 4 | 1.99 | 1.94 | 0.42 | 2.08 | 0.40 |
| 4do4 124A, 3 | 2.52 | 3.50 | 0.47 | 2.52 | 0.31 |
| 3og2 930A, 12 | 0.63 | 2.82 | 2.36 | 2.54 | 2.14 |
| 4q56 35A, 6 | 2.14 | 2.67 | 0.71 | 2.67 | 0.71 |
| 3gml 165A, 6 | 1.28 | 3.58 | 1.94 | 3.58 | 1.94 |
| 1gai 171A, 5 | 0.99 | 3.71 | 2.48 | 3.71 | 2.48 |
| 1f8d 200A, 7 | 1.40 | 4.20 | 1.59 | 4.20 | 1.59 |
| 3og2 627A, 7 | 1.79 | 4.37 | 2.81 | 4.37 | 2.81 |
| 1juh 191A, 7 | 3.56 | 4.69 | 3.11 | 4.69 | 3.11 |

| | | | | | |
|---|---|---|---|---|---|
| 3pfx 267A, 4 | 2.89 | 6.16 | 0.97 | 6.16 | 0.97 |
| 1gai 395A, 9 | 0.89 | 6.59 | 5.54 | 6.59 | 5.54 |
| 3nkq 524A, 6 | 1.32 | 8.12 | 4.30 | 8.12 | 4.30 |
| 3uue 253A, 5 | 0.36 | 10.24 | 4.71 | 10.24 | 4.71 |
| 3pxl 54A, 7 | 4.86 | 11.71 | 2.24 | 11.71 | 2.24 |
| 2cl2 43A, 7 | 8.15 | 14.97 | 6.95 | 14.97 | 6.95 |
| 3ave 297A, 8 | 13.93 | 24.94 | 3.63 | 24.94 | 3.63 |

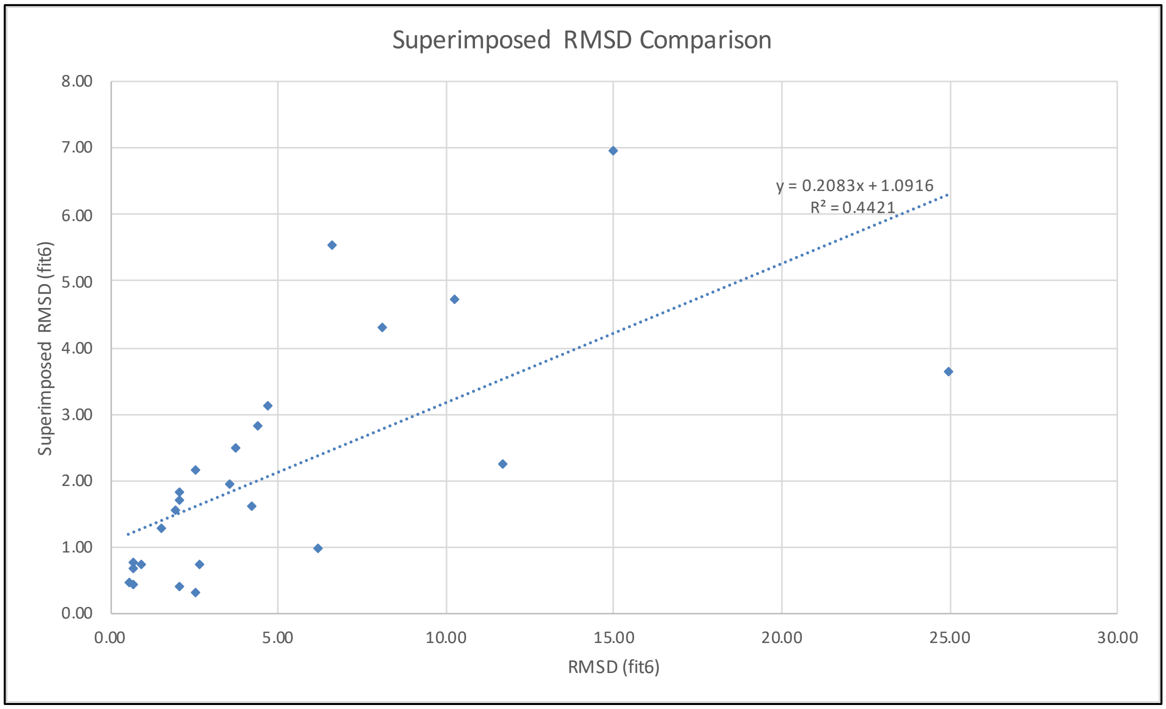


**Figure S10:** Superimposed RMSD comparisons of the top scoring model for each de novo modeled glycan tree.

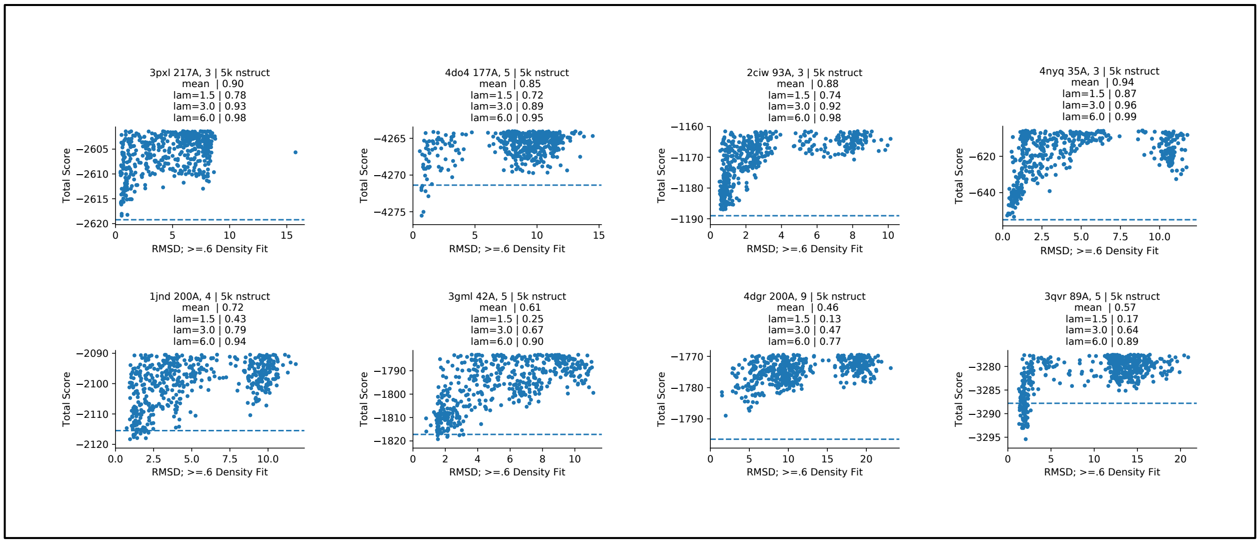

**Figure S11: Score vs. RMSD funnel plots of the best predicted glycan structures with pNear at different lambda values. Shown is the top 10% of models by total energy. Blue line is the scored native structure with symmetry.**

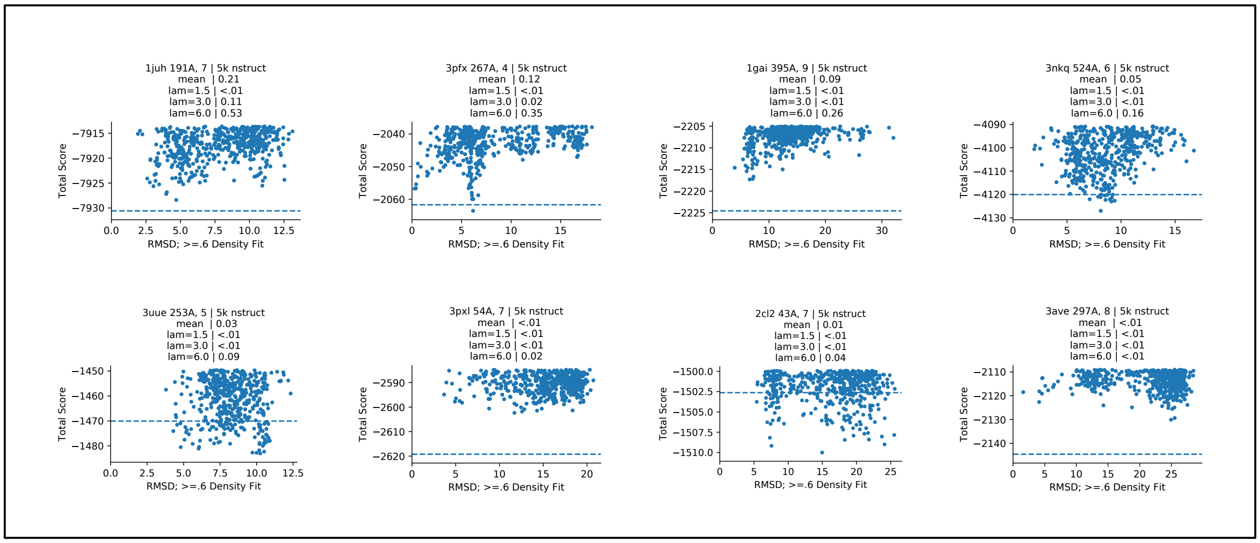


**Figure S12: Score vs. RMSD funnel plots of the worst predicted glycan structures with pNear at different lambda values. Shown is the top 10% of models by total energy. Blue line is the scored native structure with symmetry.**

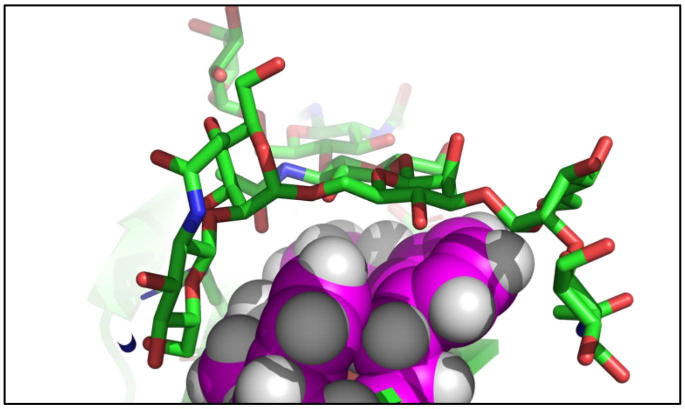

**Figure S13: Hydrophobic surface interactions with 3ave glycan at residue 297, chain A.** F241, 243F, 262V, and 264V are shown as spheres at the glycan interface.

**Table S4: Rosetta-ICO vs Rosetta-ECO mean pNear values at various lambdas. N=8**

| lambda | Implicit Solvent | sd | Explicit Solvent | sd |
|---|---|---|---|---|
| 1.5 | 0.225 | 0.303 | 0.005 | 0.012 |
| 3 | 0.400 | 0.371 | 0.051 | 0.128 |
| 6 | 0.568 | 0.409 | 0.207 | 0.256 |

**Table S5: Density-guided modeling results for each glycan tree**

| pdb-branch-low-size | layer01_rmsd | rmsd | super_rmsd | fit6_rmsd | fit6_super_rmsd |
|---|---|---|---|---|---|
| 3gml 165A, 6 | 0.10 | 0.09 | 0.08 | 0.09 | 0.08 |
| 3og2 627A, 7 | 0.15 | 0.10 | 0.08 | 0.10 | 0.08 |
| 4nyq 35A, 3 | 0.09 | 0.11 | 0.10 | 0.11 | 0.10 |
| 2cl2 43A, 7 | 0.17 | 0.14 | 0.12 | 0.14 | 0.12 |
| 3uue 253A, 5 | 0.20 | 0.16 | 0.13 | 0.16 | 0.13 |
| 1juh 191B, 4 | 0.18 | 0.37 | 0.29 | 0.17 | 0.20 |
| 3pxl 217A, 3 | 0.18 | 0.21 | 0.18 | 0.21 | 0.18 |
| 1f8d 200A, 7 | 0.36 | 0.22 | 0.20 | 0.22 | 0.20 |
| 4dgr 200A, 9 | 0.36 | 0.22 | 0.21 | 0.22 | 0.21 |
| 1gai 395A, 9 | 0.18 | 0.22 | 0.21 | 0.22 | 0.21 |
| 3gml 42A, 5 | 0.21 | 0.63 | 0.56 | 0.22 | 0.21 |
| 1juh 191A, 7 | 0.22 | 0.27 | 0.23 | 0.27 | 0.23 |
| 3og2 930A, 12 | 0.13 | 0.31 | 0.31 | 0.28 | 0.27 |
| 3pxl 54A, 7 | 0.16 | 0.29 | 0.27 | 0.29 | 0.27 |
| 3nkq 524A, 6 | 0.19 | 0.31 | 0.30 | 0.31 | 0.30 |
| 4q56 35A, 6 | 0.33 | 0.36 | 0.34 | 0.36 | 0.34 |
| 3pfx 267A, 4 | 0.13 | 0.36 | 0.35 | 0.36 | 0.35 |
| 4do4 177A, 5 | 0.25 | 2.43 | 2.03 | 0.42 | 1.01 |
| 3ave 297A, 8 | 0.23 | 0.46 | 0.39 | 0.46 | 0.39 |
| 4do4 124A, 3 | 0.46 | 0.53 | 0.39 | 0.46 | 0.25 |
| 4f8x 336A, 5 | 0.07 | 1.34 | 1.24 | 0.52 | 0.73 |
| 2ciw 93A, 3 | 0.16 | 0.55 | 0.47 | 0.55 | 0.47 |
| 3qvr 89A, 5 | 1.02 | 0.70 | 0.65 | 0.70 | 0.65 |
| 1jnd 200A, 4 | 0.10 | 0.79 | 0.62 | 0.79 | 0.62 |
| 1gai 171A, 5 | 0.14 | 0.89 | 0.84 | 0.89 | 0.84 |

### Glycan Masking

**Computational design of N-linked sequons into solvent-exposed protein surfaces.** First, all possible residues on the outward facing surfaces of I53-50A trimers[22] when assembled into nanoparticles were manually selected as candidate locations for designing in a NxT sequon for N-linked glycan placement. Next, the RosettaScripts protocol and .sh file below were used to sequentially knock-in a single NxT sequon at these selected locations and obtain calculated energies of the new protein structure using the Rosetta score function. The *CreateGlycanSequonMover* was used to design in these sequons. Both typical and enhanced[23,24] sequons were attempted at each position . The resulting computational outputs consisted of a .pdb file and a .sc file (a "score" file that lists various Rosetta score function outputs) for each new protein that had a single NxT sequon added to it. Protein structures were first scored by Rosetta without a model glycan tree present to eliminate any potential interference of the glycan atoms. To filter out bad designs, outputs with a "total_energy" of >500 and a RMSD >0.45 Å compared to the original I53-50A scaffold were discarded. The re-designed protein structures that passed this filtering step were then glycosylated using the *SimpleGlycosylateMover* with a model tri-antennary man9 N-linked glycan, modeled using the *GlycanTreeModeler*, and finally scored by Rosetta. A second round of filtering was performed using the same criteria as above. After proteins with a single sequon were experimentally screened for expression and glycan occupancy (see below), combinations of sequons were designed into the outer surface of I53-50A proceeded through the same computation pipeline described above with the experimentally screened glycan sites as the lead sequon for combinations. The XML file for this combinatorial selection is shown below.

**Plasmid construction.** For each protein design that resulted from the above computational pipeline, the final construct contained a N-terminal secretion signal sequence derived from the modified bovine prolactin (MDSKGSSQKGSRLLLLLVVSNLLLPQGVLA) and C-terminal myc and hexa-histidine tags (LEEQKLISEEDLHHHHHH). These constructs were then cloned by GenScript into the pCMV/R plasmid using the restriction sites Xba1 and AvrII.

**Small-scale screening of proteins with computationally designed sequons.** Small-scale 2.0 mL cultures of Expi293F cells were grown in suspension to a density of $3.0 \times 10^6$ cells per mL and transiently transfected using PEI-MAX (Polyscience) and cultivated for 5 days in Expi293F expression medium (Life Technologies) at 37ºC, 70% humidity, 8% $CO_2$, and rotating at 150 rpm. Supernatants were clarified by centrifugation (5 min at 4000 rcf), PDADMAC solution was added to a final concentration of 0.0375% (Sigma Aldrich, #409014), and a final spin was performed (5 min at 4000 rcf). Supernatants were concentrated using a 5 kDa MWCO spin filter (Sartorius) to a final volume of ~50 µL. These concentrated supernatants were then assessed for protein expression by Western blot using an anti-myc mouse primary antibody and an anti-mouse HRP-conjugated goat secondary antibody. Glycan occupancy for each protein design was assessed by comparing SDS-PAGE gels of untreated and PNGaseF-treated (NEB) protein.

**Large-scale expression and purification of glycosylated protein.** For large-scale protein expression, 800 mL cultures of Expi293F cells were transiently transfected and cultivated for 5 days as described above. Proteins were purified from clarified supernatant via a batch bind method where Talon cobalt affinity resin (Takara) was added to supernatants and allowed to incubate for 15 min with gentle shaking. Resin was isolated using 0.2 µm vacuum filtration and transferred to a gravity column, where it was washed with 20 mM Tris pH 8.0, 300 mM NaCl, and protein was eluted with 3 column volumes of 20 mM Tris pH 8.0, 300 mM NaCl, 300 mM imidazole. This batch bind process was repeated a second time on the supernatant flow-through from the filtration step. Eluate with protein was concentrated to ~2 mL using a 30 kDa MWCO Amicon concentrator (Millipore Sigma).

The concentrated sample was sterile filtered (0.2 μm) and applied to a Superdex 200 Increase 10/300 SEC column (Cytiva) using 25 mM Tris pH 8.0, 150 mM NaCl, 0.75% CHAPS, 5% glycerol buffer.

***In vitro* nanoparticle assembly and purification.** The protein concentration of individual nanoparticle components (I53-50A trimer and I53-50B.4PT1 pentamer) was determined by measuring 280 nm absorbance using a UV/vis spectrophotometer (Agilent Cary 8454) and estimated extinction coefficients[22]. Particle assembly was performed by adding equimolar amounts of I53-50A and I53-50B to reach a final protein concentration of 20 μM (10 μM for each individual component) and resting on ice for at least 30 min. Assembled particles were sterile filtered (0.2 μm) immediately before SEC purification using a Superose 6 Increase 10/300 GL column to remove residual unassembled component.

**Dynamic light scattering.** Dynamic light scattering (DLS) was used to measure the hydrodynamic diameter of I53-50 nanoparticles with either I53-50A non-glycosylated trimers or I53-50A glycosylated trimers on a DynaPro NanoStar instrument (Wyatt Technologies). 2 μL of 0.1 mg/mL protein was applied to a quartz cuvette to obtain intensity measurements from 10 acquisitions of 10 s each. Increased viscosity due to 5% glycerol in the buffer was accounted for by the software.

**Mouse immunization.** Four-week-old female BALB/c mice (Jackson Laboratory, Stock: 000651) were purchased and maintained at the Comparative Medicine Facility at the University of Washington, Seattle, WA, which is accredited by the American Association for the Accreditation of Laboratory Animal Care International (AAALAC). At six weeks of age, mice were inoculated with 5.57 μg bare or glycosylated I53-50 particles and again at 3 and 6 weeks later. Prior to inoculation, immunogen suspensions were gently mixed 1:1 vol/vol with AddaVax adjuvant (Invivogen, San Diego, CA). Mice were injected intramuscularly into the gastrocnemius muscle of each hind leg using a 27-gauge needle with 50 μL per injection site (100 μL total) of immunogen under isoflurane anesthesia. For sera collection, mice were bled via submental venous puncture 2 weeks following each inoculation. Serum was isolated from hematocrit via centrifugation at 2,000 g for 10 min, and stored at -80ºC until use.

**ELISA**. First, 50 μL of 2.0 μg/mL I53-50A or I53-50A(gly) trimer per well was incubated for 1 hr in 96-well Nunc MaxiSorp plates (Thermo Scientific). Then 200 μL of TBST buffer (25 mM Tris pH 8.0, 150 mM NaCl, 0.05% v/v Tween20) with 2% w/v BSA was added to each well and incubated 1 hr incubation. Plates were washed three times using a robotic plate washer (BioTek) with TBST. Then 50 μL of serum dilutions starting at 1:100 and serially diluting 5-fold seven times using TBST with 2% w/v BSA (8 total dilutions) were added to each well and incubated for 1 hr. After washing plates 3x with TBST, 50 μL of anti-mouse HRP-conjugated goat secondary antibody (CellSignaling Technology) diluted 1:5,000 in TBST with 2% w/v BSA incubated in each well for 1 hr. Following a final 3x TBST plate wash, 100 μL of TMB was added to each well and rested for 2.0 min, then 100 μL of 1.0 M HCl was added to each well to quench the reaction. Absorbance at 450 nm was immediately collected for each well on a SpectraMax M5 plate reader (Molecular Devices). Data were plotted and fit in Prism (GraphPad) using nonlinear regression sigmoidal, 4PL, X is concentration, to determine $EC_{50}$ values from curve fits. All steps were performed at ambient temperature.

**.xml file**:

```
<ROSETTASCRIPTS>
  <SCOREFXNS>
        <ScoreFunction name="sfx_clean" weights="beta" symmetric="0" /> //function to obtain a
score
  </SCOREFXNS>
  <RESIDUE_SELECTORS>
        <Index name="select_i_enh0" resnums="%%resi_enh0%%" /> //select the residue(s) to
glycosylate (these residues are a "non-enhanced" sequon)
               Not name="not_resis" selector="select" /> //all other residues not selected
        <Index name="select_i_enh1" resnums="%%resi_enh1%%" /> //select the residue(s) to
glycosylate (these residues are an "enhanced" sequon)
               Not name="not_resis" selector="select" /> //all other residues not selected
        Index name="select_i-2" resnums="%%enhresi%%" /> //select the residue i-2 from the N-
linked glycosylation site
        Index name="select_i_score" resnums="%%iresi%%" /> //select Asn residue i that is N-liked
glycosylated to get its score
  </RESIDUE_SELECTORS>
  <FILTERS>
        EnergyPerResidue name="total_energy_per_res_filter_i" scorefxn="sfx_clean"
energy_cutoff="10000" resnums="%%iresi%%" /> // tests the energy of a particular residue, or
interface, or whole protein, or a set of residues; energy must be less than 10000
        EnergyPerResidue name="total_energy_per_res_filter_i-2" scorefxn="sfx_clean"
energy_cutoff="10000" resnums="%%enhresi%%" /> // tests the energy of a particular residue, or
interface, or whole protein, or a set of residues; energy must be less than 10000
        EnergyPerResidue name="fa_atr_per_res_filter" scorefxn="sfx_clean" score_type="fa_atr"
energy_cutoff="10000" resnums="%%resi%%" />
        EnergyPerResidue name="fa_rep_per_res_filter" scorefxn="sfx_clean" score_type="fa_rep"
energy_cutoff="10000" resnums="%%resi%%" />
        EnergyPerResidue name="fa_dun_per_res_filter" scorefxn="sfx_clean" score_type="fa_dun"
energy_cutoff="10000" resnums="%%resi%%" />
        EnergyPerResidue name="fa_elec_per_res_filter" scorefxn="sfx_clean" score_type="fa_elec"
energy_cutoff="10000" resnums="%%resi%%" />
  </FILTERS>

  <MOVERS>
        <CreateGlycanSequonMover name="create_motif_enh0" residue_selector="select_i_enh0"
basic_enhanced_n_sequon="0" design_x_positions="1" pack_neighbors="1" scorefxn="sfx_clean" />
        <CreateGlycanSequonMover name="create_motif_enh1" residue_selector="select_i_enh1"
basic_enhanced_n_sequon="1" design_x_positions="1" pack_neighbors="1" scorefxn="sfx_clean" />
        <SimpleGlycosylateMover name="glycosylate_enh0" residue_selector="select_i_enh0"
glycosylation="a-D-Manp-(1->3)-[a-D-Manp-(1->3)-[a-D-Manp-(1->6)]-a-D-Manp-(1->6)]-[b-d-GlcpNAc-
(1->4)]-b-D-Manp-(1->4)-b-D-GlcpNAc-(1->4)-[a-L-Fucp-(1->6)]-b-D-GlcpNAc-" strip_existing="1" />
        <SimpleGlycosylateMover name="glycosylate_enh1" residue_selector="select_i_enh1"
glycosylation="a-D-Manp-(1->3)-[a-D-Manp-(1->3)-[a-D-Manp-(1->6)]-a-D-Manp-(1->6)]-[b-d-GlcpNAc-
(1->4)]-b-D-Manp-(1->4)-b-D-GlcpNAc-(1->4)-[a-L-Fucp-(1->6)]-b-D-GlcpNAc-" strip_existing="1" />
        <SymMinMover name="bb_min" scorefxn="sfx_clean" bb="1" chi="1" jump="0"
type="lbfgs_armijo_nonmonotone" tolerance="0.005" max_iter="100" />
        <GlycanTreeModeler name="tree_modeler" quench_mode="false" rounds="1" layer_size="1"
window_size="0" hybrid_protocol="1" shear="1" use_gaussian_sampling="1"
glycan_sampler_rounds="150" />
  </MOVERS>

  <PROTOCOLS>
        // wiggle backbone to loosen up a bit
        <Add mover_name="bb_min" />

        // generate sequon (enhanced or not) and add glycan
        <Add mover_name="create_motif_enh0" />
        <Add mover_name="create_motif_enh1" />
```

```
        // wiggle backbone to loosen up a bit
        <Add mover_name="bb_min" />

        // filter to extract energy of residues
        Add filter_name="total_energy_per_res_filter_i" />
        Add filter_name="total_energy_per_res_filter_i-2" />
        Add filter_name="fa_atr_per_res_filter" />
        Add filter_name="fa_rep_per_res_filter" />
        Add filter_name="fa_dun_per_res_filter" />
        Add filter_name="fa_elec_per_res_filter" />

        // add glycan and model glycan
        <Add mover_name="glycosylate_enh0" />
        <Add mover_name="glycosylate_enh1" />
        <Add mover_name="tree_modeler" />
  </PROTOCOLS>
</ROSETTASCRIPTS>
```

**.sh file**:

```
#input arguments
scaffold=$1 ; resi_enh0=$2 ; resi_enh1=$3 ; glycansites=$4 ; enhresi=$5 ; iresi=$6
#("glycansites" lists the combined residues in "resi_enh0" and "resi_enh1", but is underscore-
separated, NOT comma-separated; this is to open in PyMol, which doesn't like commas in the
filename)
#"enhanced" is the on/off switch for the sequon mover to make an "enhanced sequon" by adding an
aromatic at the i-2 position (this variable was removed on 9-5-2019 because for combinations we
need the ability to make sequon combinations where some are "enhanced" and others are not)
#"enhresi" is the 'enhanced' residue that is i-2, where i is the N-linked glycosylation residue

if [ ! -e output/ ]; then mkdir output/; fi
outpath="output/"

#symfile="symdef/I/I53.sym"
#symdof1="JCP00"; symdof2="JCT00"

/software/rosetta/latest/bin/rosetta_scripts.hdf5.linuxgccrelease \
        -parser:protocol xml/Glycosylate.xml \
        -include_sugars \
        -alternate_3_letter_codes pdb_sugar \
        -auto_detect_glycan_connections \
        -min_bond_length 1.1 \
        -max_bond_length 1.7 \
        -ignore_zero_occupancy false \
        -ignore_unrecognized_res \
        -parser:script_vars outpath="$outpath" resi_enh0="$resi_enh0" resi_enh1="$resi_enh1"
glycansites="$glycansites" \
        -overwrite \
        -unmute all \
        -unmute protocols.rosetta_scripts \
        -out:suffix "_${glycansites}" \
        -out::path::all ${outpath} > ${outpath}/"${glycansites}".log \
        -s input/scaffolds/${scaffold}.pdb \
        -native input/scaffolds/${scaffold}.pdb \
        -beta
```

### Rosetta Functionality Extensions

**SimpleMetrics.** Rosetta as a software suite has been used successfully for modeling and designing biomolecules, but rigorous analysis of results is typically done in other programs with some notable exceptions. The *interface_analyzer* application is widely used for analysis of Protein-Protein interfaces and has recently been updated to allow its use in *RosettaScripts*[25]. The Rosetta *FeaturesReporter* system was one of the first developed analysis frameworks in Rosetta and can be used to output sets of data into databases[26]. This framework can be extremely useful; however, it requires expensive database licenses to output results concurrently in a streamlined fashion, it can be difficult for a developer to create new *FeaturesReporter* classes, and the databases themselves can easily become unmanageable in size. Finally, the Rosetta filter system has been used to do some metric calculation for single numerical values, but a simple and more robust metric interface was still needed.

The *SimpleMetric* system complements existing methods of data analysis by streamlining how analysis is done. A *SimpleMetric* calculates a single value type (numeric or string), stores it within the Pose, and then adds the data to the resulting Rosetta Data File (scorefile) at the end of each run of the protocol, typically for each output decoy. There are three major types of *SimpleMetrics*: normal metrics which calculate a single value, *CompositeMetrics* that calculate multiple named values, and *PerResidueMetrics* which calculate values for each residue specified by a Residue Selector.

*SimpleMetrics* are available through the C++ and PyRosetta interfaces, but have been tailored especially for *RosettaScripts* through a new section in the *RosettaScript* XML. To run these metrics at any arbitrary point in a protocol, the *RunSimpleMetricsMover* takes a list of specific *SimpleMetrics* created in the new section, with an optional prefix and suffix, runs the metrics, and stores the data in the pose for further Rosetta use and ultimate scorefile output. In conjunction with optional JSON scorefile output (option *-scorefile_format json*), analysis can be further simplified using the python programming language with the pandas module for simple DataFrame creation and plotting.

In order to increase the utility of the *SimpleMetric* system, a *SimpleMetricFilter* can be used, which takes an arbitrary metric and runs them as a filter. This data can be calculated at the time of the filtering or the cached metric calculation can be used, saving run time for hefty metrics.

Finally, the *SimpleMetric* system is fully compatible with the Rosetta *FeaturesReporter* with the *SimpleMetricFeaturesReporter* in order to enable analysis through databases and backwards-compatibility.

A number of general *SimpleMetrics* have been written and used for this work including metrics for outputting **R**oot **M**ean **S**quare **D**eviation (RMSD), **S**olvent **A**ccessible **S**urface **A**rea (SASA), dihedral distance, sequence, secondary structure, and hydrogen bonds (Table S4). The SimpleMetric framework was an instrumental tool in our glycan benchmarking and should prove to be an important asset for the future of Rosetta's protocol development and scientific research.

### Table S6: Initial SimpleMetrics created and used in this work

| SimpleMetric | Description | Type |
|---|---|---|
| DihedralDistanceMetric | Calculates the normalized dihedral angle distance in degrees from directional statistics on a set of dihedrals/residues of two poses or two regions of a pose. | *RealMetric* |
| InteractionEnergyMetric | Calculates the (long range and short range) interaction energy between a selection and all other residues or another selection. Can be set to only calculate short or long or only use certain score terms such as *fa_rep*. | *RealMetric* |
| ResidueSummaryMetric | A metric that takes a *PerResidueRealMetric* and summarizes the data in different ways, such as the sum, mean, or the number of residues that match a certain criteria. Can use cached data. | *RealMetric* |
| RMSDMetric | Calculates the RMSD between two poses or on a subset of residues. Many options for RMSD including bb, heavy, all, etc. | *RealMetric* |
| SasaMetric | Calculates the Solvent Accessible Surface Area (SASA). | *RealMetric* |
| SelectedResidueCountMetric | Count the number of residues in a selection (or whole pose). | *RealMetric* |
| TotalEnergyMetric | Calculates the Total Energy of a pose using a Scorefunction OR the delta total energy between two poses. | *RealMetric* |
| TimingProfileMetric | Calculates the time passed in minutes or hours from from construction to apply (i.e. from when declared in the RS block to when it is run). Useful for obtaining timing information of protocols. | *RealMetric* |
| SecondaryStructureMetric | Returns the DSSP secondary structure of the pose or set of selected residues. | *StringMetric* |
| SelectedResiduesMetric | Returns a comma-separated list of selected residues in PDB or Rosetta numbering. | *StringMetric* |
| SelectedResiduesPyMOLMetric | Returns a PyMOL selection of a set of selected residues. | *StringMetric* |
| SequenceMetric | Returns the one or three-letter sequence of the pose or set of selected residues. | *StringMetric* |
| HbondMetric | Calculate number of hydrogen bonds of residues in a selector or between two selectors | *PerResidueRealMetric* |
| PerResidueDensityFitMetric | Calculate the Fit of a model to the loaded density either by Correlation or a Zscore. | *PerResidueRealMetric* |
| PerResidueClashMetric | Calculates the number of atomic clashes per residue using two residue selectors. Clashes are calculated through the Leonard Jones radius of each atom type. | *PerResidueRealMetric* |
| PerResidueEnergyMetric | Calculate any energy term for each residue. Total energy is default. If a native or repose is given, can calculate the energy delta for each residue. | *PerResidueRealMetric* |
| PerResidueRMSDMetric | Calculate the RMSD for each residue between the input and either the native or a reference pose. | *PerResidueRealMetric* |
| PerResidueSasaMetric | Calculate the Solvent Accessible Surface Area (SASA) of each residue. | *PerResidueRealMetric* |
| WaterMediatedHbondMetric | A metric to measure hydrogen bonds between a set of residues that are water-mediated (bridged). Can calculate different depths to traverse complex Hbond networks. | *PerResidueRealMetric* |

| | | |
|---|---|---|
| CompositeEnergyMetric | Calculates each individual scoreterm of a scorefunction or the DELTA of each scoreterm between two poses. Each named value is the scoreterm | *CompositeRealMetric* |
| ProtocolSettingsMetric | Outputs currently set user options (cmd-line, xml, or both). Allows one to only output specific metrics or set a tag for the particular experiment. Useful for benchmarking/plotting or historical preservation of options tied to a pose | *CompositeStringMetric* |

**RosettaScripts JD3.** A known limitation of the original *RosettaScripts* application is that only a single *RosettaScript* and associated configuration can be run at a time. For production runs of a single design or modeling task, this is adequate, but for benchmarking of multiple experimental configurations or for input structures that require associated input files, this can be problematic, as multiple runs of Rosetta would need to be scripted together in order to attain results. If this was run on a compute cluster, each run of Rosetta would use all the cores given and only spin down all the cores when the full job was finished. In order to improve benchmarking efficiency and to simplify the programming workflow for benchmarking tasks, a more streamlined *RosettaScripts* application was developed.

This application uses a new job distribution system that requires MPI and specific serialization routines for node to node communication and data transfer. This job distribution system is called JD3, with JD2 being the job distribution system used by the majority of Rosetta3 applications.

RosettaScripts JD3 uses a new file called a *Job Definition* to configure each independent job. This file is also an XML file, like *RosettaScripts*, and each job can be configured with different *RosettaScripts* XML files, substitution variables (*script_vars*), and command-line options. Combined with the *ProtocolSettings* SimpleMetric, benchmarks for each experiment and glycan tree can be run within a single, efficient cluster run. For our benchmarking, this file was created through a python script that specified each input PDB, glycan tree, and the path and name of each input file (symmetry definition, .ccp4 density file).

A general version of this script can be found in the open-source Jade2 repository (https://github.com/jadolfbr/jade2/blob/master/apps/pilot/jadolfbr/substitute_job_definition_with_pdbs.py) , while the script used for this work is included in the supplemental for reproducibility purposes.

***RosettaCarbohydrate* and General Rosetta Extensions.** During the creation of the RosettaScripts JD3 app, we made RosettaScripts able to be called within PyRosetta itself. This has already led to an increase in the utility of PyRosetta, as it can be much simpler to work in RosettaScripts versus PyRosetta, due to the complexity of some core components. A PyRosetta notebook for this additional functionality can be found here: https://github.com/RosettaCommons/PyRosetta.notebooks/blob/master/notebooks/02.07-RosettaScripts-in-PyRosetta.ipynb

Many components were created and used for this work. The following table includes new major Rosetta components and a short description of each of them. Most of these are available as *RosettaScripts*, and all are accessible in PyRosetta. These classes are currently in Rosetta weekly releases as of the December 2019, and can be used for complex scripting purposes. Documentation on each of these can be found on the Rosetta docs page. Classes that are not scriptable have documentation in-code, which can be accessed through PyRosetta's help features, or the PyRosetta API webpage.

Table S7: *RosettaCarbohydrate* and General Extensions (RS indicates accessibility in Rosetta Scripts)

| Component | Description | RS? |
|---|---|---|
| **Modeling** | | |
| *GlycanTreeModeler* | Model glycans through a layer-based, optimized algorithm | Yes |
| *GlycanSampler* | Sample a variety of glycan DOFs through a weighted sampler | Yes |
| *SmallBBSampler* | Sample any backbone torsion in a residue based on a delta for that torsion | No |
| *SugarBBSampler* | Sample sugar torsions using *sugar_bb* QM data as probabilities | No |
| *SimpleGlycosylateMover* | Add a glycan to a pose or to a specific residue on a protein through common names or IUPAC definitions | Yes |
| *GlycanInfoMover* | Get detailed information about the glycans in your pose, especially connectivity information and glycoprotein connections. | Yes |
| **Design** | | |
| *CreateGlycanSequonMover* | Mutates residues to create a potential glycosylation site using known sequence motifs of N- or C-linked glycans. Includes options for Enhanced Sequons for N-linked glycans that have been shown to have higher rates of glycosylation as well as other positions that have been shown to influence the glycosylation chemistry. | Yes |
| *CreateSequenceMotifMover* | Simple mover to Create a sequence motif in a region of protein using the SequenceMotifTaskOperation. Uses pseudo-regular expressions to define the motif. | Yes |
| *SequenceMotifTaskOperation* | A TaskOperation that takes a regex-like pattern and turns it into a set of design residues. The string should identify what to do for each position. | Yes |
| *ResfileCommandOperation* | A TaskOperation for design. Apply a resfile command to a set of residues from a residue selector. | Yes |
| **ResidueSelectors** | | |

| | | |
|---|---|---|
| *GlycanResidueSelector* | A ResidueSelector for carbohydrates and individual carbohydrate trees. Selects all Glycan residues if no option is given or the branch going out from the root residue. Selecting from root residues allows you to choose the whole glycan branch or only tips, etc. | Yes |
| *GlycanLayerSelector* | Selects glycan residues by layer | Yes |
| *GlycanPositionSelector* | Selects glycan residues by position in the tree. Max position is the length of the particular tree | Yes |
| *RandomGlycanFoliageSelector* | Selects a random carbohydrate residue from a subset or selector, then selects the rest of the glycan foliage. Used for sampling. | Yes |
| *DensityFitResidueSelector* | Selects residues based on their correlation to the associated density. Can use a SimpleMetric cache | Yes |
| *ResiduePropertySelector* | A residue selector that selects based on set residue properties. Default is to use AND logic for multiple properties. This can be changed via set_selection_logic. | Yes |
| **SimpleMetrics** | | |
| *RunSimpleMetricsMover* | Runs a set of SimpleMetrics and adds the data to the pose for output into the scorefile. Accepts prefix and suffix options. | Yes |
| *SimpleMetricFilter* | Allows use of SimpleMetrics as filters | Yes |
| *SimpleMetricFeatures* | Use SimpleMetrics in the Features Reporter framework | Yes |
| **Other** | | |
| *GlycanTreeSet* | Holds connectivity information about all glycan trees in a pose. Part of the Conformation object. Created on loading a PDB, or adding glycan residues to a pose. Auto-updates when adding or removing residues from a pose. Holds GlycanTree objects | No |
| *GlycanTree* | Holds connectivity information of a glycan, including it's connection to any glycoprotein. Holds GlycanNodes | No |
| *GlycanNode* | Holds extra information about a glycan residue including it's parent residue, children, and current distance to the root of the tree. | No |
| *ConvertRealToVirtualMover* | Convert a set of residues to virtual - where they are not scored. Used for layer-based sampling. | Yes |
| *ConvertVirtualToRealMover* | Convert a 'virtual' residue back to real residue. Used for layer-based sampling. | Yes |

### *Water-mediated Hydrogen Bonds*

Water-mediated hydrogen bonds were calculated using the *WaterMediatedHbondMetric* with the following script. The options -include_waters and -flip_HNQ were set to true to load in crystallographic waters and allow HNQ flipping during the optH protocol.

```
<ROSETTASCRIPTS>
	<SCOREFXNS>
	</SCOREFXNS>
	<RESIDUE_SELECTORS>
		<Glycan name="tree" branch="%%branch%%" include_root="0" />
		<Glycan name="tree_and_root" branch="%%branch%%" include_root="1"/>
		<Index name="root" resnums="%%branch%%" />
		<GlycanLayerSelector name="first_layer" start="0" end="1"/>
		<And name="layer01" selectors="tree,first_layer" />
		<Neighborhood name="tree_root_neighbors" selector="tree_and_root" include_focus_in_subset="1"/>
		<Not name="not_tree_and_neighbors" selector="tree_root_neighbors" />

		<Neighborhood name="tree_and_neighbors" selector="tree" include_focus_in_subset="1"/>
		<ResidueName name="waters" residue_name3="HOH"/>
	</RESIDUE_SELECTORS>
	<SIMPLE_METRICS>
		<WaterMediatedHbondMetric name="wmhb1" depth="1" residue_selector="tree" residue_selector2="not tree and
not waters"/>
		<WaterMediatedHbondMetric name="wmhb2" depth="2" residue_selector="tree" residue_selector2="not tree and
not waters"/>
		<WaterMediatedHbondMetric name="wmhb3" depth="3" residue_selector="tree" residue_selector2="not tree and
not waters"/>
```

```
                        <ResidueSummaryMetric name="wmhb1_mean" metric="wmhb1" action="mean"
custom_type="wmhb_mean_depth1" />
                        <ResidueSummaryMetric name="wmhb2_mean" metric="wmhb2" action="mean"
custom_type="wmhb_mean_depth2" />
                        <ResidueSummaryMetric name="wmhb3_mean" metric="wmhb3" action="mean"
custom_type="wmhb_mean_depth3" />

                        <ResidueSummaryMetric name="wmhb1_sum" metric="wmhb1" action="sum"
custom_type="wmhb_sum_depth1"/>
                        <ResidueSummaryMetric name="wmhb2_sum" metric="wmhb2" action="sum"
custom_type="wmhb_sum_depth2"/>
                        <ResidueSummaryMetric name="wmhb3_sum" metric="wmhb3" action="sum"
custom_type="wmhb_sum_depth3"/>

                        <SelectedResidueCountMetric name="tree_length" residue_selector="tree" custom_type="tree_length"/>
                        <SelectedResidueCountMetric name="total_waters" residue_selector="waters" custom_type="total_waters"/>
                        <SelectedResidueCountMetric name="total_waters_nbr" residue_selector="waters AND tree_root_neighbors"
custom_type="water_nbrs"/>

            <ProtocolSettingsMetric name="protocol" get_user_options="0" limit_to_options="branch" job_tag="%%exp%%"/>
                        <SelectedResiduesPyMOLMetric name="focus_selection" custom_type="hoh_area"
residue_selector="tree_root_neighbors"/>

            <SelectedResiduesPyMOLMetric   name="pymol_tree"   residue_selector="tree"   custom_type="glycans"/>
            <SelectedResiduesPyMOLMetric   name="pymol_branch" residue_selector="root"   custom_type="branch"/>
            <SelectedResiduesMetric        name="pdb_glycans"  residue_selector="tree"   rosetta_numbering="0" custom_type="glycans"/>
            <SelectedResiduesMetric        name="pdb_branch"   residue_selector="root"   rosetta_numbering="0" custom_type="branch"/>

            </SIMPLE_METRICS>
            <TASKOPERATIONS>
                        <OptH name="opth"/>
                        <InitializeFromCommandline name="init"/>
                        <RestrictToRepacking name="rtrp"/>
                        <OperateOnResidueSubset name="freeze_others" selector="not_tree_and_neighbors">
                                    <PreventRepackingRLT/>
                        </OperateOnResidueSubset>

                        <OperateOnResidueSubset name="only_waters" selector="not waters">
                                    <PreventRepackingRLT/>
                        </OperateOnResidueSubset>

                        <OperateOnResidueSubset name="only_waters_and_tree" selector="tree_and_root OR not waters">
                                    <PreventRepackingRLT/>
                        </OperateOnResidueSubset>
            </TASKOPERATIONS>
            <MOVERS>
                        <ExplicitWaterMover name="solvate" mode="replace" gen_fixed="0" scorefxn="commandline"
task_operations="opth,init,rtrp,freeze_others" />

                        <PackRotamersMover name="pack_nbr_waters_and_tree" scorefxn="commandline"
task_operations="opth,init,rtrp,freeze_others"/>
                        <PackRotamersMover name="pack" scorefxn="commandline" task_operations="opth,init,rtrp,freeze_others"/>
                        <PackRotamersMover name="pack_waters" scorefxn="commandline"
task_operations="opth,init,rtrp,only_waters"/>
                        <RunSimpleMetrics name="selections" metrics="pymol_tree,pymol_branch,pdb_glycans,pdb_branch" />
                        <RunSimpleMetrics name="run_metrics"
metrics="protocol,tree_length,focus_selection,wmhb1_mean,wmhb2_mean,wmhb3_mean,wmhb1_sum,wmhb2_sum,wmhb3_sum"/>
            </MOVERS>
            <PROTOCOLS>
                        <Add mover_name="pack"/>
                        <Add mover_name="run_metrics"/>
                        <Add mover_name="selections"/>
            </PROTOCOLS>
            <OUTPUT />
</ROSETTASCRIPTS>
```

#### *Explicit Solvent modeling*

Rosetta-ECO was used to explicitly model waters around the modeling glycan for each decoy after glycan building. In this way, the explicit waters were to potentially improve decoy discrimination. Glycans were modeled and solvated with the following script. 8 glycans were used as examples for this small benchmark. 4 best and 4 worst-performing glycans. Glycans that had no native interactions with symmetry mates were used as Rosetta-ECO is not compatible with symmetry.

```
<ROSETTASCRIPTS>
	<SCOREFXNS>
	</SCOREFXNS>

	NEEDED FOR CACHING density fit info
	<RESIDUE_SELECTORS>
		<Glycan name="tree" branch="%%branch%%" include_root="0" />
	</RESIDUE_SELECTORS>
	<SIMPLE_METRICS>
		<PerResidueDensityFitMetric name="fit_native" residue_selector="tree" output_as_pdb_nums="1"
sliding_window_size="1" match_res="1"/>
	</SIMPLE_METRICS>

	<RESIDUE_SELECTORS>

		<Index name="root" resnums="%%branch%%" />
		<Glycan name="tree_and_root" branch="%%branch%%" include_root="1"/>
		<Neighborhood name="tree_root_neighbors" selector="tree_and_root" include_focus_in_subset="1"/>
		<GlycanLayerSelector name="first_layer" start="0" end="1"/>
		<And name="layer01" selectors="tree,first_layer" />
		<DensityFitResidueSelector name="fits8" den_fit_metric="fit_native" cutoff=".8" use_cache="1"
fail_on_missing_cache="1" prefix="native_"/>
		<DensityFitResidueSelector name="fits6" den_fit_metric="fit_native" cutoff=".6" use_cache="1"
fail_on_missing_cache="1" prefix="native_"/>
		<DensityFitResidueSelector name="fits4" den_fit_metric="fit_native" cutoff=".4" use_cache="1"
fail_on_missing_cache="1" prefix="native_"/>

		<ResidueName name="waters" residue_name3="HOH"/>

		<Neighborhood name="tree_and_neighbors" selector="tree" include_focus_in_subset="1"/>

	</RESIDUE_SELECTORS>
	<SIMPLE_METRICS>
	<RMSDMetric  name="rmsd"	use_native="1" rmsd_type="rmsd_all_heavy"	residue_selector="tree"/>
	<RMSDMetric  name="rmsd_layer01" use_native="1" rmsd_type="rmsd_all_heavy"	residue_selector="layer01"
custom_type="layer01"/>

	<RMSDMetric  name="rmsd_layer01_super" use_native="1" rmsd_type="rmsd_all_heavy"	residue_selector="layer01"
custom_type="layer01_super" super="1"/>

	<RMSDMetric  name="rmsd_super"	use_native="1" rmsd_type="rmsd_all_heavy" residue_selector="tree"
custom_type="super" super="1"/>
	<RMSDMetric  name="rmsd_aligned_layer01" use_native="1" rmsd_type="rmsd_all_heavy" residue_selector="layer01"
custom_type="layer12_aln"  super="1"/>

	<PerResidueRMSDMetric name="rmsd_rsd"	use_native="1" rmsd_type="rmsd_all_heavy" residue_selector="tree"
output_as_pdb_nums="1"/>
	<PerResidueRMSDMetric name="rmsd_aligned_rsd"	use_native="1" rmsd_type="rmsd_all_heavy" residue_selector="tree"
output_as_pdb_nums="1" super="1" custom_type="aln"/>

	<TimingProfileMetric name="timing" />
```

```
		<SelectedResidueCountMetric name="n_tree" custom_type="tree_size" residue_selector="tree"/>
		<SelectedResidueCountMetric name="n_fits8" custom_type="fit8" residue_selector="fits8"/>
		<SelectedResidueCountMetric name="n_fits6" custom_type="fit6" residue_selector="fits6"/>
		<SelectedResidueCountMetric name="n_fits4" custom_type="fit4" residue_selector="fits4"/>
		<SelectedResidueCountMetric name="n_layer01" custom_type="layer01" residue_selector="layer01"/>

		<RMSDMetric name="rmsd_fits8" use_native="1" custom_type="fit8" rmsd_type="rmsd_all_heavy" residue_selector="fits8"/>
		<RMSDMetric name="rmsd_fits6" use_native="1" custom_type="fit6" rmsd_type="rmsd_all_heavy" residue_selector="fits6"/>
		<RMSDMetric name="rmsd_fits4" use_native="1" custom_type="fit4" rmsd_type="rmsd_all_heavy" residue_selector="fits4"/>


		<RMSDMetric name="rmsd_fits8_super" use_native="1" custom_type="fit8_super" rmsd_type="rmsd_all_heavy"
residue_selector="fits8" super="1" residue_selector_super="tree"/>
		<RMSDMetric name="rmsd_fits6_super" use_native="1" custom_type="fit6_super" rmsd_type="rmsd_all_heavy"
residue_selector="fits6" super="1" residue_selector_super="tree"/>
		<RMSDMetric name="rmsd_fits4_super" use_native="1" custom_type="fit4_super" rmsd_type="rmsd_all_heavy"
residue_selector="fits4" super="1" residue_selector_super="tree"/>


		<PerResidueGlycanLayerMetric   name="layers"        residue_selector="tree" output_as_pdb_nums="1"/>
		<SelectedResiduesPyMOLMetric   name="pymol_tree"   residue_selector="tree"   custom_type="glycans"/>
		<SelectedResiduesPyMOLMetric   name="pymol_branch" residue_selector="root"   custom_type="branch"/>
		<SelectedResiduesMetric        name="pdb_glycans"  residue_selector="tree"   rosetta_numbering="0" custom_type="glycans"/>
		<SelectedResiduesMetric        name="pdb_branch"   residue_selector="root"   rosetta_numbering="0" custom_type="branch"/>

		<TotalEnergyMetric name="total" scorefxn="commandline"/>
		<TotalEnergyMetric name="total_glycans" scorefxn="commandline" residue_selector="tree" custom_type="glycan"/>
		<ProtocolSettingsMetric name="protocol" get_user_options="0"
limit_to_options="rounds,glycan_sampler_rounds,window_size,layer_size,quench_mode,conformer_probs,gaussian_sampling"
job_tag="%%exp%%"/>

		<SelectedResiduesPyMOLMetric name="focus_selection" custom_type="hoh_area" residue_selector="tree_root_neighbors"/>
			<WaterMediatedHbondMetric name="wmhb1" depth="1" residue_selector="tree" residue_selector2="not tree and
not waters"/>
			<WaterMediatedHbondMetric name="wmhb2" depth="2" residue_selector="tree" residue_selector2="not tree and
not waters"/>
			<WaterMediatedHbondMetric name="wmhb3" depth="3" residue_selector="tree" residue_selector2="not tree and
not waters"/>

			<ResidueSummaryMetric name="wmhb1_mean" metric="wmhb1" action="mean"
custom_type="wmhb_mean_depth1" />
			<ResidueSummaryMetric name="wmhb2_mean" metric="wmhb2" action="mean"
custom_type="wmhb_mean_depth2" />
			<ResidueSummaryMetric name="wmhb3_mean" metric="wmhb3" action="mean"
custom_type="wmhb_mean_depth3" />

			<ResidueSummaryMetric name="wmhb1_sum" metric="wmhb1" action="sum"
custom_type="wmhb_sum_depth1"/>
			<ResidueSummaryMetric name="wmhb2_sum" metric="wmhb2" action="sum"
custom_type="wmhb_sum_depth2"/>
			<ResidueSummaryMetric name="wmhb3_sum" metric="wmhb3" action="sum"
custom_type="wmhb_sum_depth3"/>

			<SelectedResidueCountMetric name="tree_length" residue_selector="tree" custom_type="tree_length"/>
			<SelectedResidueCountMetric name="total_waters" residue_selector="waters" custom_type="total_waters"/>
			<SelectedResidueCountMetric name="total_waters_nbr" residue_selector="waters AND tree_root_neighbors"
custom_type="water_nbrs"/>


		</SIMPLE_METRICS>
		<TASKOPERATIONS>
			<OptH name="opth"/>
			<InitializeFromCommandline name="init"/>
			<RestrictToRepacking name="rtrp"/>
			<OperateOnResidueSubset name="freeze_others" selector="not tree_and_neighbors">
				<PreventRepackingRLT/>
			</OperateOnResidueSubset>

			<OperateOnResidueSubset name="only_waters" selector="not waters">
```

```
                    <PreventRepackingRLT/>
                </OperateOnResidueSubset>

                <OperateOnResidueSubset name="only_waters_and_tree" selector="tree_and_root OR not waters">
                    <PreventRepackingRLT/>
                </OperateOnResidueSubset>
        </TASKOPERATIONS>
        <MOVERS>
        <SetupForSymmetry name="setup_symm" definition="%%symmdef%%"/>
        <LoadDensityMap name="loaddens" mapfile="%%map%%"/>
        <SetupForDensityScoring name="setupdens"/>
                <GlycanTreeModeler name="tree_relax" quench_mode="%%quench_mode%%" layer_size="%%layer_size%%"
window_size="%%window_size%%" residue_selector="tree" cartmin="%%cartmin%%" scorefxn="commandline"
glycan_sampler_rounds="%%glycan_sampler_rounds%%" rounds="%%rounds%%" use_conformer_probs="%%conformer_probs%%"
use_gaussian_sampling="%%gaussian_sampling%%" shear="%%shear%%" hybrid_protocol="%%hybrid_protocol%%"
match_window_one="%%match%%" root_populations_only="%%root_probs%%"/>

                <RunSimpleMetrics name="native_metrics" metrics="fit_native,total" prefix="native_"/>
                <RunSimpleMetrics name="selections" metrics="layers,pymol_tree,pymol_branch,pdb_glycans,pdb_branch" />
                <RunSimpleMetrics name="counts"     metrics="n_tree,n_layer01,n_fits6,n_fits8"/>
                <RunSimpleMetrics name="timings"    metrics="timing" />
                <RunSimpleMetrics name="rmsd"       metrics="rmsd,rmsd_layer01,rmsd_rsd,rmsd_super"/>
                <RunSimpleMetrics name="rmsd_fits"  metrics="rmsd_fits8,rmsd_fits6,rmsd_fits6_super"/>
                <RunSimpleMetrics name="energies" metrics="total,total_glycans"/>
                <RunSimpleMetrics name="settings" metrics="protocol" />
                <RunSimpleMetrics name="water_metrics"
metrics="tree_length,focus_selection,wmhb1_mean,wmhb2_mean,wmhb3_mean,wmhb1_sum,wmhb2_sum,wmhb3_sum"/>

                <ExplicitWaterMover name="solvate" mode="replace" gen_fixed="0" scorefxn="commandline"
task_operations="opth,init,rtrp,freeze_others" />

                <PackRotamersMover name="pack" scorefxn="commandline" task_operations="opth,init,rtrp,freeze_others"/>
        </MOVERS>
        <APPLY_TO_POSE>
        </APPLY_TO_POSE>
        <PROTOCOLS>
                <Add mover_name="loaddens"/>
                <Add mover_name="setupdens"/>
                <Add mover_name="selections"/>
                <Add mover_name="native_metrics" />
                <Add mover_name="counts"/>
                <Add mover_name="tree_relax" />
                <Add mover_name="solvate"/>
                <Add mover_name="pack"/>
                <Add mover_name="water_metrics"/>
                <Add mover_name="energies" />
                <Add mover_name="rmsd"/>
                <Add mover_name="rmsd_fits" />
                <Add mover_name="timings"/>
                <Add mover_name="settings"/>
        </PROTOCOLS>
        <OUTPUT />
</ROSETTASCRIPTS>
```

All water modeling used the following additional flags:
-corrections::water::wat_rot_sampling 20

#### *Benchmarking Options and Scripts for Reproducibility*

For each experimental group, a Job Definition (JD) file was used to define the experimental conditions. For each experiment within that group, a RosettaScript XML was used to define the protocol and metrics. The script, *create_substituted_JD.py* was used to create a final, substituted JD file by substituting specific variables for each PDB and glycan tree that became each job (i.e. 3 experiments across 25 glycan trees becomes 75 independent jobs). This script is included in this work for reproducibility purposes.

The Rosetta flags (options) file that was used for all benchmarking is transcribed below. Additional flags are then given for each experiment.

```
# Input
-ignore_unrecognized_res
-ignore_zero_occupancy false
-load_PDB_components false

# Output
-pdb_comments
-out:pdb_gz
-scorefile_format json

# Rotamers/packing
-ex1
-ex2
-use_input_sc

# Minimization
-ideal_sugars

# Glycan
-include_sugars
-auto_detect_glycan_connections
-alternate_3_letter_codes pdb_sugar
-maintain_links
-write_pdb_link_records
-write_glycan_pdb_codes

# JD3
-mpi_fraction_outputters .05
-skip_connect_info
-cryst::crystal_refine
```

The primary RosettaScript XML is shown below. Most of the content of the script is setting up and running SimpleMetrics at key points in the protocol. Any additional specific scripts for experiments are given in their respective section.

***glycan_tree_modeler.xml***

```
<ROSETTASCRIPTS>
		<SCOREFXNS>
		</SCOREFXNS>
		NEEDED FOR CACHING density fit info
		<RESIDUE_SELECTORS>
			<Glycan name="tree" branch="%%branch%%" include_root="0" />
		</RESIDUE_SELECTORS>
		<SIMPLE_METRICS>
			<PerResidueDensityFitMetric name="fit_native" residue_selector="tree" output_as_pdb_nums="1" sliding_window_size="1" match_res="1"/>
		</SIMPLE_METRICS>
		<RESIDUE_SELECTORS>
			<Index name="root" resnums="%%branch%%" />
			<GlycanLayerSelector name="first_layer" start="0" end="1"/>
			<And name="layer01" selectors="tree,first_layer" />
			<DensityFitResidueSelector name="fits8" den_fit_metric="fit_native" cutoff=".8" use_cache="1" fail_on_missing_cache="1" prefix="native_"/>
			<DensityFitResidueSelector name="fits6" den_fit_metric="fit_native" cutoff=".6" use_cache="1" fail_on_missing_cache="1" prefix="native_"/>
		</RESIDUE_SELECTORS>
		<SIMPLE_METRICS>
			<RMSDMetric name="rmsd"     use_native="1" rmsd_type="rmsd_all_heavy"     residue_selector="tree"/>
			<RMSDMetric name="rmsd_layer01" use_native="1" rmsd_type="rmsd_all_heavy"     residue_selector="layer01" custom_type="layer01"/>
```

```
			<RMSDMetric name="rmsd_super"	use_native="1" rmsd_type="rmsd_all_heavy" residue_selector="tree"  custom_type="super" super="1"/>
			<PerResidueRMSDMetric name="rmsd_rsd"	use_native="1" rmsd_type="rmsd_all_heavy" residue_selector="tree" output_as_pdb_nums="1"/>
			<PerResidueRMSDMetric name="rmsd_aligned_rsd"	use_native="1" rmsd_type="rmsd_all_heavy" residue_selector="tree" output_as_pdb_nums="1" super="1"
custom_type="aln"/>
			<TimingProfileMetric name="timing" />
			<RMSDMetric name="rmsd_fits8" use_native="1" custom_type="fit8" rmsd_type="rmsd_all_heavy" residue_selector="fits8"/>
			<RMSDMetric name="rmsd_fits6" use_native="1" custom_type="fit6" rmsd_type="rmsd_all_heavy" residue_selector="fits6"/>
			<RMSDMetric name="rmsd_fits6_super" use_native="1" custom_type="fit6_super" rmsd_type="rmsd_all_heavy" residue_selector="fits6" super="1"
residue_selector_super="tree"/>

			<PerResidueGlycanLayerMetric  name="layers"	residue_selector="tree" output_as_pdb_nums="1"/>
			<SelectedResiduesPyMOLMetric  name="pymol_tree"  residue_selector="tree"  custom_type="glycans"/>
			<SelectedResiduesPyMOLMetric  name="pymol_branch" residue_selector="root"  custom_type="branch"/>
			<SelectedResiduesMetric	name="pdb_glycans" residue_selector="tree"  rosetta_numbering="0" custom_type="glycans"/>
			<SelectedResiduesMetric	name="pdb_branch"  residue_selector="root"  rosetta_numbering="0" custom_type="branch"/>

			<TotalEnergyMetric name="total" scorefxn="commandline"/>
			<TotalEnergyMetric name="total_glycans" scorefxn="commandline" residue_selector="tree" custom_type="glycan"/>
			<ProtocolSettingsMetric name="protocol" get_user_options="0"
limit_to_options="rounds,glycan_sampler_rounds,window_size,layer_size,quench_mode,conformer_probs,gaussian_sampling" job_tag="%%exp%%"/>
		</SIMPLE_METRICS>
		<MOVERS>
			<SetupForSymmetry name="setup_symm" definition="%%symmdef%%"/>
			<LoadDensityMap name="loaddens" mapfile="%%map%%"/>
			<SetupForDensityScoring name="setupdens"/>
			<GlycanTreeModeler name="tree_relax" quench_mode="%%quench_mode%%" layer_size="%%layer_size%%" window_size="%%window_size%%"
residue_selector="tree" cartmin="%%cartmin%%" scorefxn="commandline" glycan_sampler_rounds="%%glycan_sampler_rounds%%" rounds="%%rounds%%"
use_conformer_probs="%%conformer_probs%%" use_gaussian_sampling="%%gaussian_sampling%%" shear="%%shear%%" hybrid_protocol="%%hybrid_protocol%%"
match_window_one="%%match%%" root_populations_only="%%root_probs%%"/>
			<RunSimpleMetrics name="native_metrics" metrics="fit_native,total" prefix="native_"/>
			<RunSimpleMetrics name="selections" metrics="layers,pymol_tree,pymol_branch,pdb_glycans,pdb_branch" />
			<RunSimpleMetrics name="timings"  metrics="timing" />
			<RunSimpleMetrics name="rmsd"	metrics="rmsd,rmsd_layer01,rmsd_rsd,rmsd_super"/>
			<RunSimpleMetrics name="rmsd_fits" metrics="rmsd_fits8,rmsd_fits6,rmsd_fits6_super"/>
			<RunSimpleMetrics name="energies" metrics="total,total_glycans"/>
			<RunSimpleMetrics name="settings" metrics="protocol" />
		</MOVERS>
		<APPLY_TO_POSE>
		</APPLY_TO_POSE>
		<PROTOCOLS>
			<Add mover_name="setup_symm" />
			<Add mover_name="loaddens"/>
			<Add mover_name="setupdens"/>
			<Add mover_name="selections"/>
			<Add mover_name="native_metrics" />
			<Add mover_name="counts"/>
			<Add mover_name="tree_relax" />
			<Add mover_name="energies" />
			<Add mover_name="rmsd"/>
			<Add mover_name="rmsd_fits" />
			<Add mover_name="timings"/>
			<Add mover_name="settings"/>
		</PROTOCOLS>
		<OUTPUT />
</ROSETTASCRIPTS>
```

### An example Job Definition file that was expanded for each pdb and branch using the previously mentioned script is below. A job for each experiment run in parallel for different script_vars was used during benchmarking.:

```
<JobDefinitionFile>
  <Job>
    <Input>
      <PDB filename="%%fname%%"/>
    </Input>
    <Output>
      <PDB filename_pattern="final3_%%branch%%/final3_beta_%%branch%%_$"/>
    </Output>
    <Options>
      <parser__protocol value="xmls/glycan_tree_relax.xml"/>
      <parser__script_vars value="branch=%%branch%% cartmin=0 layer_size=1 window_size=0 glycan_sampler_rounds=100
quench_mode=0 map=%%map%% symmdef=%%symmdef%% shear=1 rounds=1 conformer_probs=0 gaussian_sampling=1
hybrid_protocol=1 exp=final3 root_probs=0 match=1"/>
      <score__set_weights value="sugar_bb .5"/>
      <in__file__native value="%%fname%%" />
    </Options>
  </Job>
</JobDefinitionFile>
```
