## supplemental_data_and_movie for "Growing Glycans in Rosetta: Accurate *de novo* glycan modeling, density fitting, and rational sequon design": create_substituted_JD.py

```
#!/usr/bin/env python
from __future__ import print_function
from collections import defaultdict
from argparse import ArgumentParser
import re,os,sys,glob


def replace_line(line, pdb, map_file, sym_file, main_outdir, branch="", skip_dir_creation=False):
    """
    Replace the line with passed variables. 
    :param line: 
    :param pdb: 
    :param map: 
    :param branch: 
    :return: 
    """

    new_line = line.replace("%%fname%%", pdb)
    if re.search("%%map%%", line):
        new_line = new_line.replace("%%map%%", map_file)
    if re.search("%%symmdef%%", line):
        new_line = new_line.replace("%%symmdef%%", sym_file)
    if branch:
        new_line = new_line.replace("%%branch%%", branch)


    if re.search("<PDB filename_pattern=", new_line):
        outpath = new_line.replace("<PDB filename_pattern=", "")
        outpath = outpath.replace("/>", " ")
        outpath = outpath.strip()
        outpath = outpath.strip('"')
        #print(outpath)

        d = os.path.join( main_outdir, os.path.dirname(outpath))
        if not os.path.exists(d) and not skip_dir_creation:
            os.mkdir(d)
            os.system('touch '+d+'/.gitignore')

    return new_line


def split_input_jd_to_jobs(lines):
    """
    Return a dict of the filename_pattern and the Job (including Job tags)
    
    :param lines: 
    :return: defaultdict
    """

    jobs = defaultdict(list)
    final_jobs = defaultdict()
    job_num = 0
    for line in lines:
        if not re.search('<', line):
            continue
        elif re.search("JobDefinition", line):
            continue
        elif re.search("<Job>", line):
            job_num+=1
            jobs[job_num].append(line)
        else:
            jobs[job_num].append(line)

    for job_num in jobs:
        #print("\n"+str(job_num))
       #print(repr(jobs[job_num]))

        pdb_path = ""
        for line in jobs[job_num]:
            if (re.search("<PDB filename_pattern=", line)):
                lineSP = line.split('=')
                pdb_path = lineSP[1].strip("/>\n").strip('"').strip('$')
                final_jobs[pdb_path] = jobs[job_num]
                break

    #for pdb in final_jobs:
        #print("\n")
        #print(pdb)
        #print(repr(final_jobs[pdb]))

    return final_jobs

def is_job_done(job_name, pdb, outdir, nstruct=2500, ext=".pdb.gz"):
    """
    Checks to see if all nstruct are present. 
    Returns true if job is done
    
    :param job_name: 
    :param nstruct: 
    :return: bool
    """
    final_name = outdir+"/"+job_name+pdb+"_refined"
    #print("GLOB: "+final_name+"*"+ext)
    files = glob.glob(final_name+"*"+ext)
    #print("NFILES: "+(str(len(files))))
    #print("nstruct:", nstruct)
    if len(files) == int(nstruct):
        print(final_name, "Is Complete")
        return True
    else:
        print(final_name, "Is missing", str(int(nstruct) - len(files)))
        return False


if __name__ == "__main__":
    parser = ArgumentParser("Input a base xml and output a substituted XML using a TSV of pdb/braches/skipped_residues "
                            "for glycan benchmarking.  Also creates PDB List for benchmarks.")

    parser.add_argument("-j", "--jd", help = "Input Job Definition XML", required=True)
    parser.add_argument("-i", "--pdbs", help = "Input PDB TSV", default = "pdb_roots_density_sym.txt")
    parser.add_argument("-o", "--prefix", help = "Any additional prefix for output XML (will use xml name)", default="")
    parser.add_argument("-p", "--pdbdir", help = "Input directory for PDBs", default = "pdbs")
    parser.add_argument("-m", "--mapdir", help = "Map directory for PDBs", default="maps")
    parser.add_argument("-e", "--ext", help = "PDB Extension. ", default = "_refined.pdb.gz")
    parser.add_argument("-s", "--symdir", help = "Dir with .symm files", default="pdbs/symmetrized_structures")
    parser.add_argument("-d", "--decoy_dir", help = "Decoy dir for jobs.  Will create sub-directories with gitignore for cluster run here", default="decoys")
    parser.add_argument("-c", "--checkpoint",
                        default = False,
                        action = "store_true",
                        help ="Checkpoint the output since we have problems with JD3.  "
                                "Write out a JD with only those jobs missing full PDBs.")

    parser.add_argument("-n", "--nstruct",
                        default=2500,
                        help = "Nstruct we are using.  only used for checkpointing fix.")

    parser.add_argument("--skip_dir_creation", 
                        action = "store_true",
                        default=False, 
                        help="Skip creation of subdirectories")

    options = parser.parse_args()

    if not os.path.exists(options.decoy_dir):
        os.mkdir(options.decoy_dir)

    xml_name = ".".join(os.path.basename(options.jd).split('.')[:-1])
    if options.checkpoint:
        outname = options.prefix+xml_name+"_checkpoint_substituted.xml"
    else:
        outname = options.prefix + xml_name + "_substituted.xml"

    pdblist = open( options.prefix+xml_name+"_PDBLIST.txt", 'w')

    pdb_branches = defaultdict(list)
    for line in open(options.pdbs):
        line = line.strip()
        if not line or line.startswith("#"): continue
        lineSP = line.split()
        pdb = lineSP[0].strip()

        pdb_path = options.pdbdir + "/"+ pdb + options.ext
        pdblist.write(pdb_path+'\n')
        for branch in lineSP[1].split(','):
            branch = branch.strip()

            pdb_branches[pdb_path].append(branch)

    pdblist.close()

    print("wrote: "+options.prefix+xml_name+"_PDBLIST.txt")
    new_lines = []
    unparsed = open(options.jd, 'r').readlines()

    jobs = split_input_jd_to_jobs(unparsed)


    new_lines.append("<JobDefinitionFile>\n")

    branches = False
    for line in unparsed:
        if re.search("%%branch%%", line ):
            branches = True
            break

    #Parse experiments:
    n_jobs = 0
    for pdb in pdb_branches:
        #n_jobs+=1
        if not os.path.exists(pdb):
            sys.exit(pdb + " does not exist!")

        pdbid = os.path.basename(pdb).split(".")[0].split('_')[0]
        print("Working On:", pdbid)
        map_file = options.mapdir+"/"+ pdbid+"_2mFo-DFc_map.ccp4"
        #print(map_file)
        sym_file = options.symdir+"/"+ pdbid+ "_crys.symm"
        if not os.path.exists(map_file):
            sys.exit(map_file+" does not exist!")
        if not os.path.exists(sym_file):
            sys.exit(sym_file+" does not exist!")
        if branches:
            for branch in pdb_branches[pdb]:
                n_jobs+=1
                if options.checkpoint:
                    for job in jobs:
                        #print("JOB: "+job)
                        job_name = replace_line(job, pdb, map_file, sym_file, options.decoy_dir, branch, options.skip_dir_creation)
                        job_complete = is_job_done(job_name, pdbid, options.decoy_dir, options.nstruct)
                        if job_complete:
                            continue
                        else:
                            for line in jobs[job]:
                                new_line = replace_line(line, pdb, map_file, sym_file, options.decoy_dir, branch, options.skip_dir_creation)
                                new_lines.append(new_line)
                else:
                    for line in unparsed:
                        if re.search("JobDefinitionFile", line): continue
                        new_line = replace_line(line, pdb, map_file, sym_file, options.decoy_dir, branch, options.skip_dir_creation)
                        new_lines.append(new_line)
        else:

            n_jobs+=1
            if options.checkpoint:
                for job in jobs:
                    job_name = replace_line(job, pdb, map_file, sym_file, options.decoy_dir, "", options.skip_dir_creation)

                    job_complete = is_job_done(job_name, pdbid, options.decoy_dir, options.nstruct)
                    if job_complete:
                        continue
                    else:
                        for line in jobs[job]:
                            new_line = replace_line(line, pdb, map_file, sym_file, options.decoy_dir, "", options.skip_dir_creation)
                            new_lines.append(new_line)
            else:
                for line in unparsed:
                    if re.search("JobDefinitionFile", line): continue
                    new_line = replace_line(line, pdb, map_file, sym_file, options.decoy_dir, "", options.skip_dir_creation)
                    new_lines.append(new_line)

    print("N Jobs: ", n_jobs)
    new_lines.append("</JobDefinitionFile>")
    OUT = open("job_definitions/"+outname, 'w')
    for new_line in new_lines:
        OUT.write(new_line)
    OUT.close()
    pdblist.close()

    print("Done.\nSubstituted file writted to: "+"job_definitions/"+outname)
```
