## supplemental_data_and_movie for "Growing Glycans in Rosetta: Accurate *de novo* glycan modeling, density fitting, and rational sequon design": create_symmetry_mates.py

```
#!/usr/bin/env python

from __future__ import print_function
import sys, os, re
from argparse import ArgumentParser

base_cmd = "cd {dir} && make_symmdef_file.pl -r 12 -m CRYST -p  {pdb}.pdb > {pdb}_crys.symm && cd -"

def run_symmdef_creator(pdb, dir):
    cmd = base_cmd.format(pdb=pdb, dir=dir)

    print("working on " + pdb)
    print(cmd)

    os.system(cmd)
    print(pdb + " complete")

if __name__ == "__main__":

    parser = ArgumentParser("This script uses the pdb_roots_density file to create symm defs.  We then manually check them")

    parser.add_argument("-i", "--pdb_roots_density", default = "pdb_roots_density.txt")
    parser.add_argument("-d", "--dir", default = "pdbs/prelim_benchmark_structures")
    parser.add_argument("-s", "--single")

    options = parser.parse_args()


    if options.single:
        pdb = options.single.strip(".pdb")
        run_symmdef_creator(pdb, options.dir)
    else:

        INFILE = open(options.pdb_roots_density, 'r')

        for line in INFILE:
            line = line.strip()
            if not line: continue;
            if line.startswith('#'): continue;

            lineSP = line.split()
            pdb = lineSP[0]

            run_symmdef_creator(pdb, options.dir)

        print("complete")
```
