## supplemental_data_and_movie for "Growing Glycans in Rosetta: Accurate *de novo* glycan modeling, density fitting, and rational sequon design": glycan_conformer_table.txt

#libraries compiled by Maxim Shapovalov, Jared Adolf-Bryfogle, Jason Labonte, and Roland Dunbrack
#Raw torsion data provided by Thomas Lutteke (Glycosciences.de) 
#Fox Chase Cancer Center, Philadelphia, PA, USA, 2017 (C) and The Scripps Research Institute (C) and Jons Hopkins University (C) 
#Compiled on 6/7/2017 

#METHODS:
# Raw data was obtained from Thomas Lutteke from the PDB as XML files. 
# These were translated to pandas DataFrames and CSV files. 

# The raw data was then filtered for:
# 1) Crystal structures - NO NMR or Unknown methods
# 2) <= 2.0 A resolution. 
# 3) Full torsions for each linkage - Phi/Psi Phi/Psi/Omega, etc. No missing torsions.
# 4) At least 10 datapoints for a specific linkage. 
# 5) Torsion.quality.  These come from the original Glycosciences.de paper:

#      THOMAS LUTTEKE & CLAUS-W VON DER LIETH, The protein data bank (PDB) as a versatile resource for glycobiology and glycomics,
#      Biocatalysis and Biotransformation, Jan /Feb/March /April 2006; 24(1/2): 147 /155
# Filter these values: [2, 3, 4, 5, 6, 10, 16, 32, 1024, 4096, 65536, 524288, 1048576]

#        1  = Positive control
#        2  = Wrong assignment of anomeric
#        3  = PDB residue name and detected monosaccharide are inconsistent
#        4 = Residue name given in PDB file is unknown in the list of residues
#        5 = Stereochemistry of the carbohydrate could not be assigned
#        6 = No glycosidic O -(S -, N -) atom at C1 could be detected
#        7 = Bond between the anomeric C-atom and adjacent O-atom (not part of a ring) was added based on distance criteria
#        8 = Monosaccharide is connected to a chain, which is neither contained in the list of residues norin the list of substituents
#        9 = Dihedral angles derived to calculate the stereochemistry show unusual values
#        10 = Residue at the reducing end of a chain is neither a carbohydrate nor one of the assigned substituents
#        11 = Chain contains a non-oxygen glycosidic atom
#        12 = Inconsistencies between LINUCS notation of composed residues and the PDB residue list
#        13 = Number of monosaccharides is inconsistent with the number of residues given in the PDB-file
#        14 = A non-carbohydrate ring has been detected in a carbohydrate chain
#        15 = Carbohydrate chain is connected to a nucleotide
#        16 = Glycosidic linkage reported in the PDB is not consistent with the detected one
#        17 = A carbohydrate residuum reported in the PDB-file was not detected
#	 32 = residue has no oxygen or respective atom attached to anomeric carbon
#	 64 = residue was 1-deoxy, linkage to o1 or respective atom was added 
#	 128 = residue contains non-standard "sidechain"
#	 256 = residue contains unusual angle in stereochemistry checks 
#	 512 = chain contains non-standard residue at reducing end 
#	 1024 = residue is linked to another residue (except ASN) by an non-oxygen glycosidic atom 
#	 2048 = linucs code comparison error (oligosaccharides / combined residues) 
#	 4096 = number of rings in oligosaccharide doesn't match expected value 
#	 8192 = chain contains "inserted" non-carbohydrate ring 
#	 16384 = chain contains nucleotide residue 
#	 32768 = linkage error (oligosaccharides / combined residues) 
#	 65536 = residue could not be assigned 
#	 131072 = residue consists of more backbone carbons than umfconverter can handle 
#	 262144 = residue has more than one oxygen (or equiv. atoms) attached to the anomeric center, not counting the ring oxygen
#	 524288 = N-glycan chain does not match known biological pathway
#	 1048576 = a monosaccharide is linked to its parent one via a carbon other than the anomeric carbon

#  Conformers were then generated as follows for each linkage type and dihedral angle:
#  A) Bin Identification
#   1) Density was generated using Adaptive Kernal Density estimates using a Von Misses Kernal as was done for the 2010 Rotamer Library. (Matlab - Maxim Shapovolov)
#     1a).  We use a lowPopN strategy to increase kernal width for grid points that have less than 5 members.  This effectively smooths data where it needs to be smoothed
#           and keeps us from overfitting the data. 
#
#                 So what does it do? The code measures that every single density estimate in any grid location is supported by at least N data points within their 
#                 adapted bandwidth. 

#                 If some grid location relies less on N data points, the code forces to expand bandwidth (reduces kappa) across all points. 
#                 Technically we could say LowPopulN does kappa optimization with a constraint that every grid data point is supported by at least N data points. 
#                 Depending on data basically it forbids optimal kappa be above some level.

#                 It is better since you can interpret it statistically even calculate error interval for density estimate. 


#   2) Cubic splines were generated in Matlab on the density using the Spline Package. 
#   3) Inflection points were found by solving the derivitive at zero of the spline. 
#   4) For each inflection point, the second derivitive was used to find the direction of each point.
#   5) A list of maxima was found for each dihedral angle.  These maxima correspond to one-dimensional 'bins/conformer'
#   6) Minima on the left and right of each maxima was found as the boundaries of each torsional bin. 
#   7) For each bin, we integrate over the minima to approximate the probability of the bin. 

#  B) Conformer Identification
#   1) For each linkage, we assign all torsions to a particular bin.  This is the conformer. 
#   2) Fore each linkage type, we calculate means and SDs of each dihedral in the bin. 
#   3) We calculate the probability of the Conformer by taking the N of each conformer over the total number of points for that linkage type. 


# The phi, psi, and omega values listed here are for the non-reducing-end
# residue (the leftmost residue in the table below), in all cases.

# Note that for glycosidic linkages, phi, psi, and omega go upstream (toward
# the LOWER_TERMINUS, or reducing end) and not the other way around.  That is,
# the non-reducing-end residue is numbered after the reducing-end residue in
# the sequence.

# The main-chain designation of the non-reducing-end residue has no effect on
# the torsion values.  Likewise, the anomeric designation of the reducing-end
# residue has no effect on the values.

# For peptide-glycan N-linkages, psi is equivalent to ASN's chi3 (the peptide
# bond, omega1 is equivalent to its chi2 + 180 degrees, and omega2 to chi1.

# IF ADDING NEW ROWS, please sort them such that higher population conformers
# come first.

# reducing-end_res is the lower-numbered residue, while non-reducing-end_res is
# the higher-numbered residued, which is attached to the lower-numbered
# residue.  This ordering from right to left matches IUPAC ordering of chains.

#Conformers with one data member are not used. 


#non-reducing-end	reducing-end	conformer	p	points	phi-mean	phi-sd	psi-mean	psi-sd	omega-mean	omega-sd	omega2-mean	omega2-sd

#beta-D-GlcpNAc ->4)-D-GlcpNAc	TotalConformers=2 PossibleConformers=2 PossibleCombos=2.1
beta-D-GlcpNAc ->4)-D-GlcpNAc	2.1	0.9876	2315	280.880	14.334	235.258	20.313
beta-D-GlcpNAc ->4)-D-GlcpNAc	1.1	0.0124	29	67.481	23.527	245.531	25.986

#beta-D-GlcpNAc ASN	TotalConformers=10 PossibleConformers=12 PossibleCombos=2.2.1.3
beta-D-GlcpNAc ASN	2.1.1.2	0.4755	2116	261.204	18.738	176.582	12.940	173.004	42.546	190.375	12.882
beta-D-GlcpNAc ASN	2.1.1.3	0.3218	1432	266.260	19.488	175.113	11.793	164.505	40.855	289.290	9.885
beta-D-GlcpNAc ASN	2.1.1.1	0.1613	718	260.329	21.021	178.349	15.474	193.810	39.885	63.684	8.287
beta-D-GlcpNAc ASN	1.1.1.1	0.0169	75	63.320	13.298	183.561	16.069	189.020	22.727	66.313	9.523
beta-D-GlcpNAc ASN	1.1.1.3	0.0092	41	61.052	28.933	187.818	27.368	150.766	60.632	293.143	16.627
beta-D-GlcpNAc ASN	1.1.1.2	0.0063	28	66.268	38.218	198.794	46.773	164.958	61.471	195.240	21.508
beta-D-GlcpNAc ASN	2.2.1.2	0.0058	26	276.963	7.572	356.389	6.746	162.310	7.754	193.407	3.919
beta-D-GlcpNAc ASN	2.2.1.1	0.0020	9	250.838	41.359	10.789	21.143	152.733	27.057	63.676	18.049
beta-D-GlcpNAc ASN	1.2.1.2	0.0009	4	62.445	16.943	0.374	2.137	178.520	9.758	201.426	4.712
#beta-D-GlcpNAc ASN	2.2.1.3	0.0002	1	339.000	-0.000	358.700	-0.000	147.600	-0.000	299.300	-0.000

#alpha-D-Glcp ->4)-D-Glcp	TotalConformers=2 PossibleConformers=2 PossibleCombos=1.2
alpha-D-Glcp ->4)-D-Glcp	1.2	0.9805	602	99.937	18.154	229.310	16.992
alpha-D-Glcp ->4)-D-Glcp	1.1	0.0195	12	93.431	30.497	72.140	16.946

#alpha-D-Manp SER	TotalConformers=1 PossibleConformers=1 PossibleCombos=1.1
alpha-D-Manp SER	1.1	1.0000	197	68.584	15.182	178.133	33.276

#beta-D-Manp ->4)-D-GlcpNAc	TotalConformers=1 PossibleConformers=1 PossibleCombos=1.1
beta-D-Manp ->4)-D-GlcpNAc	1.1	1.0000	794	272.910	25.183	228.055	23.562

#alpha-L-Fucp ->3)-D-GlcpNAc	TotalConformers=2 PossibleConformers=2 PossibleCombos=1.2
alpha-L-Fucp ->3)-D-GlcpNAc	1.1	0.9901	200	286.763	15.812	139.149	13.564
alpha-L-Fucp ->3)-D-GlcpNAc	1.2	0.0099	2	265.750	2.150	289.300	0.900

#beta-D-Xylp ->2)-D-Manp	TotalConformers=2 PossibleConformers=2 PossibleCombos=2.1
beta-D-Xylp ->2)-D-Manp	2.1	0.9787	46	277.074	10.063	131.723	7.102
#beta-D-Xylp ->2)-D-Manp	1.1	0.0213	1	66.000	-0.000	157.000	-0.000

#alpha-D-Manp ->3)-D-Manp	TotalConformers=1 PossibleConformers=1 PossibleCombos=1.1
alpha-D-Manp ->3)-D-Manp	1.1	1.0000	628	76.504	25.055	125.953	23.759

#alpha-D-Manp ->6)-D-Manp	TotalConformers=2 PossibleConformers=2 PossibleCombos=1.1.2
alpha-D-Manp ->6)-D-Manp	1.1.2	0.5599	299	71.152	24.784	173.040	33.093	296.683	11.407
alpha-D-Manp ->6)-D-Manp	1.1.1	0.4401	235	75.813	38.077	161.378	38.128	61.404	27.884

#beta-D-Galp ->4)-D-Glcp	TotalConformers=1 PossibleConformers=1 PossibleCombos=1.1
beta-D-Galp ->4)-D-Glcp	1.1	1.0000	254	284.323	13.943	241.317	20.757

#beta-D-Galp ->4)-D-GlcpNAc	TotalConformers=2 PossibleConformers=2 PossibleCombos=1.2
beta-D-Galp ->4)-D-GlcpNAc	1.2	0.9864	218	286.858	19.942	245.149	16.571
beta-D-Galp ->4)-D-GlcpNAc	1.1	0.0136	3	295.361	11.097	83.668	3.895

#alpha-D-Manp ->4)-D-GlcpNAc	TotalConformers=1 PossibleConformers=1 PossibleCombos=1.1
alpha-D-Manp ->4)-D-GlcpNAc	1.1	1.0000	54	107.637	61.690	207.178	54.549

#beta-D-Xylp ->4)-D-Xylp	TotalConformers=1 PossibleConformers=1 PossibleCombos=1.1
beta-D-Xylp ->4)-D-Xylp	1.1	1.0000	211	277.211	20.446	262.934	20.425

#alpha-L-IdopA2SO3 ->4)-D-GlcpNSO36SO3	TotalConformers=2 PossibleConformers=2 PossibleCombos=1.2
alpha-L-IdopA2SO3 ->4)-D-GlcpNSO36SO3	1.2	0.8947	17	281.096	14.802	242.062	22.656
alpha-L-IdopA2SO3 ->4)-D-GlcpNSO36SO3	1.1	0.1053	2	238.300	11.539	62.450	1.050

#alpha-D-GlcpNSO36SO3 ->4)-L-IdopA2SO3	TotalConformers=1 PossibleConformers=1 PossibleCombos=1.1
alpha-D-GlcpNSO36SO3 ->4)-L-IdopA2SO3	1.1	1.0000	19	80.017	20.326	219.410	18.442

#alpha-D-Manp ->2)-D-Manp	TotalConformers=1 PossibleConformers=1 PossibleCombos=1.1
alpha-D-Manp ->2)-D-Manp	1.1	1.0000	317	77.598	17.631	137.555	33.021

#alpha-D-Manp THR	TotalConformers=1 PossibleConformers=1 PossibleCombos=1.1
alpha-D-Manp THR	1.1	1.0000	139	86.797	24.349	127.166	22.706

#alpha-D-Galp ->4)-D-Galp	TotalConformers=3 PossibleConformers=3 PossibleCombos=1.3
alpha-D-Galp ->4)-D-Galp	1.1	0.9200	23	70.630	3.594	102.757	2.364
#alpha-D-Galp ->4)-D-Galp	1.2	0.0400	1	80.100	-0.000	115.600	-0.000
#alpha-D-Galp ->4)-D-Galp	1.3	0.0400	1	43.000	-0.000	134.800	-0.000

#alpha-D-6-deoxy-Glcp4N ->4)-D-Glcp	TotalConformers=2 PossibleConformers=2 PossibleCombos=1.2
alpha-D-6-deoxy-Glcp4N ->4)-D-Glcp	1.2	0.9412	32	103.565	19.020	232.624	21.033
alpha-D-6-deoxy-Glcp4N ->4)-D-Glcp	1.1	0.0588	2	92.650	1.250	54.650	5.254

#alpha-L-Fucp ->6)-D-GlcpNAc	TotalConformers=2 PossibleConformers=2 PossibleCombos=1.1.2
alpha-L-Fucp ->6)-D-GlcpNAc	1.1.2	0.8187	149	287.067	23.079	180.443	35.526	297.074	18.501
alpha-L-Fucp ->6)-D-GlcpNAc	1.1.1	0.1813	33	287.514	45.620	137.790	77.920	65.935	21.670

#alpha-D-Glcp ->1)-D-Glcp	TotalConformers=4 PossibleConformers=4 PossibleCombos=4.1
alpha-D-Glcp ->1)-D-Glcp	1.1	0.9531	61	71.658	11.148	188.233	8.569
#alpha-D-Glcp ->1)-D-Glcp	2.1	0.0156	1	136.900	-0.000	158.000	-0.000
#alpha-D-Glcp ->1)-D-Glcp	4.1	0.0156	1	314.300	-0.000	218.300	-0.000
#alpha-D-Glcp ->1)-D-Glcp	3.1	0.0156	1	183.400	-0.000	189.000	-0.000

#beta-D-Manp ->6)-D-Manp	TotalConformers=8 PossibleConformers=8 PossibleCombos=2.2.2
beta-D-Manp ->6)-D-Manp	1.1.2	0.3514	13	214.213	24.614	191.704	16.640	267.390	25.161
beta-D-Manp ->6)-D-Manp	2.2.2	0.1622	6	309.382	42.635	251.652	12.042	310.048	24.285
beta-D-Manp ->6)-D-Manp	2.1.1	0.1622	6	340.907	11.588	130.741	8.759	150.815	8.514
beta-D-Manp ->6)-D-Manp	1.1.1	0.0811	3	192.189	28.692	88.362	41.291	102.188	36.949
beta-D-Manp ->6)-D-Manp	1.2.1	0.0811	3	249.759	14.121	280.549	8.892	158.193	31.880
beta-D-Manp ->6)-D-Manp	2.2.1	0.0811	3	301.372	5.288	257.168	3.176	92.811	13.586
beta-D-Manp ->6)-D-Manp	2.1.2	0.0541	2	286.250	13.006	169.000	12.651	222.450	15.138
#beta-D-Manp ->6)-D-Manp	1.2.2	0.0270	1	224.200	-0.000	248.100	-0.000	309.400	-0.000

#alpha-D-GlcpNAc ->4)-D-GlcpNAc	TotalConformers=2 PossibleConformers=2 PossibleCombos=1.2
alpha-D-GlcpNAc ->4)-D-GlcpNAc	1.2	0.7812	25	44.718	41.277	217.903	25.668
alpha-D-GlcpNAc ->4)-D-GlcpNAc	1.1	0.2188	7	79.607	7.448	84.848	15.435

#alpha-D-Glcp2fluoro GLU	TotalConformers=2 PossibleConformers=2 PossibleCombos=2.1
alpha-D-Glcp2fluoro GLU	1.1	0.6667	14	87.701	3.102	185.686	7.350
alpha-D-Glcp2fluoro GLU	2.1	0.3333	7	167.884	21.422	190.765	8.277

#beta-D-Glcp ->4)-D-Glcp2fluoro	TotalConformers=1 PossibleConformers=310 PossibleCombos=310.1
beta-D-Glcp ->4)-D-Glcp2fluoro	235.1	1.0000	10	285.650	0.768	252.663	7.919

#beta-D-Glcp ->4)-D-Glcp	TotalConformers=2 PossibleConformers=2 PossibleCombos=1.2
beta-D-Glcp ->4)-D-Glcp	1.2	0.9902	507	280.661	23.055	230.724	18.432
beta-D-Glcp ->4)-D-Glcp	1.1	0.0098	5	252.894	61.367	79.168	20.225

#alpha-D-Glcp ->2)-D-Fruf	TotalConformers=4 PossibleConformers=6 PossibleCombos=2.3
alpha-D-Glcp ->2)-D-Fruf	1.2	0.9594	189	99.959	13.909	198.920	14.759
alpha-D-Glcp ->2)-D-Fruf	1.1	0.0203	4	96.432	6.925	91.902	15.954
alpha-D-Glcp ->2)-D-Fruf	1.3	0.0152	3	113.315	19.051	339.695	20.457
#alpha-D-Glcp ->2)-D-Fruf	2.2	0.0051	1	209.800	-0.000	254.000	-0.000

#beta-L-Fucp ->6)-D-GlcpNAc	TotalConformers=2 PossibleConformers=2 PossibleCombos=1.2.1
beta-L-Fucp ->6)-D-GlcpNAc	1.2.1	0.6154	16	98.892	33.629	225.941	37.115	332.498	25.843
beta-L-Fucp ->6)-D-GlcpNAc	1.1.1	0.3846	10	103.414	40.246	92.141	24.712	23.310	85.871

#alpha-D-Neup5Ac ->3)-D-Galp	TotalConformers=3 PossibleConformers=4 PossibleCombos=2.2
alpha-D-Neup5Ac ->3)-D-Galp	1.1	0.7344	94	63.046	11.339	106.937	14.447
alpha-D-Neup5Ac ->3)-D-Galp	2.1	0.2500	32	308.594	7.821	100.459	3.960
alpha-D-Neup5Ac ->3)-D-Galp	1.2	0.0156	2	76.850	15.545	275.000	11.640

#alpha-D-GlcpNAc ASN	TotalConformers=3 PossibleConformers=3 PossibleCombos=1.1.1.3
alpha-D-GlcpNAc ASN	1.1.1.2	0.5122	21	61.937	69.342	176.813	73.633	171.955	52.020	192.487	13.850
alpha-D-GlcpNAc ASN	1.1.1.3	0.3171	13	79.559	52.157	165.220	71.657	152.540	40.353	284.164	11.909
alpha-D-GlcpNAc ASN	1.1.1.1	0.1707	7	75.966	31.371	149.241	87.898	171.980	70.823	62.977	12.932

#alpha-L-Fucp ->2)-D-Galp	TotalConformers=2 PossibleConformers=2 PossibleCombos=1.2
alpha-L-Fucp ->2)-D-Galp	1.2	0.7463	153	290.958	9.135	259.727	9.177
alpha-L-Fucp ->2)-D-Galp	1.1	0.2537	52	252.769	24.576	217.982	8.745

#beta-D-Glcp ->3)-D-Glcp	TotalConformers=4 PossibleConformers=4 PossibleCombos=1.4
beta-D-Glcp ->3)-D-Glcp	1.3	0.9579	91	281.183	16.328	120.318	18.697
beta-D-Glcp ->3)-D-Glcp	1.2	0.0211	2	279.700	10.126	63.350	2.751
#beta-D-Glcp ->3)-D-Glcp	1.4	0.0105	1	258.200	-0.000	291.000	-0.000
#beta-D-Glcp ->3)-D-Glcp	1.1	0.0105	1	244.400	-0.000	11.300	-0.000

#beta-D-Manp ->4)-D-Manp	TotalConformers=4 PossibleConformers=4 PossibleCombos=2.2
beta-D-Manp ->4)-D-Manp	2.2	0.9176	78	279.236	18.947	229.255	29.133
beta-D-Manp ->4)-D-Manp	2.1	0.0471	4	216.486	39.579	55.058	21.767
beta-D-Manp ->4)-D-Manp	1.1	0.0235	2	62.100	3.801	68.600	11.135
#beta-D-Manp ->4)-D-Manp	1.2	0.0118	1	43.100	-0.000	113.100	-0.000

#alpha-D-GalpA ->4)-D-GalpA	TotalConformers=1 PossibleConformers=1 PossibleCombos=1.1
alpha-D-GalpA ->4)-D-GalpA	1.1	1.0000	59	87.304	27.798	127.757	30.679

#beta-D-Manp ->3)-D-Manp	TotalConformers=1 PossibleConformers=1 PossibleCombos=1.1
beta-D-Manp ->3)-D-Manp	1.1	1.0000	21	283.585	59.065	155.652	52.897

#beta-D-GlcpNAc ->2)-D-Manp	TotalConformers=6 PossibleConformers=10 PossibleCombos=2.5
beta-D-GlcpNAc ->2)-D-Manp	2.3	0.8416	85	279.370	12.638	152.356	10.516
beta-D-GlcpNAc ->2)-D-Manp	2.2	0.1089	11	272.658	11.632	112.072	3.013
beta-D-GlcpNAc ->2)-D-Manp	1.3	0.0198	2	94.100	18.254	142.850	7.159
#beta-D-GlcpNAc ->2)-D-Manp	2.1	0.0099	1	185.000	-0.000	41.500	-0.000
#beta-D-GlcpNAc ->2)-D-Manp	2.5	0.0099	1	267.700	-0.000	302.400	-0.000
#beta-D-GlcpNAc ->2)-D-Manp	2.4	0.0099	1	245.600	-0.000	252.800	-0.000

#beta-D-GalpNAc ->3)-D-Galp	TotalConformers=4 PossibleConformers=6 PossibleCombos=2.3
beta-D-GalpNAc ->3)-D-Galp	2.2	0.5882	10	326.010	4.351	115.691	6.601
beta-D-GalpNAc ->3)-D-Galp	1.1	0.1765	3	272.567	7.357	63.440	12.910
beta-D-GalpNAc ->3)-D-Galp	1.2	0.1765	3	274.823	6.571	104.999	3.943
#beta-D-GalpNAc ->3)-D-Galp	1.3	0.0588	1	273.200	-0.000	270.400	-0.000

#alpha-D-GalpNAc THR	TotalConformers=1 PossibleConformers=1 PossibleCombos=1.1
alpha-D-GalpNAc THR	1.1	1.0000	12	74.484	6.354	126.217	15.617

#beta-D-Galp ->3)-D-GalpNAc	TotalConformers=1 PossibleConformers=1 PossibleCombos=1.1
beta-D-Galp ->3)-D-GalpNAc	1.1	1.0000	77	281.049	11.563	100.001	16.569

#alpha-D-Galp ->2)-D-Manp	TotalConformers=2 PossibleConformers=2 PossibleCombos=1.2
alpha-D-Galp ->2)-D-Manp	1.2	0.9000	9	83.333	5.913	149.531	5.722
#alpha-D-Galp ->2)-D-Manp	1.1	0.1000	1	91.900	-0.000	95.400	-0.000

#alpha-D-Manp ->4)-D-Manp	TotalConformers=6 PossibleConformers=6 PossibleCombos=2.3
alpha-D-Manp ->4)-D-Manp	1.2	0.4737	9	94.047	36.669	241.081	15.655
alpha-D-Manp ->4)-D-Manp	1.1	0.1579	3	85.024	33.107	99.095	17.796
alpha-D-Manp ->4)-D-Manp	2.1	0.1579	3	295.868	34.639	128.523	52.715
alpha-D-Manp ->4)-D-Manp	1.3	0.1053	2	146.050	5.454	341.400	7.712
#alpha-D-Manp ->4)-D-Manp	2.3	0.0526	1	230.000	-0.000	355.300	-0.000
#alpha-D-Manp ->4)-D-Manp	2.2	0.0526	1	308.200	-0.000	251.500	-0.000

#alpha-D-Glcp ->6)-D-Glcp	TotalConformers=4 PossibleConformers=4 PossibleCombos=1.1.4
alpha-D-Glcp ->6)-D-Glcp	1.1.4	0.5172	15	58.210	17.971	176.663	18.380	299.005	12.765
alpha-D-Glcp ->6)-D-Glcp	1.1.1	0.3793	11	69.743	14.622	179.788	30.168	76.268	15.476
alpha-D-Glcp ->6)-D-Glcp	1.1.3	0.0690	2	37.050	23.999	124.150	16.974	234.000	3.101
#alpha-D-Glcp ->6)-D-Glcp	1.1.2	0.0345	1	139.300	-0.000	145.000	-0.000	168.400	-0.000

#alpha-D-Galp ->6)-D-Galp	TotalConformers=4 PossibleConformers=6 PossibleCombos=3.1.2
alpha-D-Galp ->6)-D-Galp	1.1.1	0.4000	10	73.899	3.987	182.341	21.305	180.111	9.406
alpha-D-Galp ->6)-D-Galp	2.1.1	0.3200	8	101.387	1.670	198.385	97.500	169.360	18.160
alpha-D-Galp ->6)-D-Galp	1.1.2	0.1600	4	63.075	0.965	166.750	3.167	300.450	1.736
alpha-D-Galp ->6)-D-Galp	3.1.1	0.1200	3	149.064	11.873	178.042	15.786	181.608	6.088

#alpha-D-Kdop ->4)-D-Kdop	TotalConformers=1 PossibleConformers=1 PossibleCombos=1.1
alpha-D-Kdop ->4)-D-Kdop	1.1	1.0000	14	54.221	12.470	85.776	36.387

#beta-D-Galp ->3)-D-GlcpNAc	TotalConformers=2 PossibleConformers=2 PossibleCombos=2.1
beta-D-Galp ->3)-D-GlcpNAc	2.1	0.9516	59	289.552	12.890	127.382	17.414
beta-D-Galp ->3)-D-GlcpNAc	1.1	0.0484	3	39.931	4.049	124.801	3.103

#beta-D-GlcpNAc ->3)-D-Galp	TotalConformers=2 PossibleConformers=2 PossibleCombos=2.1
beta-D-GlcpNAc ->3)-D-Galp	2.1	0.5172	45	270.374	5.787	94.245	11.378
beta-D-GlcpNAc ->3)-D-Galp	1.1	0.4828	42	286.822	14.510	103.200	18.593

#alpha-D-GalpNAc ->3)-D-Galp	TotalConformers=1 PossibleConformers=1 PossibleCombos=1.1
alpha-D-GalpNAc ->3)-D-Galp	1.1	1.0000	56	59.987	13.358	65.129	14.510

#alpha-D-Xylp ->6)-D-Glcp	TotalConformers=5 PossibleConformers=6 PossibleCombos=1.2.3
alpha-D-Xylp ->6)-D-Glcp	1.2.1	0.7045	31	71.600	19.416	178.698	20.112	65.784	13.454
alpha-D-Xylp ->6)-D-Glcp	1.1.1	0.0909	4	84.224	15.833	86.422	3.959	56.407	8.977
alpha-D-Xylp ->6)-D-Glcp	1.1.3	0.0909	4	65.216	10.824	82.194	10.167	285.009	9.772
alpha-D-Xylp ->6)-D-Glcp	1.2.3	0.0909	4	65.738	9.882	165.376	15.781	296.475	2.767
#alpha-D-Xylp ->6)-D-Glcp	1.2.2	0.0227	1	113.600	-0.000	204.000	-0.000	187.500	-0.000

#beta-D-Galp ->2)-D-Xylp	TotalConformers=2 PossibleConformers=2 PossibleCombos=1.2
beta-D-Galp ->2)-D-Xylp	1.2	0.8235	14	288.268	10.716	220.495	18.164
beta-D-Galp ->2)-D-Xylp	1.1	0.1765	3	271.065	4.016	91.406	6.402

#alpha-D-Neup5Ac ->6)-D-Galp	TotalConformers=4 PossibleConformers=12 PossibleCombos=3.2.2
alpha-D-Neup5Ac ->6)-D-Galp	1.2.1	0.5000	10	67.001	10.001	198.226	13.909	62.239	6.524
alpha-D-Neup5Ac ->6)-D-Galp	1.1.1	0.3000	6	60.800	1.739	104.299	3.675	44.349	2.180
alpha-D-Neup5Ac ->6)-D-Galp	2.2.1	0.1500	3	111.976	8.704	219.293	10.601	40.294	5.870
#alpha-D-Neup5Ac ->6)-D-Galp	3.2.2	0.0500	1	215.300	-0.000	235.600	-0.000	307.600	-0.000

#beta-L-Fucp ->3)-D-GlcpNAc	TotalConformers=1 PossibleConformers=1 PossibleCombos=1.1
beta-L-Fucp ->3)-D-GlcpNAc	1.1	1.0000	20	46.203	52.363	102.147	24.897

#alpha-D-Galp ->6)-D-Glcp	TotalConformers=2 PossibleConformers=2 PossibleCombos=1.1.2
alpha-D-Galp ->6)-D-Glcp	1.1.2	0.5333	8	41.857	20.239	193.732	10.566	298.436	4.682
alpha-D-Galp ->6)-D-Glcp	1.1.1	0.4667	7	68.044	3.428	150.400	29.665	62.461	12.762

#alpha-L-Fucp ->3)-D-Glcp	TotalConformers=1 PossibleConformers=1 PossibleCombos=1.1
alpha-L-Fucp ->3)-D-Glcp	1.1	1.0000	14	291.764	7.582	142.549	2.760

#beta-D-GalpNAc ->4)-D-Galp	TotalConformers=1 PossibleConformers=1 PossibleCombos=1.1
beta-D-GalpNAc ->4)-D-Galp	1.1	1.0000	30	276.791	6.129	129.329	6.504

#alpha-D-Galp ->3)-D-Galp	TotalConformers=6 PossibleConformers=10 PossibleCombos=5.2
alpha-D-Galp ->3)-D-Galp	2.1	0.5397	34	56.003	9.108	70.615	11.788
alpha-D-Galp ->3)-D-Galp	3.2	0.3810	24	97.935	3.573	121.982	6.014
alpha-D-Galp ->3)-D-Galp	4.2	0.0317	2	140.050	2.650	140.300	7.611
#alpha-D-Galp ->3)-D-Galp	5.1	0.0159	1	331.600	-0.000	66.500	-0.000
#alpha-D-Galp ->3)-D-Galp	1.1	0.0159	1	9.500	-0.000	95.100	-0.000
#alpha-D-Galp ->3)-D-Galp	3.1	0.0159	1	86.500	-0.000	54.700	-0.000

#alpha-D-Neup5Ac ->8)-D-Neup5Ac	TotalConformers=2 PossibleConformers=2 PossibleCombos=2.1.1
alpha-D-Neup5Ac ->8)-D-Neup5Ac	1.1.1	0.7500	33	66.826	23.375	118.325	13.338	69.104	24.030
alpha-D-Neup5Ac ->8)-D-Neup5Ac	2.1.1	0.2500	11	295.641	16.622	93.703	17.809	76.414	22.874

#alpha-L-Rhap ->2)-L-Rhap	TotalConformers=1 PossibleConformers=1 PossibleCombos=1.1
alpha-L-Rhap ->2)-L-Rhap	1.1	1.0000	29	285.371	12.008	202.197	24.531

#beta-D-GlcpNAc ->2)-L-Rhap	TotalConformers=2 PossibleConformers=2 PossibleCombos=1.2
beta-D-GlcpNAc ->2)-L-Rhap	1.2	0.6190	13	289.194	4.553	254.758	3.767
beta-D-GlcpNAc ->2)-L-Rhap	1.1	0.3810	8	292.750	3.262	221.512	1.831

#alpha-L-Rhap ->3)-D-GlcpNAc	TotalConformers=1 PossibleConformers=1 PossibleCombos=1.1
alpha-L-Rhap ->3)-D-GlcpNAc	1.1	1.0000	25	276.043	7.690	134.487	7.796

#alpha-L-Araf ->5)-L-Araf	TotalConformers=3 PossibleConformers=4 PossibleCombos=2.1.2
alpha-L-Araf ->5)-L-Araf	2.1.2	0.6000	9	297.726	9.013	208.443	46.818	293.294	13.884
alpha-L-Araf ->5)-L-Araf	2.1.1	0.3333	5	288.068	19.117	152.175	18.272	82.099	26.011
#alpha-L-Araf ->5)-L-Araf	1.1.2	0.0667	1	32.700	-0.000	233.200	-0.000	345.500	-0.000

#alpha-L-Fucp ->4)-D-GlcpNAc	TotalConformers=1 PossibleConformers=1 PossibleCombos=1.1
alpha-L-Fucp ->4)-D-GlcpNAc	1.1	1.0000	24	288.208	8.394	259.949	4.767

#alpha-L-Rhap ->3)-L-Rhap	TotalConformers=1 PossibleConformers=1 PossibleCombos=1.1
alpha-L-Rhap ->3)-L-Rhap	1.1	1.0000	22	289.259	6.760	231.035	3.491

#alpha-D-Glcp ->4)-L-Rhap	TotalConformers=2 PossibleConformers=4 PossibleCombos=2.2
alpha-D-Glcp ->4)-L-Rhap	2.2	0.5000	8	145.137	3.749	147.807	4.107
alpha-D-Glcp ->4)-L-Rhap	1.1	0.5000	8	52.575	3.236	95.913	2.656

#beta-D-GlcpN ->4)-D-GlcpN	TotalConformers=1 PossibleConformers=1 PossibleCombos=1.1
beta-D-GlcpN ->4)-D-GlcpN	1.1	1.0000	24	276.002	22.115	223.131	15.553
