## supplemental_data_and_movie for "Growing Glycans in Rosetta: Accurate *de novo* glycan modeling, density fitting, and rational sequon design": README

# List of elements in supplemental:

## create_substituted_JD.py
  - Used to create substituted job description file for rosetta_scripts_jd3.  
  -  See density_build_example.xml, when when run with this script and the pdb_roots_density_sym.txt file produces the density_build_example_substituted.xml file

 ## create_symmetry_mates.py
  - This was used to create symmetry mates with a 12 A radius using the tool in rosetta tools, which was added to the path. 

 ## density_build_example.xml
  - An an example of a main job definition file before being substituted for each glycan

 ## density_build_example_substituted.xml
  - Substituted version

 ## glycan_conformer_table.txt
  - The resultant glycan conformer table used in Rosetta and benchmarked against

 ## glycan_tree_modeler_example.mov
  - Example of the GlycanTreeModeler made during devopment.  This is not the final version with the best settings, but illustrates glycans growing from their roots layer by layer.  In this case, we had a layer size of two and this was not the hybrid protocol that is currently used and described as the final algorithm in the main text.
   
 ## glycan_tree_relax.xml
  - Main script used for benchmarking.  Note that ALL GlyanTreeModeler options that were benchmarked are now the default. 

 ## glycan_tree_relax_dens.xml
  - Main script used for density building. Note that ALL GlycanTreeModeler options besides the refine optiona are now the default. 

 ## glycan_tree_relax_with_solvation.xml
  - Script used for running the GTM with explicit waters

 ## glycan_water_mediated_hbonds.xml
  - Script for calculating water mediated hydrogen bonds

 ## pdb_roots_density_sym.txt
  - Benchmark glycan list used for create_substituted_JD.py

 ## refine_with_map_dih2.xml
  - Script used for refining input structures into the rosetta energy function using density. Note that this is dihedral space as our glycan tree modeler was shown to work better in dihedral space energy function and minimization than cartesian. 
